# A phenome-wide association and Mendelian Randomisation study of polygenic risk for depression in UK Biobank

**DOI:** 10.1101/617969

**Authors:** Xueyi Shen, David M Howard, Mark J Adams, 23andMe Research Team, Major Depressive Disorder Working Group of the Psychiatric Genomics Consortium, Ian J Deary, Heather C Whalley, Andrew M McIntosh

**Affiliations:** Division of Psychiatry, University of Edinburgh, Edinburgh, United Kingdom; Centre for Cognitive Ageing and Cognitive Epidemiology, University of Edinburgh, Edinburgh, United Kingdom; Department of Psychology, University of Edinburgh, Edinburgh, United Kingdom; 23andMe, Inc., Mountain View, CA, USA

## Abstract

Depression is the leading cause of worldwide disability but there remains considerable uncertainty regarding its neural and behavioural associations. Depression is known to be heritable with a polygenic architecture, and results from genome-wide associations studies are providing summary statistics with increasing polygenic signal that can be used to estimate genetic risk scores for prediction in independent samples. This provides a timely opportunity to identify traits that are associated with polygenic risk of depression in the large and consistently phenotyped UK Biobank sample. Using the Psychiatric Genomics Consortium (PGC), 23andMe and non-imaging UK Biobank datasets as reference samples, we estimated polygenic risk scores for depression (depression-PRS) in a discovery sample of 10,674 people and a replication sample of 11,214 people from the UK Biobank Imaging Study, testing for associations with 210 behavioural and 278 neuroimaging phenotypes. In the discovery sample, 93 traits were significantly associated with depression-PRS after multiple testing correction. Among these, 92 traits were in the same direction, and 69 were significant in the replication analysis. For imaging traits that replicated across samples, higher depression-PRS was associated with lower global white matter microstructure, association-fibre and thalamic-radiation microstructural integrity (absolute β: 0.023 to 0.040, P_FDR_: 0.045 to 3.92×10^-4^). Mendelian Randomisation analysis showed a causal effect of liability to depression on these structural brain measures (β: 0.125 to 0.707, p_FDR_<0.048). Replicated behavioural traits that positively associated with depression-PRS included sleep problems, smoking status, measures of pain and stressful life experiences, and those negatively associated with depression-PRS included subjective ratings of physical health (absolute β: 0.014 to 0.180, P_FDR_: 0.046 to 8.54×10^-15^). Effect of depression PRS on mental health in the presence of reported childhood trauma, stressful life events and those living in more socially deprived areas showed increased variance explained by 1.42 – 4.08 times (p_FDR_ for their interaction with depression-PRS: 0.049 to 0.003). Overall, the present study revealed replicable associations between depression-PRS and white matter microstructure that appeared to be a causal consequence of liability to depression. Analyses provided further evidence that greater effects of polygenic risk of depression are found in individuals exposed to risk-conferring environments.

## Introduction

Major Depression is the leading contributor to the overall global burden of disease^1^, mainly due to its high prevalence^2,3^, disabling consequences^2^ and low treatment response^4^. Twin studies have shown that Major Depression is partially heritable (h^2^=37%)^3^. Recent genome-wide association studies (GWAS) have revealed genetic loci that have variants associated with Major Depression, such as the study by Wray *et al.* for the Psychiatric Genomics Consortium (PGC) that identified 44 risk variants^5^, and a more recent study by Howard *et al.* which identified 102 genetic variants^6^. Although each single genetic variant contributes very little to disease liability, the genetic risk scores based on the additive effect of common genetic variants over the whole genome, i.e. polygenic risk scores (PRS), can account for a significant amount of phenotypic variance^7^. The latest phase of GWAS on Major Depression now provide the ability to more precisely estimate polygenic risk of depression in independent samples^6^ and thereby identify traits whose genetic architecture is shared with Major Depression.

Major Depression is phenotypically correlated with many traits including several behavioural measures, brain structure and function, specific cognitive functions, and several physical conditions^8–13^. It is important to investigate the associations between the genetic predisposition to Major Depression and a wide range of phenotypes, to help identify causal risk factors, risk-conferring mechanisms and the causal consequences of Major Depression^14^. Until recently, however, this approach has received relatively little attention owing to a lack of data resources with the appropriate coverage of genetic, behavioural and neuroimaging traits to test for these polygenic risk associations with sufficient statistical power^15–17^.

In order to tackle the above difficulties, the present study used the largest data sets available to date for both depression-PRS generation and a wide range of phenotypes, including neuroimaging. Depression-PRS were generated using summary statistics from the most recent meta-analysis combining the Psychiatric Genomics Consortium (PGC), UK Biobank, and 23andMe (N=0.8 million)^6^. A phenome-wide association study (PheWAS) approach was used to estimate the strength and significance of associations between depression-PRS and other behavioural, cognitive and neuroimaging traits. PheWAS was conducted on the latest neuroimaging data releases from the UK Biobank imaging project^18^ that included a discovery sample of 10,674 people, and a replication sample of 11,214 people (21,888 individuals in total), the largest dataset to date that contains both genetic and cross-modality neuroimaging data. Where depression-PRS were associated with neuroimaging phenotypes, we additionally tested whether this was a causal consequence of depression, or conversely, whether neuroimaging measures had a causal effect on depression, using Mendelian Randomisation and structural equational modelling. We also tested for the presence of gene-by-environment interactions using measures of early-life risk factors and sociodemographic variables available in UK Biobank^19,20^.

## Methods

### Participants

Data from 21,888 individuals who participated in the UK Biobank imaging study^18^ were included in the current study (released in two waves, in May and October 2018). The discovery sample included participants mainly from the first data release, and the replication sample from the second release (details for the discovery and replication samples can be found in supplementary materials and Figure S1). The majority of participants were assessed in the Cheadle MRI site (80.1%) and the rest in the Newcastle site (19.9%). Comparisons between the sites are reported in the supplementary materials. All imaging data was collected using a 3T Siemens Skyra (software platform VD13) machine.

Behavioural and neuroimaging data acquisition were conducted under standard protocols^18,21^. Written consent was acquired for all participants. Data acquisition and analyses in the present study were conducted under UK Biobank Application #4844. Ethical approval was accepted by the National Health Service (NHS) Research Ethics Service (11/NW/0382).

### Depression-PRS

In the present study, the sample used for generating GWAS summary statistics is referred to as the training dataset. The samples in which depression-PRS were generated and tested are referred to as the testing samples, which include both discovery and replication samples (as described above). We removed any overlapping individuals from the training sample (used to estimate allele effects for polygenic profiling) and testing datasets (where the effects of PRS scores were estimated) (see supplementary methods).

Polygenic risk scores were calculated using the summary statistics from a meta-analysis of depression genome-wide association study (GWAS) from three cohorts, including PGC analysis of major depression^5^, the 23andMe discovery sample in the Hyde et al. analysis of self-reported clinical depression^22^, and a broad depression phenotype from UK Biobank within individuals who had not participated in the imaging study^23^. This meta-analysis provided a total training dataset of 785,581 individuals (238,360 cases and 547,221 controls; for further details see the study by Howard *et al*.^6^). In order to utilise genetic variants that replicated across all 3 datasets, we used the summary statistics that included only the 8,099,819 single nucleotide polymorphisms that were present in the GWAS data from all three cohorts^6^.

PRSice 2.0 (used with PLINK 1.9)^24^ was used to calculate the depression-PRS. Before the analyses were conducted, individuals who met the following criterion were removed from the testing dataset: related or non-European-ancestry individuals and those that were included in PGC, 23andMe and UK Biobank GWAS on depression (details can be found in Supplementary materials). All sample sizes reported below are the numbers after these data removal steps. Genotyping and quality control were conducted by UK Biobank as described in an earlier protocol paper^25^. Details of SNP quality control and imputation can be found in the supplementary materials. Eight p-value thresholds were applied to select genetic variants included in calculating polygenic risk scores, as p<0.0005, p<0.001, p<0.005, p<0.01, p<0.05, p<0.1, p<0.5 and p<1.

### Behavioural phenotypes

The behavioural phenotypes consisted of six broad categories, containing 210 variables in total. Where summary data were available (e.g. neuroticism total score), the individual items used to derive the summary data were not included. Phenotypes that were available on fewer than 2,000 people in the discovery sample were also excluded from further analysis. Mean sample sizes for all traits contained in each category are included in brackets below. For further details see in Tables 1 and S1. Categories included: (1) Mental health (N_discovery_=7,910 and N_replication_=3,845), including self-reported symptoms of major psychiatric conditions^26^. In this category, three definitions for depression were included: broad depression, which was a self-declared definition of whether the participant had seen a psychiatrist for nerves, anxiety, tension or depression^6,23^, probable depression which was derived from an abbreviated set of self-declared symptoms of major depression and hospital admission history^27^, and CIDI depression, a measure assessing full diagnostic criteria for depression based on questions from a shortened version of the structured Composite International Diagnostic Interview^26^. (2) Sociodemographic measures (N_discovery_=8,759 and N_replication_=4,352), such as household income and educational attainment. (3) Early-life risk factors (N_discovery_=9,775 and N_replication_=10,872), containing physical measures such as birth weight, and environmental variables like adoption and maternal smoking. (4) Lifestyle measures (N_discovery_=9.232 and N_replication_=4,4796), which mainly included items on sleep, smoking, alcohol consumption and diet, (5) Physical measures (N_discovery_=8,961 and N_replication_=4,618), consisting of self-declared medical conditions such as recent pains, cancers, operations, heart and artery diseases and other major illnesses, and also measures of blood pressure, arterial stiffness and hand-grip strength, and finally (6) Cognition (N_discovery_=8,153 and N_replication_=4,105). This included four tests conducted at the assessment centres, four tests conducted online and a general measure derived based on the tests conducted at the assessment centres which have larger sample sizes^28^ (see more details in the supplementary methods).

All of the behavioural phenotypes, with the exception of mental health items derived from online-follow up questionnaires (see Table 1), were primarily acquired at the same time as the imaging assessment. Missing data for the imaging assessment were imputed using data available from the baseline assessment. The mean age difference between imaging assessment and the initial visit was 8.53 years (SD=1.56 years). Sample sizes and descriptions for all the behavioural phenotypes can be found in Table S1.

**Table 1.** A summary of phenotypes. A total of 213 behavioural phenotypes (six categories) and 278 neuroimaging variables (four categories) are included.

| Category | General description | Number of traits | Discovery Sample |  | Replication sample |  | UK Biobank data modality | Covariates |
| --- | --- | --- | --- | --- | --- | --- | --- | --- |
|  |  |  | Sample sizes | Mean sample size | Sample sizes | Mean sample size |  |  |
| Mental health | Mental health questionnaires from Touchscreen's mental-health section and questions from the online follow-up section<sup>26</sup> were included. | 45 | 3,299 - 10,674 | 7,910 | 1,519 - 5,565 | 3,845 | Touchscreen, Online follow-up | Age, age<sup>2</sup>, sex, genotyping array, 15 genetic principal components (PCs) |
| Sociodemographic | Items include education, household income, ethnicity, immigration status and social deprivation. | 5 | 8,054 - 9,941 | 8,759 | 3,824 - 5,199 | 4,352 | Touchscreen |  |
| Early-life risk factor | Self-declared early-life risk factors. Mainly derived based on another study by Ruth et al<sup>58</sup>. Items include developmental factors such as birth weight and comparative weight and height at early ages. Ages of parental death were additionally included as parental factors. | 11 | 7,484 - 10,674 | 9,775 | 8,727 - 11,730 | 10,872 | Touchscreen, Online follow-up |  |
| Lifestyle measures | Self-declared lifestyle questions, mainly including items relevant to sleep quality, smoking, alcohol consumption, usage of electronic devices, food and beverage intake, appearance, and social activities. | 69 | 2,789 - 10,674 | 9,294 | 1,392 - 5,565 | 4,829 | Touchscreen |  |
| Physical measures | This category contains data from self-declared physical conditions from the Touchscreen data modality and measured physical data from the Physical Measures modality. Self-declared items include general physical health, chronic and recent pains, cardiovascular problems, general diabetic and cancer problems and bone fractures. Measured physical data contains general and regional body mass/fat index, impedance and hand grip strength. A significant amount of phenotypes under this category were excluded from analysis due to insufficient number of participants included (N<2,000). | 67 | 2,155 - 10,674 | 8,961 | 1,047 - 5,565 | 4,618 | Touchscreen |  |
| Cognition | Tasks were selected for having acceptable reliability based on peer-reviewed publications<sup>59-61</sup>. These include four tasks conducted at the assessment centres and four tasks completed online. A variable of g score derived from the tasks completed at the assessment centres was added. | 16 | 5,250 - 10,674 | 8,153 | 2,586 - 5,565 | 4,105 | Touchscreen, Online follow-up |  |
| Intracranial/subcortical volume | Measures were derived from T1 data. Eight subcortical regions were mapped and measured. A derived measure of the intracranial volume was generated by adding up the volumes for grey and white matter and ventricular cerebrospinal fluid. | 9 | 10,663 | -- | 5,561 | -- | Brain imaging | Age, age<sup>2</sup>, sex, genotyping array, 15 PCs, ICV, scanner position on the x,y and z axes |
| White matter microstructure | Weighted-mean fractional anisotropy and mean diffusivity of major tracts were derived for 27 major tracts (12 bilateral and 3 unilateral), mapped using probabilistic tractography. In addition to individual tracts, general variance in all tracts (gTotal) and variance in association/commissural fibres (gAF), thalamic radiations (gTR) and projection fibres (gPF) were derived using principal component analysis. | 38 | FA: 9,374; MD: 9,350 | -- | FA: 5,211; MD: 5,203 | -- | Brain imaging |  |
| Resting-state functional connectivity | Two-two paired, partial correlation matrix of 21 parcellated nodes generated by group-ICA was estimated and used as a measure for functional connectivity. | 210 | 9782 | -- | 5269 | -- | Brain imaging | Age, age<sup>2</sup>, sex, genotyping array, 15 PCs, mean motion, scanner position on the x,y and z axes |
| Resting-state fluctuation amplitude | Fluctuation amplitude of low-frequency signal in the 21 parcellated nodes. | 21 | 9782 | -- | 5269 | -- | Brain imaging | Age, age<sup>2</sup>, sex, genotyping array, 15 PCs, mean motion, scanner position on the x,y and z axes |

### Neuroimaging phenotypes

Neuroimaging data consisted of: (1) intracranial and subcortical volumes (N_discovery_=10,663 and N_replication_=5,561), containing eight major structures^29^; (2) white matter microstructure, indexed by fractional anisotropy (FA, N_discovery_=9,374 and N_replication_=5,211) and mean diffusivity (MD, N_discovery_=9.350 and N_replication_=5,203) for measures of white matter microstructure, in which we included three measures of association, projection and thalamic radiation subsets, and 15 major individual white matter tracts^29^; (3) pair-wise resting-state (rsfMRI) functional connectivity (N_discovery_=9,782 and N_replication_=5,269) of 21 nodes over the whole brain^30^; and finally (4) the amplitude of low-frequency rsfMRI signal fluctuation of the 21 nodes (N_discovery_=9,782 and N_replication_=5,269). All four types of neuroimaging data consisted of the imaging-derived phenotypes (IDPs) provided by UK Biobank (see Figures S2-4). Images were acquired, pre-processed and quality controlled by UK Biobank using FMRIB Software Library (FSL) packages by a standard protocol (URL: https://biobank.ctsu.ox.ac.uk/crystal/docs/brain_mri.pdf), which was also described in two protocol papers^18,31^. All pilot study data with inconsistent scanner settings and data that did not pass the initial quality assessment conducted by UK Biobank imaging team were not included in the analysis. All imaging data was collected using a 3T Siemens Skyra (software platform VD13) machine. For clarity, major steps of pre-processing were described in the supplementary materials.

### Statistic models for PheWAS

The GLM function in R was used to test the PheWAS associations^32^, and the LME function from the ‘nlme’ package in R^33^ was used to test bilateral brain structures where hemisphere was included as a within-subject variable. Depression-PRSs were set as independent fixed-effects, and behavioural and neuroimaging phenotypes were set as dependent variables. Overall, 488 phenotypes (210 behavioural phenotypes + 9 intracranial/subcortical volumes + 38 white matter microstructural measures + 210 rsfMRI connectivity + 21 rsfMRI fluctuation amplitude) * 8 depression-PRS (under 8 p thresholds) = 3,904 tests across phenotypes and depression-PRS p thresholds were corrected altogether by FDR-correction^34^ using p.adjust function in R (q<0.05).

Covariates included in all association tests were sex, age, age^2^, the first 15 genetic principal components and genotyping array^23^. For the replication analysis, MRI site was added in addition to the above covariates for all association tests. In addition to these covariates, adjustments were made for other confounders that were relevant to each phenotypic category, as listed below. Scanner positions on the x, y and z axes were included in the models for all brain phenotypes to control for static-field heterogeneity^35^. Mean head motion was set as a covariate for the rsfMRI data^30,36^. Subcortical volumetric tests controlled for intracranial volume^29,37^. Hemisphere was controlled for where applicable in bilateral brain structural phenotypes^29^. A list of covariates for each type of phenotype can be found in Table 1.

In order to help compare the results of logistic and linear regression models, we report the standardised regression coefficients for the models as effect sizes (β) for both types of models. Log-transformed odds ratio for binary dependent variables using logistic regression models are therefore reported. FDR-corrected p values are reported throughout. When effect sizes of different signs were presented together, we reported the range of absolute effect sizes.

### Replication analysis for PheWAS

Traits that were found to be significantly associated with depression-PRS at a minimum of four GWAS association p-thresholds were selected for re-analysis in the independent replication sample. The replication analysis was conducted on the selected traits across all eight depression-PRS thresholds. Results were considered to be replicated where they showed an identical direction of effect across discovery and replication samples, and where the p value for the replication sample analysis was significant after correction for multiple testing for depression-PRS at a minimum of four p-thresholds. FDR correction was applied to all the tests conducted in the replication analysis across all traits and GWAS p thresholds (e.g., if m traits were taken into replication analysis, then p-value adjustment was applied to all m*8 thresholds).

### Bidirectional Mendelian Randomisation analyses on depression and neuroimaging variables

We used the ‘twosampleMR’ package in R to conduct bidirectional Mendelian Randomisation analyses between depression and neuroimaging variables in order to test for causal effects^38^. Mendelian Randomisation uses genetic data as instruments for testing whether there is any causal effect between an exposure and an outcome variable. A chart illustrating the underlying models can be found in Figure 5, a flow chart of all the steps in Figure S5, and the main procedure is described briefly below.

**Figure 1:**
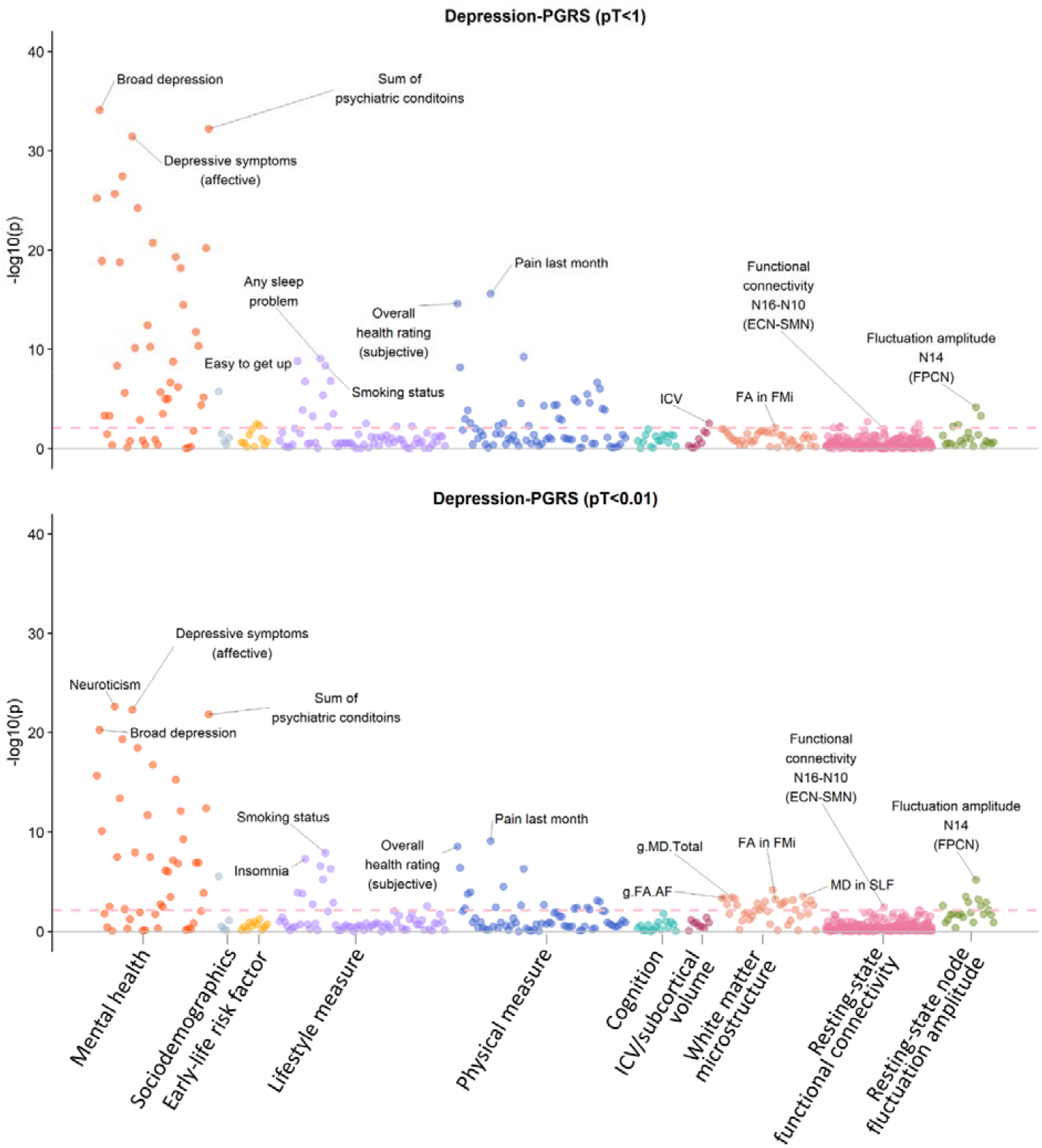
Significance plot for all phenotypes for depression-PRS at p-threshold (pT) < 1 (top figure) and pT < 0.01 (bottom figure), with the x axis showing phenotypes, and the y axis showing the -log_10_ of uncorrected p values. Each dot represents one phenotype, and the colours indicate their according categories. The dashed lines indicate the threshold to survive FDR-correction. FDR-correction was applied over all the traits and all depression-PRS (see Methods). From left to right on the x axis, categories were shown by the sequence of mental health measure, early-life risk factor, sociodemographics, lifestyle measure, physical measure, cognition, intracranial/subcortical volume, white matter microstructure, resting-state functional connectivity and resting-state fluctuation amplitude. Representative top findings are annotated in the figure. For the abbreviations: FPCN=fronto-parietal control network, ECN=executive control network, SMN=sensori-motor network, FA=fractional anisotropy, MD for white matter microstructure=mean diffusivity, AF=association fibres, FMa=forceps major, FMi=forceps minor, SLF=superior longitudinal fasciculus.

**Figure 2.**
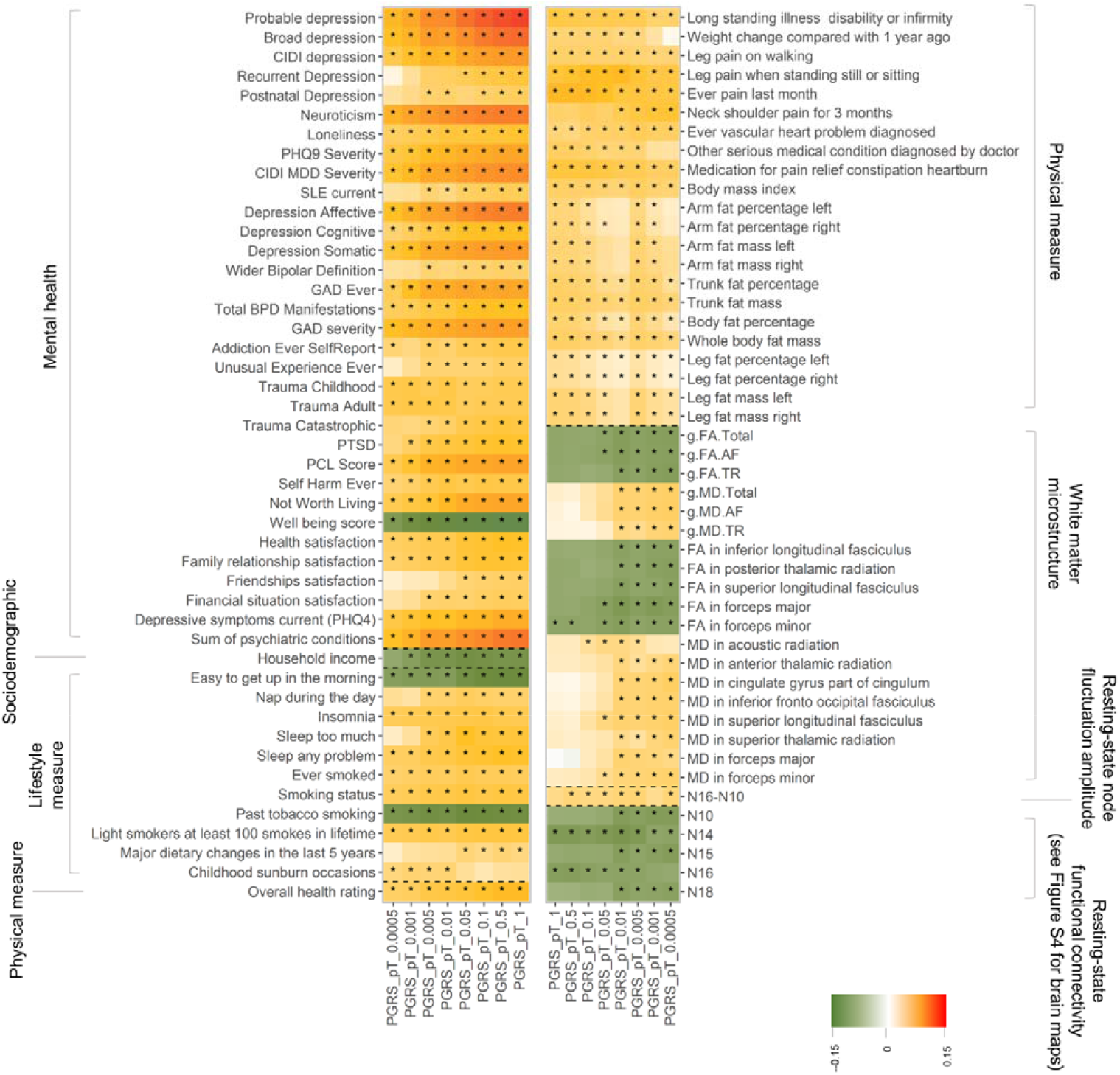
Heatmap for the traits that were significantly associated with depression-PRS at a minimum of four p thresholds. Shades of cells indicate the standardised effect sizes (β). A larger effect size was shown by a darker colour. Cells with an asterisk were significant after FDR-correction. Descriptions for the variables in detail can be found in Table 1 and S1.

**Figure 3.**
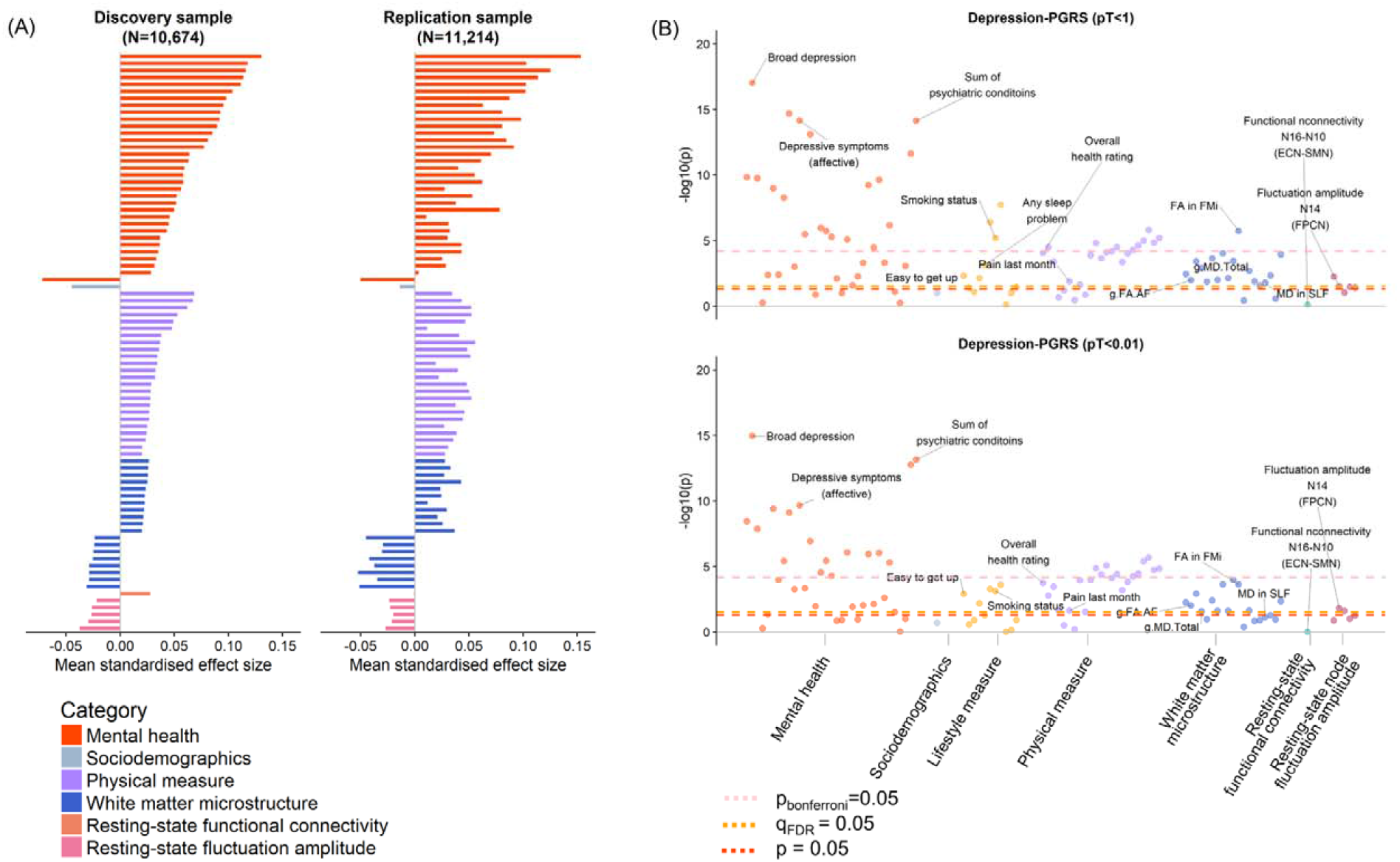
Results for replication analysis. (A) Comparisons of effect sizes for the discovery and replication samples. The x axes represent the mean standard effect size across depression-PRS at all eight p-thresholds (pT). Colours for the bars indicate their categories (from top to bottom: mental health measure, sociodemographics, lifestyle measure, physical measure, white matter microstructure, resting-state functional connectivity and resting-state fluctuation amplitude). (B) Significance plot for the replication analysis on representative depression-PRS at pT<1 and pT<0.01, in accordance with Figure 1. Top hits shown in the discovery sample (Figure 1) are annotated in the figure. Explanations for the abbreviations can be found in the legend of Figure 1.

**Figure 4.**
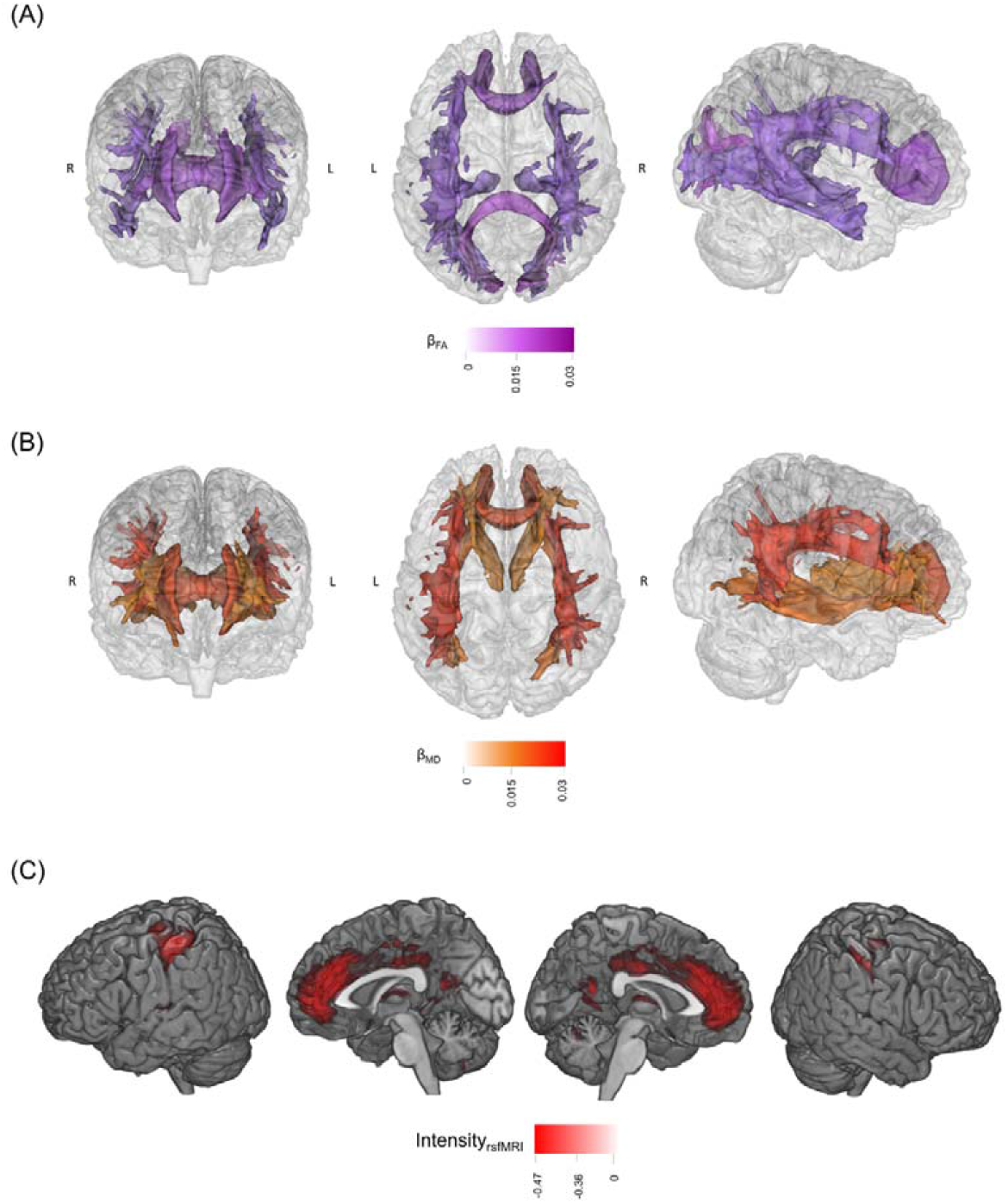
Panels A and B are the brain maps for the significant associations between depression-PRS and white matter microstructure in fractional anisotropy (FA, panel A) and mean diffusivity (MD, panel B) of major tracts. The shade for each tract represents the standardised effect size (β), with a darker shade showing a greater mean β across all depression-PRS at the significant p-thresholds (pT). From left to right are from anterior, superior and right view. For clarity, among the tracts presented in Figure 2, the ones that showed consistent associations across at least four depression-PRS p-thresholds are presented. Panel C shows the brain maps for regions involved in significant associations between resting-state fluctuation amplitude and depression-PRS. Regions that show consistent associations across at a minimum of four depression-PRS p-thresholds are presented. Visualisation of results is achieved by calculating the average intensity of ICA maps, weighted by their mean β across the pT. For clarity, the brain maps shown below have a threshold applied on (intensity over 25% of the highest global intensity).

**Figure 5.**
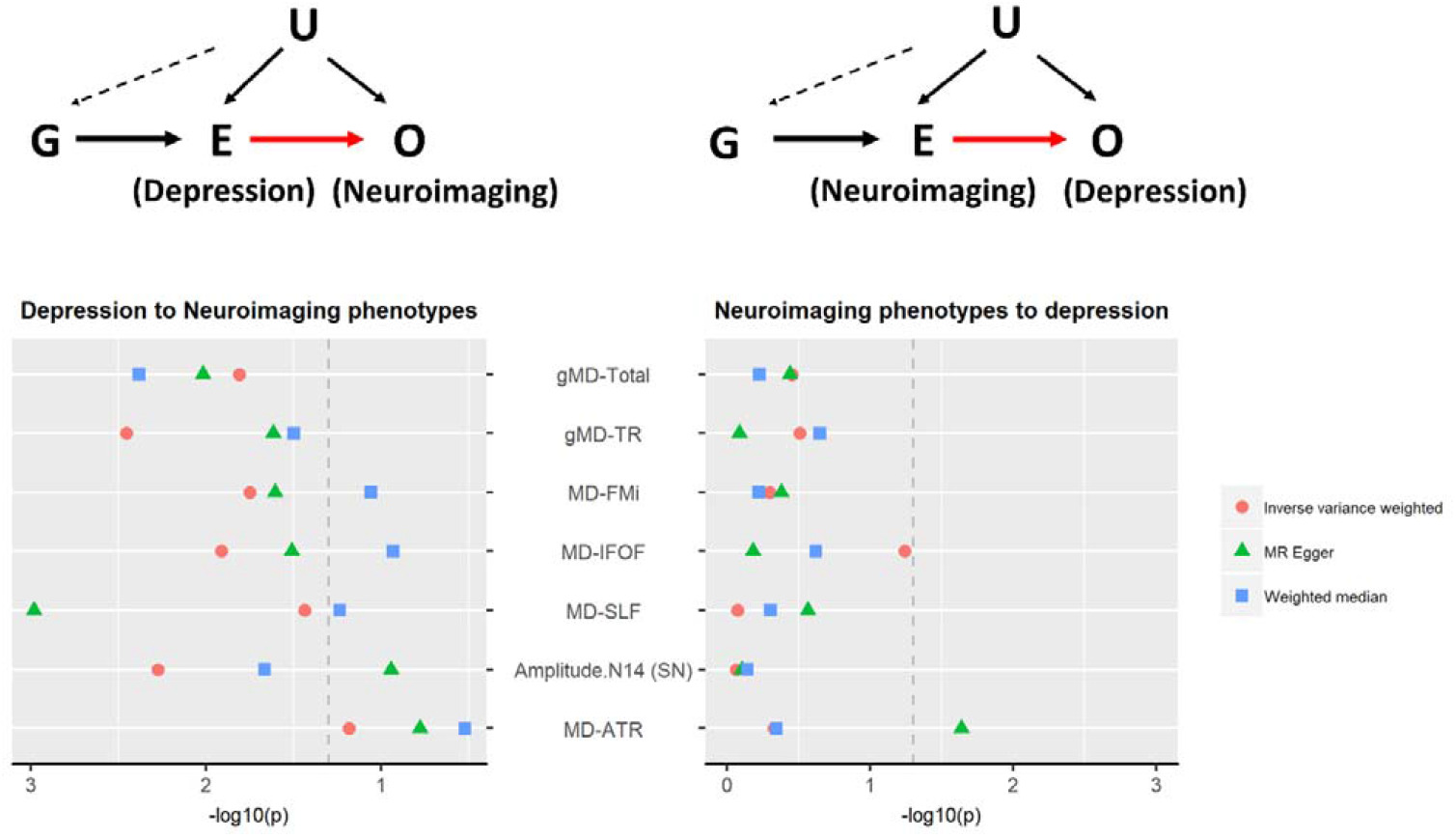
Bidirectional Mendelian Randomisation analysis for the association between neuroimaging phenotypes and depression. The left panel shows the model and results for Mendelian Randomisation results for the causal effect of depression to neuroimaging phenotypes, and the right panel shows the model and results for effect of neuroimaging phenotypes to depression. For the model illustrations, G=genetic instruments extracted from GWAS summary statistics of the exposure, E=exposure variable, O=outcome variable, U=unmeasured confounders (have no systematic association with G). In the scatter plots, x axes represent -log10 transformed p values for the Mendelian Randomisation results, and the y axes represent the neuroimaging traits tested in the models. Three types dots represent the three Mendelian Randomisation methods used. Dashed grey lines are the p=0.05 threshold for nominal significance. FDR-corrected p values were reported in supplementary materials. In the figure, MD = mean diffusivity, TR = thalamic radiations, FMi = forceps minor, IFOF = inferior fronto-occipital fasciculus, SLF = superior longitudinal fasciculus, ATR = anterior thalamic radiation and Amplitude.N14 (SN) = fluctuation amplitude in Node 14 (i.e. the Salience Network).

GWAS summary statistics for depression came from the meta-analysis used to generate the PRS as described above. For the neuroimaging variables, the ones that were found associated with depression-PRS in both the discovery and replication samples were chosen. GWAS were conducted using BGENIE impv3 on these neuroimaging variables in the UK Biobank imaging sample. Genetic data quality check, steps to ensure sample homogeneity and covariates mirrored the settings for the depression GWAS. The neuroimaging variables were scaled to obtain standardised estimates. SNP-heritability of depression and number of genome-wide significant hits were reported elsewhere^6^. SNP-heritability of white matter microstructure measures estimated using LDSC ranged from 12.3% to 29.2% and resting-state fluctuation amplitude ranged from 13.1% to 15.7%. The number of genome-wide significant loci ranged from 4 to 11 for all neuroimaging phenotypes. More details of neuroimaging GWAS summary statistics can be found in Table S2.

To test the causal effect of depression on neuroimaging variables, genetic instruments were chosen from the GWAS summary statistics of depression^6^, at a p threshold of 5×10^-8^. These SNPs were then clumped with a distance of 3,000 kb and a maximum LD r^2^ of 0.001, resulting in 101 independent genetic instruments. These SNPs were then identified within the GWAS summary statistics for each outcome, and those that were not present in both GWAS datasets were removed. SNP effect data on both the exposure and outcome were then harmonised to match the effect alleles before entering into MR analyses.

For the causal effects of neuroimaging variables on depression, genetic instruments were chosen at a lower p threshold of 5×10^-6^, due to a lack of genome-wide significant hits. At this threshold, after clumping with the same parameters as for choosing genetic instruments for depression, 20 to 29 independent genetic instruments were identified for each neuroimaging variable (see Table S3).

Three robust Mendelian Randomisation methods were chosen: MR-Egger, inverse-variance weighted estimator (IVW) and the weighted median method. We also conducted three additional analyses (i) to test for horizontal pleiotropy by estimating the MR-Egger intercept, and to test global heterogeneity of the genetic instruments using (ii) the Q test^38^ and (iii) the MR-Presso global test^39^.

P values were corrected separately for each category of neuroimaging measure (g of white matter measures/tract measures/resting-state fluctuation amplitude) and each MR method using FDR correction in R.

### Statistical models for mediation effect of neuroimaging variables

Following the PheWAS and Mendelian Randomisation analyses, we sought to test whether manifestations of depression were mediating the causal effect of depression-PRS on brain imaging phenotypes, as well as whether the neuroimaging variables act as neural mediators of genetic risk on depressive traits (i.e. neuroimaging traits were ‘endophenotypes’). These tests were applied using structural equational modelling (SEM) with the ‘lavaan’ package in R^40^. Two types of mediation analysis were conducted. The first one aimed to test whether the neuroimaging effects were the consequence of depression by testing if depression mediated the relationship between polygenic risk and neuroimaging variables (predictor=depression-PRS, mediator variable= CIDI definition of depression/depressive symptoms, and dependent variables=neuroimaging traits). Neuroimaging variables were chosen from those measures that showed a significant causal effect from depression in the Mendelian Randomisation analyses. The second type of mediation models tested whether neuroimaging variables mediated the relationship between polygenic risk of depression on depressive phenotypes (predictor=depression-PRS, mediator=neuroimaging traits, and dependent variable=CIDI definition of depression /depressive symptoms). The list of mediators was restricted to the neuroimaging phenotypes that showed significant causal effects on depression by Mendelian Randomisation analyses. For both types of mediation analyses, variables for manifestations of depression include CIDI definition for depression, severity of depression assessed by CIDI short form^26^ and the current symptoms at the imaging assessment measured by PHQ-4^41^. In order to maximise statistic power, all mediation tests used the full sample that included both discovery and replication datasets (N=22,888), adjusted for site.

All covariates remained the same as for PheWAS regression models. P value correction followed the same method as the Mendelian Randomisation analysis. Illustration for the models can be found in Figure S6, Table S4 and supplementary methods.

### Interactions of depression-PRS and early risk factors or sociodemographic variables

Interactions between environmental variables, previously associated with depression, and depression-PRS were also tested. Environmental variables were chosen from early life risk factors and sociodemographic variables previously found associated with risk for depression and showed depression case-control difference in present sample (p<0.05), which include: household income, Townsend Index, childhood trauma, adulthood trauma and recent stressful life events in the past six months before imaging assessment^42,43^.

Dependent variables were the behavioural and imaging phenotypes that showed significant associations with depression-PRS at minimum four thresholds in both the discovery and replication samples. Variables that were selected as factors were not included as dependent variables. The covariates included in these GxE analyses were identical to the PheWAS analyses. FDR correction was applied in the same manners with the PheWAS (m dependent variables * 8 p thresholds).

## Results

### PheWAS

We found that 93 phenotypes (68 behavioural and 25 neuroimaging) out of 488 examined (210 behavioural and 278 neuroimaging) in the discovery sample showed significant associations with depression-PRS at a minimum of four p thresholds after correction for multiple comparisons (absolute β: 0.014 to 0.341, p_FDR_: 0.046 to 3.20×10^-31^). Overall results for depression-PRS of representative p thresholds at 1 and 0.01 are presented in Figure 1. These two thresholds were selected since pT<1 and pT<0.01 showed the largest effect sizes in behavioural traits and neuroimaging phenotypes respectively (see Figure 2 and S7). Results for other thresholds can be found in Figure S7 and Table S5.

Ninety-two of the 93 traits showed an identical direction of effect in the replication sample (Figure 3, S8-9 and Table S5-6). After multiple comparison correction, 69 traits showed associations with depression-PRS at a minimum of four p-thresholds in the replication sample (52 behavioural and 17 neuroimaging). In total, 74.2% findings were replicated, with the highest replication rates for mental health variables (81.8%), physical measures (82.6%) and for white matter microstructure (78.9%), see Figure S7-8. Consistent results were also found between the MRI acquisition sites (see Figures S10-11 and Tables S7-9) and there was no significant interaction between MRI site and depression-PRS on any of the traits (p_cor_>0.583, see Figures S10-11). Further details on between-site comparisons can be found in the supplementary materials. Results for meta-analysis combining the two samples can be found in Figures S12-13 and Table S10.

Significant associations that were found in both the discovery and replication datasets are reported below. A complete list of all results is presented in Table S5-6.

#### 1) Depression-PRS associations with definitions for depression and symptomology

Higher depression-PRS were associated with the presence of depression based on all three definitions, including broad depression (β: 0.154 to 0.300, p_FDR_: 3.51×10^-9^ to 3.20×10^-31^), probable depression (β: 0.174 to 0.341, p_FDR_: 1.00×10^-6^ to 1.35×10^-23^), and CIDI depression (β: 0.121 to 0.261 p_FDR_: 2.75×10^-4^ to 1.04×10^-17^).

Significant associations were also found between depression-PRS and depressive symptoms, assessed by PHQ-4 (Patient Health Questionnaire) and CIDI questionnaires, and other self-reported psychological traits including self-harm, subjective well-being, reported feeling of not worth living and neuroticism (absolute β: 0.027 to 0.339, p_FDR_: 0.043 to 1.26×10^-25^).

#### 2) Associations between depression-PRS and white matter microstructure

Brain structural phenotypes of white matter microstructure were associated with depression-PRS. Higher depression-PRS were in general associated with decreased white matter microstructural integrity. General changes of lower global fractional anisotropy (FA) and higher global mean diffusivity (MD) (absolute β: 0.023 to 0.040, p_FDR_: 0.045 to 3.92×10^-4^) were associated with higher depression-PRS. Lower microstructural integrity was also shown in the general measures of FA and MD for two subsets of white matter tracts, the association fibres (absolute β: 0.029 to 0.041, p_FDR_: 0.025 to 3.56×10^-4^) and thalamic radiations (absolute β: 0.025 to 0.035, p_FDR_: 0.037 to 2.32×10^-3^). For each individual tract (Figures 2 and 4), higher depression-PRS were associated with decreased FA in posterior thalamic radiation, superior longitudinal fasciculus and forceps major (β: −0.025 to −0.040, p_FDR_: 0.045 to 8.17×10^-4^), and increased MD in anterior thalamic radiation, superior thalamic radiation, cingulate gyrus part of cingulum, inferior fronto-occipital fasciculus, superior longitudinal fasciculus and forceps minor (β: 0.023 to 0.039, p_FDR_: 0.044 to 3.92×10^-4^).

#### 3) Depression-PRS associations with resting-state fluctuation amplitude

Associations were found between depression-PRS and resting-state fluctuation amplitude of low-frequency signal (β: 0.026 to 0.044, p_FDR_: 0.044 to 9.91×10^-5^) in the discovery sample (Figures 2 and 4). A full list of report is presented in Table S11.

In brief, higher depression-PRS were associated with lower fluctuation amplitude in anterior cingulate gyrus (peak coordination: −10, 54, 2; cluster size: 6,679), bilateral postcentral gyrus (peak coordination: −44, −30, 46 and 44, −24, 40 for left and right hemispheres respectively; cluster sizes: 2,278 and 1,184), bilateral insula (peak coordination: −38, −4, 16 and 30, 18, −16 for left and right hemispheres respectively; cluster sizes: 811 and 300), left thalamus (peak coordination: −2, −18, 10, cluster size: 216), bilateral orbital part of inferior frontal gyrus (peak coordination: −34, 34, −12 and 32, 36, −10 for left and right hemispheres respectively; cluster sizes: 165 and 177) and left superior frontal lobe (peak coordination: −18, 34, 40; cluster size: 111). These regions are largely contained within the salience, executive control and sensorimotor networks (Table S11)^44,45^.

#### 4) Depression-PRS associations with sleep problems, smoking and poor physical health

In the category of lifestyle measures, reporting of sleep problems (e.g. too much sleep or insomnia) (absolute β: 0.034 to 0.180, p_FDR_: 0.04 to 7.42×10^-9^), and smoking behaviours (absolute β: 0.044 to 0.105, p_FDR_: 2.08×10^-3^ to 3.38×10^-8^) were found to be significantly positively associated with depression-PRS.

Physical health items associated with depression-PRS can be summarised as the following four categories: (1) self-reported overall health rating and conditions of long-standing illnesses (absolute β: 0.034 to 0.077, p_FDR_: 4.08×10^-3^ to 1.32×10^-13^), (2) recent pains and on-going treatment (absolute β: 0.040 to 0.080, p_FDR_: 5.50×10^-4^ to 8.54×10^-15^), (3) cardiovascular/heart problems (absolute β: 0.027 to 0.046, p_FDR_: 0.027 to 1.79×10^-5^), and (4) body mass and weight change compared to one year ago (absolute β: 0.014 to 0.042, p_FDR_: 0.046 to 4.40×10^-6^).

### Bidirectional Mendelian Randomisation: causal effect from depression to neuroimaging phenotypes, and vice versa

A significant causal effect of depression was found on lower microstructural integrity in five white matter microstructural measures and lower resting-state fluctuation amplitude in the Salience Network (Node 14). For these phenotypes, the effect from depression were shown in at least two MR methods after FDR correction (Figure 5, Table S3 and Figure S14-20). The neuroimaging phenotypes include (β and p_FDR_ reported for significant effects): global gMD (gMD-Total; β: 0.125 to 0.707, p_FDR_: 0.029 to 0.012, significant for all three MR methods), gMD in thalamic radiations (gMD-TR; β: 0.131 to 0.545, p_FDR_: 0.048 to 0.011, all three MR methods), MD in forceps minor (MD-FMi; β: 0.126 to 0.636, p_FDR_: 0.041 to 0.035, IVW and MR Egger), MD in inferior fronto-occipital fasciculus (MD-IFOF; β: 0.120 to 0.558, p_FDR_: 0.041 to 0.035, IVW and MR Egger), MD in superior longitudinal fasciculus (MD-SLF; β: 0.115 to 0.945, p_FDR_: 0.049 to 0.004, IVW and MR Egger) and the resting-state fluctuation amplitude in the Salience Network (amp-N14; β: −0.136 to −0.164, p_FDR_: 0.043 to 0.011, IVW and the weighted median). No significant reversed effect of these neuroimaging phenotypes on depression was found (p ranged from 0.886 to 0.229). For the above significant effects, MD in superior longitudinal fasciculus showed significant horizontal pleiotropy (p_FDR_ for MR-Egger intercepts=0.013), and however, result for MR-Egger was significant as reported, which is a more robust method compared with the other two when a significant average pleiotropy is detected^46^. gMD in association fibres (gMD-AF) showed significant global SNP heterogeneity (p_FDR_ for MR-Presso global test=0.023). A subsequent MR-Presso distortion test showed that removing the SNP outlier did not make a significant difference to the estimation of causal effect on gMD-AF (p for MR-Presso distortion test = 0.138). No other test showed significant horizontal pleiotropy or SNP heterogeneity (p_FDR_ for MR-Egger intercept > 0.080, and p_FDR_ for MR-Presso global test > 0.190, and p_FDR_ for all Q tests > 0.105).

Conversely, the directional effect of neuroimaging phenotypes on depression were tested, and no effect reached statistical significance after FDR correction. The only nominally significant effect was shown from MD in anterior thalamic radiation to depression for MR-Egger method (MD-ATR; β=0.131, p=0.023, p_FDR_=0.092, p for MR-Egger intercept=0.030, p for MR-Presso global test=0.063, p for Q statistics=0.067).

### Mediation analyses: neuroimaging traits as the mediators of risk or outcomes of depression

In the first mediation model, we tested if polygenic risk of depression led to changes in several neuroimaging variables through the mediating effects of depression. The neuroimaging variables were chosen if they presented as a significant causal consequence of depression in the Mendelian Randomisation analyses. Conversely, in the second model the neuroimaging variable of MD in anterior thalamic radiation showed a potentially causal effect on depression at nominal significance using Mendelian Randomisation and was therefore tested for its potential role as a mediator of genetic risk on depression. Here we report the results for depression-PRS at the threshold of pT<1. For other depression-PRS thresholds, see Table S4.

We found evidence that current depressive symptoms mediated the effect of depression-PRS on: global MD (gMD-Total; β=0.002, p_FDR_=0.003), MD in thalamic radiations (gMD-TR; β=0.002, P_FDR_=0.002) and MD in superior longitudinal fasciculus (β=0.002, p_FDR_=0.040). Conversely, a significant mediation effect of MD in anterior thalamic radiation was found, mediating the effect of depression-PRS on current depressive symptoms (PHQ-4) (β=0.001, p=0.022). All significant mediation models showed good model fit characteristics (CLI ranged from 0.983 to 0.992, TLI ranged from 0.977 to 0.986, and all p_RMSEA_=1)· A full list of results for all mediation models tested can be found in Table S4.

### Interaction of depression-PRS with early life risk factors and sociodemographic variables

Environmental variables that showed significant interaction with depression-PRS included childhood trauma, Townsend Index and recent stressful life events. The dependent variables that were significantly identified as showing evidence of GxE were mainly measures of mental health, including depressive symptoms, manifestations of other psychiatric conditions such as general anxiety disorder (GAD), bipolar disorder (BPD) and post-traumatic stress disorder (PTSD) and overall health rating (see Figure 6 and S14, p_FDR_ < 0.049).

**Figure 6.**
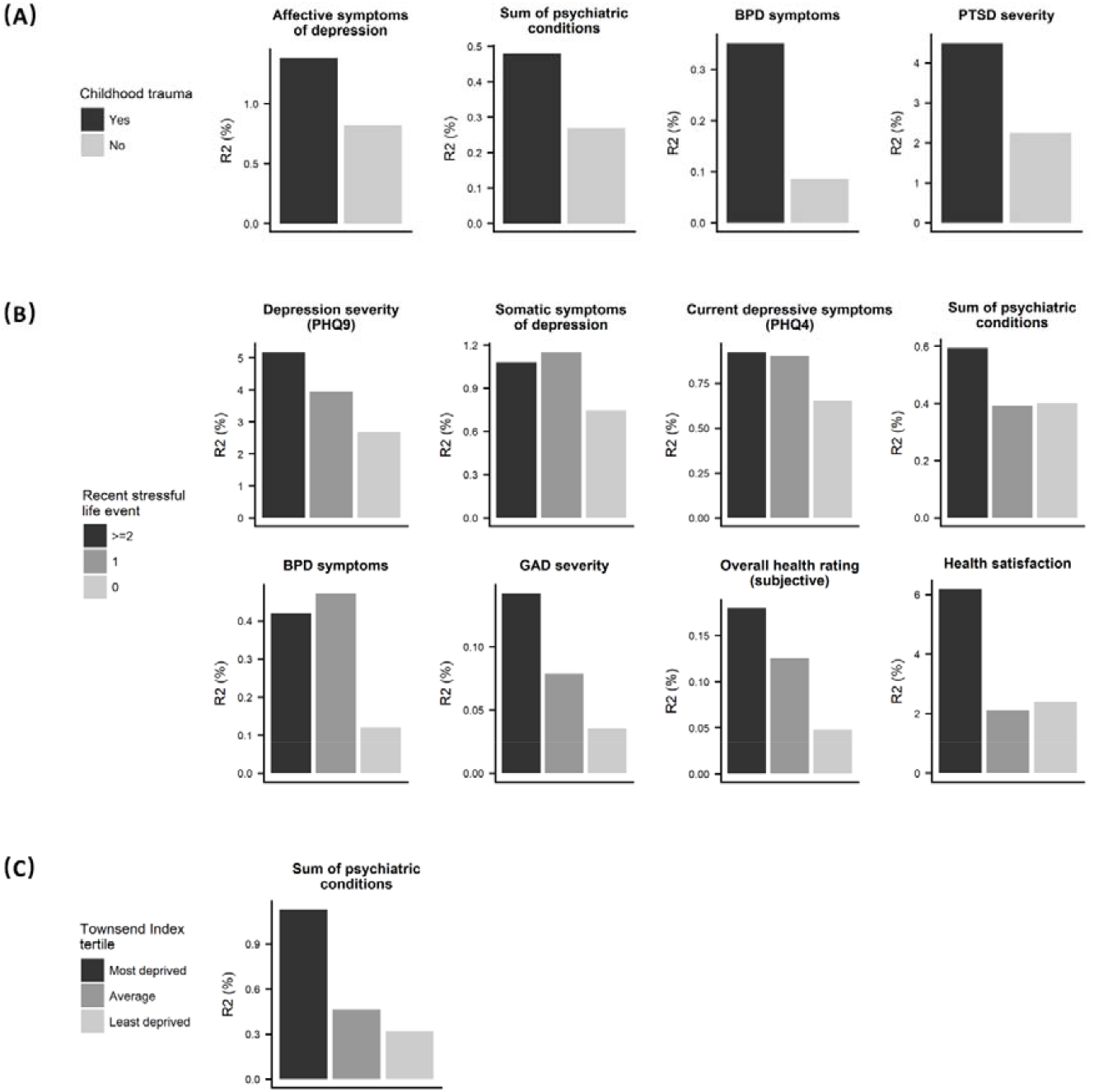
Variance explained by depression-PRS under the exposure of different environmental risk factors. The colour shade of each bar represents one condition of environmental factor, a darker shade represents a risk-conferring condition (i.e. had reported childhood trauma, had more than 2 recent stressful life events and in the most deprived area). The y axes represent the variance explained (R^2^ in %) by depression-PRS under the given environmental conditions.

In general, the effect of depression-PRS was enhanced in participants exposed to more adverse social/socioeconomic environments. (1) In participants that reported any childhood trauma versus none, the variance in the dependent variables accounted for by depression-PRS were 1.69 to 4.08 times higher for the total number of psychiatric conditions, ever having PTSD, BPD symptoms and affective symptoms of depression. (2) For those who had at least two recent stressful life events in six months, variance explained by depression-PRS were higher than those reported none at the scale of 1.42 to 4.04 times, in variables of depressive symptoms assessed by PHQ-9, somatic symptoms of depression, current depressive symptoms, GAD severity, BPD symptoms, subjective rating of overall physical health and health satisfaction. (3) Finally, for participants in the most deprived tertile band, variance explained in the sum of psychiatric conditions was 3.58 times higher than for the least deprived participants. Detailed reports can be found in Figure 6, S21 and Tables S12-16.

We found no evidence of interactions however between depression-PRS and either adulthood trauma or household income (p_FDR_>0.333).

## Discussion

Replicated associations between depression-PRS, behavioural and neuroimaging phenotypes were found in the present study using the largest independent imaging cohort to date. The strongest associations were found between depression-PRS and mental health variables. Several novel associations were detected, including associations between depression-PRS and both white matter microstructure and fluctuation amplitude of low-frequency resting-state signals. In particular, MD in the anterior thalamic radiation was found to have a causal impact on depression, and it was found to significantly mediate the relationship between depression-PRS and current depressive symptoms. In addition, Mendelian Randomisation analysis also showed evidence for changes in the MD of thalamic radiations and in superior longitudinal fasciculus that were likely to be a causal consequence of depression. Other associations with higher polygenic risk included abnormal self-reported sleep problems, smoking behaviour, cardiovascular conditions and increased body mass index. The findings regarding the interactions of early-life factors and sociodemographic variables with depression-PRS revealed that the effect of depression-PRS on mental health was stronger in participants that had reported childhood trauma, had multiple recent stressful life events and experienced socioeconomic deprivation.

Novel associations were found between depression-PRS and neuroimaging variables on structural connectivity and functional resting-state fluctuation amplitude in the brain. Findings from both Diffusion Tensor Imaging and resting-state data revealed the importance of prefrontal cortex, which is a hub for emotion regulation and executive control^47,48^. The role of the prefrontal cortex is further supported by the latest GWAS on depression, which showed the enrichment of risk-associated genes in this region^23^. In particular, white matter microstructure showed the largest effect sizes among brain phenotypes in our results, and most trait associations in this category were replicated in an independent dataset. The current findings therefore indicate a potentially risk-conferring role for white matter over other modalities. This finding is supported by previous evidence that white matter microstructure has stronger phenotypic associations with lifetime depression compared to brain structural volumes^29^ and higher SNP-heritability (20-60%) compared with other neuroimaging modalities, indicating a greater genomic contribution to individual differences in phenotypes^49^. The present paper provides evidence of associations of white matter microstructure with depression-PRS, however, these brain measures have a relatively low spatial resolution. Recent gene expression studies suggest that genetic predisposition may influence more spatially and functionally specific, neuronal-level activities such as synaptic pruning and the overproduction of synapses^50^ for regional segregation^51^ during the process of brain maturation and myelin repair which contribute largely to brain structural and functional individual variance^49^. These highly regional and functionally specific brain phenotypes are of great importance and may help explain how genetic predisposition contributes to variance in neuroimaging measures. Future large-scale genetic association studies are necessary to further replicate and extend the current findings.

Several associations between polygenic risk of depression and neuroimaging variables were subsequently identified, through Mendelian Randomisation analysis, to have directional or causal significance. Whether brain structural and functional alterations are the outcome or cause of depressive symptoms has long been debated^52^. Our results show that some brain structural and functional alterations are likely to be an outcome of depression, however whether other imaging features are also a cause is yet unclear. Although our results for the causal effect from neuroimaging phenotype to depression were null, therefore suggesting a possibly uni-directional relationship from depression to the brain, it may be premature to draw a confident conclusions from such insignificant effects. It is important to consider that the relative lack of genome-wide significant loci for most neuroimaging measures provides weaker genetic instruments for Mendelian Randomisation, which may reduce power to detect such causal associations. There is currently a global effort to conduct GWAS using neuroimaging phenotypes and these efforts are likely to provide stronger genetic instruments for future analyses. Further, white matter microstructure in anterior thalamic radiation (ATR) did demonstrate a nominally significant causal effect on depression, but notably not in the reverse direction (from depression to ATR). This is in spite of the reverse direction of testing (from depression to ATR) having a much larger set of genetic instruments and greater power to detect significant effects. This indicates that the white matter microstructure in ATR may be one of the strongest neuroimaging candidates as a causal mediator of risk for depression.

The associations found in behavioural traits with depression-PRS suggest that polygenic risk of depression may also identify a predisposition to experience particular environmental risk exposures, or a vulnerability to their effects and later recall. Firstly, the linear association of depression-PRS with sleep, recent pains, smoking behaviour and whether there is any heart/cardiovascular condition showed the largest effect sizes. One important commonality of these behavioural patterns and physical conditions is that they all have a significant impact or reciprocal association with activities in the hypothalamic-pituitary-adrenal (HPA) axis^53^. Activity of the HPA axis is a well-replicated vulnerability factor for the onset of depression^53^, it is associated with brain development and synaptic formation^54,55^, and often engages under external stress-inducing environmental stimuli^56^. Secondly and more directly, the environmental risk factors tested in this study consistently strengthened the effect of depression-PRS. Compared with previous studies that test genetic-environment (G×E) interactions, the present study revealed that the G×E effect can present on a whole-genome, polygenic level. It may be a manifestation of interactions between the environmental risk factors and some important endophenotypes (e.g. HPA-axis activity) that polygenic risk of depression confers upon.

One potential limitation is that the present study initially focused on association tests regardless of causality. Between endophenotypes, environmental factors, disease outputs and compensatory adaptations, the boundaries are ambiguous, and yet they may show very similar associations with polygenic risk of depression. It is necessary to categorise them, therefore to help specify how to use the sub-diagnostic information. The present study intended to investigate the role of environmental factors and endophenotypes on a very limited number of traits, largely dependent on prior knowledge, for the purpose of rigorousness. However, to specify the roles of some other traits that showed strong associations with depression-PRS whilst absent of certain functionalities would largely broaden the scope for downstream analyses (e.g. smoking, sleeping and recent pains). Methods such as Mendelian Randomisation provides evidence about causal inferences^57^, as shown in the analysis for the association between brain imaging phenotypes and depression. However, larger samples for genetic studies on neuroimaging traits would largely benefit such analysis in order to balance the statistic power of clinical and neuroimaging phenotypes. Secondly, the summary statistics we used was based on GWAS that included some cases identified by self-declared depressive symptoms. As it has been argued in previous papers, the self-declared phenotypes may, to some extent, be more lenient than clinically identified traits, however, the statistic power can largely overcome the noise introduced by a small amount of misclassification, which was supported by a very high genetic correlation between self-declared depression and clinically validated depression^5,6^.

To conclude, a novel and relatively unconstrained approach was used to test for associations between depression-PRS and various behavioural and neuroimaging variables of likely relevance for depression. The findings revealed that white matter microstructure, general mental and physical health and behaviours such as sleep patterns and smoking behaviour were associated with PRS of depression. In terms of likely causal directions of effect, our findings suggest that most neuroimaging associations with depression are likely to be the causal consequence of depression. Only microstructure in the anterior thalamic radiation appeared to be causally mediating the relationship between genetic risk and depression.

## Acknowledgements

The present study is supported by a Wellcome Trust Strategic Award “Stratifying Resilience and Depression Longitudinally” (STRADL) (Reference 104036/Z/14/Z) and MRC Mental Health Data Pathfinder Award (Reference MC_PC_17209). Data acquisition and analyses were conducted using the UK Biobank Resource under approved project #4844. Participation of the UK Biobank subjects is gratefully appreciated. We also thank UK Biobank team for collecting and preparing data for analyses. Funding from the Biotechnology and Biological Sciences Research Council (BBSRC) and Medical Research Council (MRC) is gratefully acknowledged. The PGC has received major funding from the US National Institute of Mental Health (5 U01MH109528-03).

The participants and employees of 23andMe, Inc. are greatly appreciated for their contribution to the present study. We particularly thank the contributors from 23andMe: Michelle Agee, Babak Alipanahi, Adam Auton, Robert K. Bell, Katarzyna Bryc, Sarah L. Elson, Pierre Fontanillas, Nicholas A. Furlotte, David A. Hinds, Karen E. Huber, Aaron Kleinman, Nadia K. Litterman, Jennifer C. McCreight, Matthew H. McIntyre, Joanna L. Mountain, Elizabeth S. Noblin, Carrie A.M. Northover, Steven J. Pitts, J. Fah Sathirapongsasuti, Olga V. Sazonova, Janie F. Shelton, Suyash Shringarpure, Chao Tian, Joyce Y. Tung, Vladimir Vacic, and Catherine H. Wilson.

XS receives support from China Scholarship Council (No. 201506040037). HCW is supported by a JMAS SIM fellowship from the Royal College of Physicians of Edinburgh and by an ESAT College Fellowship from the University of Edinburgh. DMH is supported by a Sir Henry Wellcome Postdoctoral Fellowship (Reference 213674/Z/18/Z) and a 2018 NARSAD Young Investigator Grant from the Brain & Behavior Research Foundation (Reference 27404)

AMM is supported by the Sackler Trust. The authors declare no other conflicts of interests.

## References

1. Vigo, D., Thornicroft, G. & Atun, R. Estimating the true global burden of mental illness. The Lancet Psychiatry 3, 171–178 (2018).

2. World Health Organization. Depression and other common mental disorders: global health estimates. World Health Organization (2017). doi:CC BY-NC-SA 3.0 IGO

3. Sullivan, P. F., Neale, M. C. & Kendler, K. S. Genetic Epidemiology of Major Depression: Review and Meta-Analysis. Am. J. Psychiatry 157, 1552–1562 (2000).

4. Cipriani, A. et al. Comparative efficacy and acceptability of 21 antidepressant drugs for the acute treatment of adults with major depressive disorder: a systematic review and network meta-analysis. Lancet 391, 1357–1366 (2018).

5. Wray, N. R. et al. Genome-wide association analyses identify 44 risk variants and refine the genetic architecture of major depression. 50, (2018).

6. Howard, D. M. et al. Genome-wide meta-analysis of depression in 807,553 individuals identifies 102 independent variants with replication in a further 1,507,153 individuals. bioRxiv 6288, 433367 (2018).

7. Wray, N. R. et al. Research Review: Polygenic methods and their application to psychiatric traits. J. Child Psychol. Psychiatry 55, 1068–1087 (2014).

8. Shelton, R. C. The Course of Illness After Initial Diagnosis of Major Depression. JAMA Psychiatry 73, 3–4 (2016).

9. Cooney, R. E., Joormann, J., Eugène, F., Dennis, E. L. & Gotlib, I. H. Neural correlates of rumination in depression. Cogn. Affect. Behav. Neurosci. 10, 470–478 (2010).

10. DeRubeis, R. J., Siegle, G. J. & Hollon, S. D. Cognitive therapy versus medication for depression: Treatment outcomes and neural mechanisms. Nat. Rev. Neurosci. 9, 788–796 (2008).

11. Dunn, E. C. et al. Genetic determinants of depression: recent findings and future directions. Harv Rev Psychiatry 23,1–18 (2015).

12. Yirmiya, R., Rimmerman, N. & Reshef, R. Depression as a Microglial Disease. Trends Neurosci. 38,637–658 (2015).

13. Flint, J. & Kendler, K. S. The Genetics of Major Depression. Neuron 81, 484–503 (2014).

14. Glahn, D. C. et al. Arguments for the Sake of Endophenotypes: Examining Common Misconceptions About the Use of Endophenotypes in Psychiatric Genetics. Am. J. Med. Genet. 122–130 (2014). doi:10.1002/ajmg.b.32221

15. Kaiser, R. H., Andrews-Hanna, J. R., Wager, T. D. & Pizzagalli, D. A. Large-scale network dysfunction in major depressive disorder: A meta-analysis of resting-state functional connectivity. JAMA Psychiatry 72, 603–611 (2015).

16. Milaneschi, Y. et al. Polygenic dissection of major depression clinical heterogeneity. Mol Psychiatry (2015). doi:10.1038/mp.2015.86

17. Reus, L. M. et al. Association of polygenic risk for major psychiatric illness with subcortical volumes and white matter integrity in UK Biobank. Sci. Rep. 7,42140 (2017).

18. Miller, K. L. et al. Multimodal population brain imaging in the UK Biobank prospective epidemiological study. Nat. Neurosci. (2016). doi:10.1038/nn.4393

19. Peyrot, W. J. et al. Effect of polygenic risk scores on depression in childhood trauma. Br J Psychiatry 205,113–119 (2014).

20. Duncan, L. E. & Keller, M. C. A Critical Review of the First 10 Years of Candidate Gene-by-Environment Interaction Research in Psychiatry. Am. J. Psychiatry 168,1041–1049 (2011).

21. Sudlow, C. et al. UK Biobank: An Open Access Resource for Identifying the Causes of a Wide Range of Complex Diseases of Middle and Old Age. PLoS Med. 12,1–10 (2015).

22. Hyde, C. L. et al. Identification of 15 genetic loci associated with risk of major depression in individuals of European descent. Nat. Genet. 48,1031–1036 (2016).

23. Howard, D. M. et al. Genome-wide association study of depression phenotypes in UK Biobank identifies variants in excitatory synaptic pathways. Nat. Commun. 9, 1–10 (2018).

24. Euesden, J., Lewis, C. M. & O’Reilly, P. F. PRSice: Polygenic Risk Score software. Bioinformatics 31, 1466–1468 (2015).

25. Bycroft, C. et al. Genome-wide genetic data on 500,000 UK Biobank participants. bioRxiv 166298 (2017). doi:10.1101/166298

26. Davis, K. A. S. et at. Mental health in UK Biobank: development, implementation and results from an online questionnaire completed by 157 366 participants. BJPsych Open 4, 83–90 (2018).

27. Smith, D.J. et al. Prevalence and characteristics of probable major depression and bipolar disorder within UK biobank: cross-sectional study of 172,751 participants. PLoS One 8, e75362 (2013).

28. Deary, I. J., Penke, L. & Johnson, W. The neuroscience of human intelligence differences. Nat. Rev. Neurosci. 11, 201–211 (2010).

29. Shen, X. et at. Subcortical volume and white matter integrity abnormalities in major depressive disorder: Findings from UK Biobank imaging data. Sci. Rep. 7,1–10 (2017).

30. Shen, X. et at. Resting-state connectivity and its association with cognitive performance, educational attainment, and household income in UK Biobank. Biol. Psychiatry Cogn. Neurosci. Neuroimaging 1–9 (2018). doi:10.1016/J.BPSC.2018.06.007

31. Alfaro-Almagro, F. et at. Image processing and Quality Control for the first 10,000 brain imaging datasets from UK Biobank. Neuroimage 166,400–424 (2018).

32. Nelder, J. A. & Baker, R. J. Generalized linear models. Encycl. Stat. Sci. 4, (2004).

33. Pinheiro, J., Bates, D., DebRoy, S. & Sarkar, D. nlme: Linear and Nonlinear Mixed Effects Models. R Packag. version 3 1–97 (2007). at <http://scholar.google.com/scholar?hl=en&btnG=Search&q=intitle:nlme:+Unear+and+nonlinear+mixed+effects+models#3>

34. Benjamini, Y. & Hochberg, Y. On the Adaptive Control of the False Discovery Rate in Multiple Testing With Independent Statistics. J. Educ. Behav. Stat. 25, 60–83 (2000).

35. Smith, S. M. & Nichols, T. E. Statistical Challenges in ‘Big Data’ Human Neuroimaging NeuroView. Neuron 97, 263–268 (2018).

36. Bĳsterbosch, J. et al. Investigations into within- and between-subject resting-state amplitude variations. Neuroimage 159, 57–69 (2017).

37. Schmaal, L. et al. Subcortical brain alterations in major depressive disorder: findings from the ENIGMA Major Depressive Disorder working group. Mol. Psychiatry 21,806–812 (2016).

38. Hemani, G. et al. The MR-Base platform supports systematic causal inference across the human phenome. Elife 7, 1–29 (2018).

39. Verbanck, M., Chen, C. Y., Neale, B. & Do, R. Detection of widespread horizontal pleiotropy in causal relationships inferred from Mendelian randomization between complex traits and diseases. Nat. Genet. 50, 693–698 (2018).

40. Oberski, D. L. lavaan.survey: An R ackage for complex survey analysis of structural equation models. J. Stat. Softw. 57, 1–27 (2014).

41. Batty, G. D., McIntosh, A. M., Russ, T. C., Deary, I. J. & Gale, C. R. Psychological distress, neuroticism, and cause-specific mortality: Early prospective evidence from UK Biobank. J. Epidemiol. Community Health 70, 1136–1139 (2016).

42. Vancampfort, D. et al. Risk of metabolic syndrome and its components in people with schizophrenia and related psychotic disorders, bipolar disorder and major depressive disorder: A systematic review and meta-analysis. World Psychiatry 14, 339–347 (2015).

43. Sarkar, C., Webster, C. & Gallacher, J. Residential greenness and prevalence of major depressive disorders: a cross-sectional, observational, associational study of 94□879 adult UK Biobank participants. Lancet Planet. Heal. 2, e162–e173 (2018).

44. Uddin, L. Q. Salience processing and insular cortical function and dysfunction. Nat. Rev. Neurosci. 16, 55–61 (2014).

45. Kaiser, R. H., Andrews-Hanna, J. R., Wager, T. D. & Pizzagalli, D. A. Large-Scale Network Dysfunction in Major Depressive Disorder. JAMA Psychiatry 72, 603 (2015).

46. Burgess, S., Bowden, J., Fall, T., Ingelsson, E. & Thompson, S. G. Sensitivity analyses for robust causal inference from mendelian randomization analyses with multiple genetic variants. Epidemiology 28, 30–42 (2017).

47. Miller, E. K. The prefrontal cortex and cogitive control. Nat. Rev. neu 1, 59–65 (2000).

48. Etkin, A., Büchel, C. & Gross, J. J. The neural bases of emotion regulation. Nat. Rev. Neurosci. 16, 693–700 (2015).

49. Elliott, L. T. et al. Genome-wide association studies of brain structure and function in the UK Biobank. Nature 562, 210–216 (2018).

50. Petanjek, Z. et al. Extraordinary neoteny of synaptic spines in the human prefrontal cortex. Proc. Natl. Acad. Sci. 108,13281–13286 (2011).

51. Colantuoni, C. et al. Temporal dynamics and genetic control of transcription in the human prefrontal cortex. Nature 478, 519–523 (2011).

52. Glahn, D. C. et al. High Dimensional Endophenotype Ranking in the Search for Major Depression Risk Genes. Biol. Psychiatry 71, 6–14 (2011).

53. Pariante, C. M. & Lightman, S. L. The HPA axis in major depression: classical theories and new developments. Trends Neurosci. 31,464–468 (2008).

54. Stickgold, R. J. et al. Sleep and Synaptic Homeostasisff□: Science (80-.). 1576–1581 (2011). doi:10.1126/science.1202839

55. Maret, S., Faraguna, U., Nelson, A. B., Cirelli, C. & Tononi, G. Sleep and waking modulate spine turnover in the adolescent mouse cortex. Nat. Neurosci. 14, 1418–1420 (2011).

56. Pariante, C. M. Risk factors for development of depression and psychosis: Glucocorticoid receptors and pituitary implications for treatment with antidepressant and glucocorticoids. Ann. N. Y. Acad. Sci. 1179,144–152 (2009).

57. Smith, G. D. & Ebrahim, S. Mendelian randomization: prospects, potentials, and limitations. IntJ Epidemiol 33, 30–2 (2004).

58. Ruth, K. S. et al. Events in Early Life are Associated with Female Reproductive Ageing: A UK Biobank Study. Sci. Rep. 6, 1–9 (2016).

59. Hagenaars, S. P. et at. Shared genetic aetiology between cognitive functions and physical and mental health in UK Biobank (N = 112 151) and 24 GWAS consortia. bioRxiv 031120 (2015). doi:10.1101/031120

60. Hagenaars, S. P. et at. Shared genetic aetiology between cognitive functions and physical and mental health in UK Biobank (N=112 151) and 24 GWAS consortia. Mol. Psychiatry 21, 031120 (2016).

61. Hill, W. D. et al. Molecular genetic aetiology of general cognitive function is enriched in evolutionarily conserved regions. Transl. Psychiatry 6, e980 (2016).

