## Supplementary materials for "A phenome-wide association and Mendelian Randomisation study of polygenic risk for depression in UK Biobank"

**A phenome-wide association study of polygenic risk scores for depression using behavioural and neuroimaging phenotypes from UK Biobank**

Shen et al.

Division of Psychiatry, University of Edinburgh, Edinburgh, United Kingdom

**Major Depressive Disorder Working Group of the Psychiatric Genomics Consortium**

Naomi R Wray* 1, 2

Stephan Ripke* 3, 4, 5

Manuel Mattheisen* 6, 7, 8, 9

Maciej Trzaskowski* 1

Enda M Byrne 1

Abdel Abdellaoui 10

Mark J Adams 11

Esben Agerbo 9, 12, 13

Tracy M Air 14

Till F M Andlauer 15, 16

Silviu-Alin Bacanu 17

Marie Bækvad-Hansen 9, 18

Aartjan T F Beekman 19

Tim B Bigdeli 17, 20

Elisabeth B Binder 15, 21

Douglas H R Blackwood 11

Julien Bryois 22

Henriette N Buttenschøn 8, 9, 23

Jonas Bybjerg-Grauholm 9, 18

Na Cai 24, 25

Enrique Castelao 26

Jane Hvarregaard Christensen 7, 8, 9

Toni-Kim Clarke 11

Jonathan R I Coleman 27

Lucía Colodro-Conde 28

Baptiste Couvy-Duchesne 2, 29

Nick Craddock 30

Gregory E Crawford 31, 32

Gail Davies 33

Ian J Deary 33

Franziska Degenhardt 34, 35

Eske M Derks 28

Nese Direk 36, 37

Conor V Dolan 10

Erin C Dunn 38, 39, 40

Thalia C Eley 27

Valentina Escott-Price 41

Farnush Farhadi Hassan Kiadeh 42

Hilary K Finucane 43, 44

Jerome C Foo 45

Andreas J Forstner 34, 35, 46, 47

Josef Frank 45

Héléna A Gaspar 27

Michael Gill 48

Fernando S Goes 49

Scott D Gordon 28

Jakob Grove 7, 8, 9, 50

Lynsey S Hall 11, 51

Christine Søholm Hansen 9, 18

Thomas F Hansen 52, 53, 54

Stefan Herms 34, 35, 47

Ian B Hickie 55

Per Hoffmann 34, 35, 47

Georg Homuth 56

Carsten Horn 57

Jouke-Jan Hottenga 10

David M Hougaard 9, 18

Marcus Ising 58

Rick Jansen 19

Ian Jones 59

Lisa A Jones 60

Eric Jorgenson 61

James A Knowles 62

Isaac S Kohane 63, 64, 65

Julia Kraft 4

Warren W. Kretzschmar 66

Jesper Krogh 67

Zoltán Kutalik 68, 69

Yihan Li 66

Penelope A Lind 28

Donald J MacIntyre 70, 71

Dean F MacKinnon 49

Robert M Maier 2

Wolfgang Maier 72

Jonathan Marchini 73

Hamdi Mbarek 10

Patrick McGrath 74

Peter McGuffin 27

Sarah E Medland 28

Divya Mehta 2, 75

Christel M Middeldorp 10, 76, 77

Evelin Mihailov 78

Yuri Milaneschi 19

Lili Milani 78

Francis M Mondimore 49

Grant W Montgomery 1

Sara Mostafavi 79, 80

Niamh Mullins 27

Matthias Nauck 81, 82

Bernard Ng 80

Michel G Nivard 10

Dale R Nyholt 83

Paul F O'Reilly 27

Hogni Oskarsson 84

Michael J Owen 59

Jodie N Painter 28

Carsten Bøcker Pedersen 9, 12, 13

Marianne Giørtz Pedersen 9, 12, 13

Roseann E. Peterson 17, 85

Erik Pettersson 22

Wouter J Peyrot 19

Giorgio Pistis 26

Danielle Posthuma 86, 87

Jorge A Quiroz 88

Per Qvist 7, 8, 9

John P Rice 89

Brien P. Riley 17

Margarita Rivera 27, 90

Saira Saeed Mirza 36

Robert Schoevers 91

Eva C Schulte 92, 93

Ling Shen 61

Jianxin Shi 94

Stanley I Shyn 95

Engilbert Sigurdsson 96

Grant C B Sinnamon 97

Johannes H Smit 19

Daniel J Smith 98

Hreinn Stefansson 99

Stacy Steinberg 99

Fabian Streit 45

Jana Strohmaier 45

Katherine E Tansey 100

Henning Teismann 101

Alexander Teumer 102

Wesley Thompson 9, 53, 103, 104

Pippa A Thomson 105

Thorgeir E Thorgeirsson 99

Matthew Traylor 106

Jens Treutlein 45

Vassily Trubetskoy 4

André G Uitterlinden 107

Daniel Umbricht 108

Sandra Van der Auwera 109

Albert M van Hemert 110

Alexander Viktorin 22

Peter M Visscher 1, 2

Yunpeng Wang 9, 53, 104

Bradley T. Webb 111

Shantel Marie Weinsheimer 9, 53

Jürgen Wellmann 101

Gonneke Willemsen 10

Stephanie H Witt 45

Yang Wu 1

Hualin S Xi 112

Jian Yang 2, 113

Futao Zhang 1

Volker Arolt 114

Bernhard T Baune 115

Klaus Berger 101

Dorret I Boomsma 10

Sven Cichon 34, 47, 116, 117

Udo Dannlowski 114

EJC de Geus 10, 118

J Raymond DePaulo 49

Enrico Domenici 119

Katharina Domschke 120

Tõnu Esko 5, 78

Hans J Grabe 109

Steven P Hamilton 121

Caroline Hayward 122

Andrew C Heath 89

Kenneth S Kendler 17

Stefan Kloiber 58, 123, 124

Glyn Lewis 125

Qingqin S Li 126

Susanne Lucae 58

Pamela AF Madden 89

Patrik K Magnusson 22

Nicholas G Martin 28

Andrew M McIntosh 11, 33

Andres Metspalu 78, 127

Ole Mors 9, 128

Preben Bo Mortensen 8, 9, 12, 13

Bertram Müller-Myhsok 15, 16, 129

Merete Nordentoft 9, 130

Markus M Nöthen 34, 35

Michael C O'Donovan 59

Sara A Paciga 131

Nancy L Pedersen 22

Brenda WJH Penninx 19

Roy H Perlis 38, 132

David J Porteous 105

James B Potash 133

Martin Preisig 26

Marcella Rietschel 45

Catherine Schaefer 61

Thomas G Schulze 45, 93, 134, 135, 136

Jordan W Smoller 38, 39, 40

Kari Stefansson 99, 137

Henning Tiemeier 36, 138, 139

Rudolf Uher 140

Henry Völzke 102

Myrna M Weissman 74, 141

Thomas Werge 9, 53, 142

Cathryn M Lewis 27, 143

Douglas F Levinson 144

Gerome Breen 27, 145

Anders D Børglum 7, 8, 9

Patrick F Sullivan 22, 146, 147,

1, Institute for Molecular Bioscience, The University of Queensland, Brisbane, QLD, AU

2, Queensland Brain Institute, The University of Queensland, Brisbane, QLD, AU

3, Analytic and Translational Genetics Unit, Massachusetts General Hospital, Boston, MA, US

4, Department of Psychiatry and Psychotherapy, Universitätsmedizin Berlin Campus Charité Mitte, Berlin, DE

5, Medical and Population Genetics, Broad Institute, Cambridge, MA, US

6, Department of Psychiatry, Psychosomatics and Psychotherapy, University of Wurzburg, Wurzburg, DE

7, Centre for Psychiatry Research, Department of Clinical Neuroscience, Karolinska Institutet, Stockholm, SE

8, Department of Biomedicine, Aarhus University, Aarhus, DK

9, Dept of Biological Psychology & EMGO+ Institute for Health and Care Research, Vrije Universiteit Amsterdam, Amsterdam, NL

10, Division of Psychiatry, University of Edinburgh, Edinburgh, GB

11, Centre for Integrated Register-based Research, Aarhus University, Aarhus, DK

12, National Centre for Register-Based Research, Aarhus University, Aarhus, DK

13, iPSYCH, The Lundbeck Foundation Initiative for Integrative Psychiatric Research,, DK

14, Discipline of Psychiatry, University of Adelaide, Adelaide, SA, AU

15, Department of Translational Research in Psychiatry, Max Planck Institute of Psychiatry, Munich, DE

16, Department of Neurology, Klinikum rechts der Isar, Technical University of Munich, Munich, DE

17, Department of Psychiatry, Virginia Commonwealth University, Richmond, VA, US

18, Center for Neonatal Screening, Department for Congenital Disorders, Statens Serum Institut, Copenhagen, DK

19, Department of Psychiatry, Vrije Universiteit Medical Center and GGZ inGeest, Amsterdam, NL

20, Virginia Institute for Psychiatric and Behavior Genetics, Richmond, VA, US

21, Department of Psychiatry and Behavioral Sciences, Emory University School of Medicine, Atlanta, GA, US

22, Department of Medical Epidemiology and Biostatistics, Karolinska Institutet, Stockholm, SE

23, Department of Clinical Medicine, Translational Neuropsychiatry Unit, Aarhus University, Aarhus, DK

24, iSEQ, Centre for Integrative Sequencing, Aarhus University, Aarhus, DK

25, Human Genetics, Wellcome Trust Sanger Institute, Cambridge, GB

26, Statistical genomics and systems genetics, European Bioinformatics Institute (EMBL-EBI), Cambridge, GB

27, Department of Psychiatry, University Hospital of Lausanne, Prilly, Vaud, CH

28, Social Genetic and Developmental Psychiatry Centre, King's College London, London, GB

29, Genetics and Computational Biology, QIMR Berghofer Medical Research Institute, Brisbane, QLD, AU

30, Centre for Advanced Imaging, The University of Queensland, Brisbane, QLD, AU

31, Psychological Medicine, Cardiff University, Cardiff, GB

32, Center for Genomic and Computational Biology, Duke University, Durham, NC, US

33, Department of Pediatrics, Division of Medical Genetics, Duke University, Durham, NC, US

34, Centre for Cognitive Ageing and Cognitive Epidemiology, University of Edinburgh, Edinburgh, GB

35, Institute of Human Genetics, University of Bonn, School of Medicine & University Hospital Bonn, Bonn, DE

36, Epidemiology, Erasmus MC, Rotterdam, Zuid-Holland, NL

37, Psychiatry, Dokuz Eylul University School Of Medicine, Izmir, TR

38, Department of Psychiatry, Massachusetts General Hospital, Boston, MA, US

39, Psychiatric and Neurodevelopmental Genetics Unit (PNGU), Massachusetts General Hospital, Boston, MA, US

40, Stanley Center for Psychiatric Research, Broad Institute, Cambridge, MA, US

41, Neuroscience and Mental Health, Cardiff University, Cardiff, GB

42, Bioinformatics, University of British Columbia, Vancouver, BC, CA

43, Department of Epidemiology, Harvard T.H. Chan School of Public Health, Boston, MA, US

44, Department of Mathematics, Massachusetts Institute of Technology, Cambridge, MA, US

45, Department of Genetic Epidemiology in Psychiatry, Central Institute of Mental Health, Medical Faculty Mannheim, Heidelberg University, Mannheim, Baden-Württemberg, DE

46, Department of Psychiatry (UPK), University of Basel, Basel, CH

47, Department of Biomedicine, University of Basel, Basel, CH

48, Centre for Human Genetics, University of Marburg, Marburg, DE

49, Department of Psychiatry, Trinity College Dublin, Dublin, IE

50, Psychiatry & Behavioral Sciences, Johns Hopkins University, Baltimore, MD, US

51, Bioinformatics Research Centre, Aarhus University, Aarhus, DK

52, Institute of Genetic Medicine, Newcastle University, Newcastle upon Tyne, GB

53, Danish Headache Centre, Department of Neurology, Rigshospitalet, Glostrup, DK

54, Institute of Biological Psychiatry, Mental Health Center Sct. Hans, Mental Health Services Capital Region of Denmark, Copenhagen, DK

55, iPSYCH, The Lundbeck Foundation Initiative for Psychiatric Research, Copenhagen, DK

56, Brain and Mind Centre, University of Sydney, Sydney, NSW, AU

57, Interfaculty Institute for Genetics and Functional Genomics, Department of Functional Genomics, University Medicine and Ernst Moritz Arndt University Greifswald, Greifswald, Mecklenburg-Vorpommern, DE

58, Roche Pharmaceutical Research and Early Development, Pharmaceutical Sciences, Roche Innovation Center Basel, F. Hoffmann-La Roche Ltd, Basel, CH

59, Max Planck Institute of Psychiatry, Munich, DE

60, MRC Centre for Neuropsychiatric Genetics and Genomics, Cardiff University, Cardiff, GB

61, Department of Psychological Medicine, University of Worcester, Worcester, GB

62, Division of Research, Kaiser Permanente Northern California, Oakland, CA, US

63, Psychiatry & The Behavioral Sciences, University of Southern California, Los Angeles, CA, US

64, Department of Biomedical Informatics, Harvard Medical School, Boston, MA, US

65, Department of Medicine, Brigham and Women's Hospital, Boston, MA, US

66, Informatics Program, Boston Children's Hospital, Boston, MA, US

67, Wellcome Trust Centre for Human Genetics, University of Oxford, Oxford, GB

68, Institute of Social and Preventive Medicine (IUMSP), University Hospital of Lausanne, Lausanne, VD, CH

69, Swiss Institute of Bioinformatics, Lausanne, VD, CH

70, Division of Psychiatry, Centre for Clinical Brain Sciences, University of Edinburgh, Edinburgh, GB

71, Mental Health, NHS 24, Glasgow, GB

72, Department of Psychiatry and Psychotherapy, University of Bonn, Bonn, DE

73, Statistics, University of Oxford, Oxford, GB

74, Psychiatry, Columbia University College of Physicians and Surgeons, New York, NY, US

75, School of Psychology and Counseling, Queensland University of Technology, Brisbane, QLD, AU

76, Child and Youth Mental Health Service, Children's Health Queensland Hospital and Health Service, South Brisbane, QLD, AU

77, Child Health Research Centre, University of Queensland, Brisbane, QLD, AU

78, Estonian Genome Center, University of Tartu, Tartu, EE

79, Medical Genetics, University of British Columbia, Vancouver, BC, CA

80, Statistics, University of British Columbia, Vancouver, BC, CA

81, DZHK (German Centre for Cardiovascular Research), Partner Site Greifswald, University Medicine, University Medicine Greifswald, Greifswald, Mecklenburg-Vorpommern, DE

82, Institute of Clinical Chemistry and Laboratory Medicine, University Medicine Greifswald, Greifswald, Mecklenburg-Vorpommern, DE

83, Institute of Health and Biomedical Innovation, Queensland University of Technology, Brisbane, QLD, AU

84, Humus, Reykjavik, IS

85, Virginia Institute for Psychiatric & Behavioral Genetics, Virginia Commonwealth University, Richmond, VA, US

86, Clinical Genetics, Vrije Universiteit Medical Center, Amsterdam, NL

87, Complex Trait Genetics, Vrije Universiteit Amsterdam, Amsterdam, NL

88, Solid Biosciences, Boston, MA, US

89, Department of Psychiatry, Washington University in Saint Louis School of Medicine, Saint Louis, MO, US

90, Department of Biochemistry and Molecular Biology II, Institute of Neurosciences, Center for Biomedical Research, University of Granada, Granada, ES

91, Department of Psychiatry, University of Groningen, University Medical Center Groningen, Groningen, NL

92, Department of Psychiatry and Psychotherapy, University Hospital, Ludwig Maximilian University Munich, Munich, DE

93, Institute of Psychiatric Phenomics and Genomics (IPPG), University Hospital, Ludwig Maximilian University Munich, Munich, DE

94, Division of Cancer Epidemiology and Genetics, National Cancer Institute, Bethesda, MD, US

95, Behavioral Health Services, Kaiser Permanente Washington, Seattle, WA, US

96, Faculty of Medicine, Department of Psychiatry, University of Iceland, Reykjavik, IS

97, School of Medicine and Dentistry, James Cook University, Townsville, QLD, AU

98, Institute of Health and Wellbeing, University of Glasgow, Glasgow, GB

99, deCODE Genetics / Amgen, Reykjavik, IS

100, College of Biomedical and Life Sciences, Cardiff University, Cardiff, GB

101, Institute of Epidemiology and Social Medicine, University of Münster, Münster, Nordrhein-Westfalen, DE

102, Institute for Community Medicine, University Medicine Greifswald, Greifswald, Mecklenburg-Vorpommern, DE

103, Department of Psychiatry, University of California, San Diego, San Diego, CA, US

104, KG Jebsen Centre for Psychosis Research, Norway Division of Mental Health and Addiction, Oslo University Hospital, Oslo, NO

105, Medical Genetics Section, CGEM, IGMM, University of Edinburgh, Edinburgh, GB

106, Clinical Neurosciences, University of Cambridge, Cambridge, GB

107, Internal Medicine, Erasmus MC, Rotterdam, Zuid-Holland, NL

108, Roche Pharmaceutical Research and Early Development, Neuroscience, Ophthalmology and Rare Diseases Discovery & Translational Medicine Area, Roche Innovation Center Basel, F. Hoffmann-La Roche Ltd, Basel, CH

109, Department of Psychiatry and Psychotherapy, University Medicine Greifswald, Greifswald, Mecklenburg-Vorpommern, DE

110, Department of Psychiatry, Leiden University Medical Center, Leiden, NL

111, Virginia Institute for Psychiatric & Behavioral Genetics, Virginia Commonwealth University, Richmond, VA, US

112, Computational Sciences Center of Emphasis, Pfizer Global Research and Development, Cambridge, MA, US

113, Institute for Molecular Bioscience; Queensland Brain Institute, The University of Queensland, Brisbane, QLD, AU

114, Department of Psychiatry, University of Münster, Münster, Nordrhein-Westfalen, DE

115, Department of Psychiatry, University of Münster, Münster, DE

116, Department of Psychiatry, Melbourne Medical School, University of Melbourne, Melbourne, AU

117, Florey Institute for Neuroscience and Mental Health, University of Melbourne, Melbourne, AU

118, Institute of Medical Genetics and Pathology, University Hospital Basel, University of Basel, Basel, CH

119, Institute of Neuroscience and Medicine (INM-1), Research Center Juelich, Juelich, DE

120, Amsterdam Public Health Institute, Vrije Universiteit Medical Center, Amsterdam, NL

121, Centre for Integrative Biology, Università degli Studi di Trento, Trento, Trentino-Alto Adige, IT

122, Department of Psychiatry and Psychotherapy, Medical Center - University of Freiburg, Faculty of Medicine, University of Freiburg, Freiburg, DE

123, Center for NeuroModulation, Faculty of Medicine, University of Freiburg, Freiburg, DE

124, Psychiatry, Kaiser Permanente Northern California, San Francisco, CA, US

125, Medical Research Council Human Genetics Unit, Institute of Genetics and Molecular Medicine, University of Edinburgh, Edinburgh, GB

126, Department of Psychiatry, University of Toronto, Toronto, ON, CA

127, Centre for Addiction and Mental Health, Toronto, ON, CA

128, Division of Psychiatry, University College London, London, GB

129, Neuroscience Therapeutic Area, Janssen Research and Development, LLC, Titusville, NJ, US

130, Institute of Molecular and Cell Biology, University of Tartu, Tartu, EE

131, Psychosis Research Unit, Aarhus University Hospital, Risskov, Aarhus, DK

132, Munich Cluster for Systems Neurology (SyNergy), Munich, DE

133, University of Liverpool, Liverpool, GB

134, Mental Health Center Copenhagen, Copenhagen Universtity Hospital, Copenhagen, DK

135, Human Genetics and Computational Biomedicine, Pfizer Global Research and Development, Groton, CT, US

136, Psychiatry, Harvard Medical School, Boston, MA, US

137, Psychiatry, University of Iowa, Iowa City, IA, US

138, Department of Psychiatry and Behavioral Sciences, Johns Hopkins University, Baltimore, MD, US

139, Department of Psychiatry and Psychotherapy, University Medical Center Göttingen, Goettingen, Niedersachsen, DE

140, Human Genetics Branch, NIMH Division of Intramural Research Programs, Bethesda, MD, US

141, Faculty of Medicine, University of Iceland, Reykjavik, IS

142, Child and Adolescent Psychiatry, Erasmus MC, Rotterdam, Zuid-Holland, NL

143, Psychiatry, Erasmus MC, Rotterdam, Zuid-Holland, NL

144, Psychiatry, Dalhousie University, Halifax, NS, CA

145, Division of Epidemiology, New York State Psychiatric Institute, New York, NY, US

146, Department of Clinical Medicine, University of Copenhagen, Copenhagen, DK

147, Department of Medical & Molecular Genetics, King's College London, London, GB

148, Psychiatry & Behavioral Sciences, Stanford University, Stanford, CA, US

149, NIHR Maudsley Biomedical Research Centre, King's College London, London, GB

150, Genetics, University of North Carolina at Chapel Hill, Chapel Hill, NC, US

151, Psychiatry, University of North Carolina at Chapel Hill, Chapel Hill, NC, US

**Supplementary methods**

**The testing samples – the discovery and replication sample**

The discovery sample included participants from the first data release (97% assessed at Cheadle, 3% at Newcastle) who were assessed in Cheadle, and included 10,674 individuals (age 45.9 - 80.3 years, mean=62.8, SD=7.4, 48.4% were men). The second release contained a higher proportion of individuals assessed in Newcastle (63% at Cheadle, 37% at Newcastle). The replication sample therefore included all participants included in the second release plus the smaller proportion (3%, N = 343) assessed at Newcastle in the first release. The replication sample consisted of 11,214 individuals in total (age 46.5 - 80.8 years, mean age=64.4, SD=7.4, and 49.4% were men, see Figure S1). Comparisons between the Cheadle and Newcastle sites are reported in Figures S10-11 and Tables S7-9.

**Meta-analysis of GWAS on depression and the testing samples**

Methods for the meta-analysis has been described in another paper by Howard et al. (2018)^1^. The only difference between the present paper and the paper cited above is the sample from UK Biobank. In the original meta-analysis paper, 371,435 unrelated, European-ancestry participants who have not participated in the PGC and 23andMe GWAS were included after genotyping quality check. The present paper further removed 22,404 people who are unrelated, have British ancestry, and those who attended imaging assessment, if there is a non-empty entry of ‘date of attending assessment centre’ in field 53.2.0 (<http://biobank.ctsu.ox.ac.uk/crystal/field.cgi?id=53>). For the testing sample, which includes the discovery and replication samples, 516 participants were further removed for either having NA value reported in the MRI site field (f.54.2.0, <http://biobank.ctsu.ox.ac.uk/crystal/field.cgi?id=54>) or was recruited in Reading (code: 11026, <http://biobank.ctsu.ox.ac.uk/crystal/coding.cgi?id=10>), from which imaging-derived phenotypes were not yet available. This eventually leaves 21,888 people for the analyses conducted in the present study. GWAS was conducted on the subset of UK Biobank sample where people that have attended imaging assessment were excluded, using the BGENIE pipeline v1.1^2^. Statistic models and genotyping quality control remained the same as a depression GWAS conducted on UK Biobank^3^. Meta-analysis was conducted using ‘Metal’^4^, with the same procedures described elsewhere^1^.

**Generating depression-PGRS for the testing samples**

DNA extraction and genotyping were described in an early protocol paper^5^ and in the UK Biobank protocol documentation (<http://www.ukbiobank.ac.uk/wp-content/uploads/2014/04/UKBiobank_genotyping_QC_documentation-web.pdf>). In brief, sample collection was conducted by the UK Biobank team, and genotyping was conducted at the Affymetrix Research Services. SNP quality-control was conducted by the UK Biobank team based on the criterion of heterogeneous genotype frequencies in the same samples and across genotyping arrays, Hardy-Weinberg disequilibrium (p<10^-5^), low minor allele frequency (<0.01), low imputation accuracy (<0.1) and low call rate (<95%). In addition to these steps, we further conducted sample control, removing participants who have missingness of >95%, have gender-mismatched genetic data, have non-European-ancestry and those who are related. Non-European ancestry was identified based on k-means clustering on the genetic principal components to identify white British ancestry^6^ and those who declared non-white British ancestry were further removed. Relatedness was quantified by the kinship coefficient using the King’s criteria^7^. First-degree relatives were identified as one family, and one of the participants in each family was randomly selected to add back into the sample to maximise sample size while ensuring relatedness removed.

Genotype data after quality check was then fed into PRSice 2.0 program. The clumping threshold was set as p=1, LD score r^2^=0.25, distance threshold=250 Kb. P thresholds for scoring were set at p<0.0005, p<0.001, p<0.005, p<0.01, p<0.05, p<0.1, p<0.5 and p<1. No phenotype data was fed into the program except for a dummy phenotype file that covers all the participants for the testing sample.

**Supplementary information for behavioural phenotypes**

Principal component analysis (PCA) was conducted on cognitive variables (Table 1 and S1) to extract a measure of g of cognitive abilities. Tasks conducted at the imaging assessment centres were used for this step because these measures cover much more people than online questionnaires (N~=20,000 for the former, and N~=11,000 for the latter). Variance explained by the first latent variable was 30.2%.

**Neuroimaging phenotypes in UK Biobank**

T1 data was processed to estimate intracranial and subcortical volumes. First, total volumes for white matter, grey matter and peripheral cerebrospinal fluid were calculated, and the sum of the three was the derived intracranial volume. This derived intracranial volume is highly correlated with the field “Volumetric scaling from T1 head image to standard space” (f.25000.2.0) with a correlation coefficient of -0.898. Then volumes for thalamus, caudate, putamen, pallidum, hippocampus, amygdala, accumbens and brain stem (with 4^th^ ventricle) were estimated (see Figure S2).

DTI data pre-processing included correction for eddy currents and head motion, outlier-slices correction and grand distortion correction. Maps for fractional anisotropy (FA) and mean diffusivity (MD) were generated and FA maps were used to generate tract masks, using probabilistic tractography analysis by AutoPtx package from FSL^8^. Twenty-seven tracts were generated (12 bilateral and 3 unilateral tracts, see Supplementary Figure S3 and Table S1)^9^. Weighted mean FA and MD were then calculated for each tract. To determine general variances in DTI measures and main subsets, as have validated in previous papers that weighted mean DTI measures for major white matter tracts are highly correlated, which makes generating general variances possible^10,11^, we performed PCA on (1) FA/MD of all 27 tracts (gTotal), (2) FA/MD on association/commissural fibres (gAF), which connect the prefrontal cortex to other cortexes, (3) FA/MD on thalamic radiations (gTR), consisted of tracts that link the thalamus to the cortex, and (4) FA/MD on projection fibres (gPF), locating within brain stem or spinal cord or link them to the cortex (for anatomy of these three subsets, see Figure S3). Variance explained by the first latent component can be found in Figure S22, and reports of correlation loadings of each variable on the latent factors are in Table S17. The above PCA was conducted on the total sample of both releases to achieve more accurate estimations. The scores for the first unrotated principal component were used as the indices for general variants of total variance and variances in three major subsets. In order to control for the effects driven by outliers, subjects with a gTotal for FA/MD outside of +/-3 standard deviation from mean were excluded^10^.

Resting-state data was pre-processed through FSL-style motion correction, grand-mean intensity normalisation, high-pass temporal filtering, EPI unwarping and grand-distortion-correction unwarping. A group-level independent component analysis was conducted on the first 4,100 people to reduce data dimension^12^. The brain was therefore parcellated into 25 independent components, and 21 of them were left for further analyses after 4 discarded as being identified manually as noise components. The time-series data for nodes was then used to calculate functional connectivity between node pairs. It was achieved by estimating partial Pearson correlation with an L2 regularisation applied (rho set as 0.5 in FSLNets). All r-scores were then Fisher-transformed into z-scores. This resulted in a 21*21 correlation matrix of functional connectivity for each participant. In order to aid comprehension, all connectivity values were transformed into absolute strength by multiplying the sign of group-mean value for each of the connection^13^. For amplitude of signal fluctuation, as the data has had high-pass temporal filtering, therefore the measure mainly represent temporal fluctuations of blood oxygen-level dependent signal^14^. An interactive website displaying group-mean maps for each node can be found in a URL: <http://www.fmrib.ox.ac.uk/datasets/ukbiobank/group_means/rfMRI_ICA_d25_good_nodes.html>. A connectome map of the nodes can be found in:
<http://www.fmrib.ox.ac.uk/datasets/ukbiobank/netjs_d25/>.

**References**

1.	Howard, D. M. *et al.* Genome-wide meta-analysis of depression in 807,553 individuals identifies 102 independent variants with replication in a further 1,507,153 individuals. *bioRxiv* **6288,** 433367 (2018).

2.	Bycroft, C. *et al.* Genome-wide genetic data on ~500,000 UK Biobank participants. *bioRxiv* 1–36 (2017).

3.	Howard, D. M. *et al.* Genome-wide association study of depression phenotypes in UK Biobank identifies variants in excitatory synaptic pathways. *Nat. Commun.* **9,** 1–10 (2018).

4.	Willer, C. J., Li, Y., Abecasis, G. R. & Overall, P. METAL: fast and efficient meta-analysis of genomewide association scans. *Bioinformatics* **26,** 2190–2191 (2010).

5.	Bycroft, C. *et al.* Genome-wide genetic data on 500,000 UK Biobank participants. *bioRxiv* 166298 (2017). doi:10.1101/166298

6.	Warren, H. R. *et al.* Genome-wide association analysis identifies novel blood pressure loci and offers biological insights into cardiovascular risk. *Nat. Genet.* **49,** 403–415 (2017).

7.	Manichaikul, A. *et al.* Robust relationship inference in genome-wide association studies. *Bioinformatics* **26,** 2867–2873 (2010).

8.	Mori, S. *et al.* Imaging cortical association tracts in the human brain using diffusion-tensor-based axonal tracking. *Magn. Reson. Med.* **47,** 215–223 (2002).

9.	Wakana, S., Jiang, H., Nagae-Poetscher, L. M., van Zijl, P. C. M. & Mori, S. Fiber tract-based atlas of human white matter anatomy. *Radiology* **230,** 77–87 (2004).

10.	Shen, X. *et al.* Subcortical volume and white matter integrity abnormalities in major depressive disorder: Findings from UK Biobank imaging data. *Sci. Rep.* **7,** 1–10 (2017).

11.	Cox, S. R. *et al.* Ageing and brain white matter structure in 3,513 UK Biobank participants. *Nat. Commun.* **7,** 1–34 (2016).

12.	Alfaro-Almagro, F. *et al.* Image processing and Quality Control for the first 10,000 brain imaging datasets from UK Biobank. *Neuroimage* **166,** 400–424 (2018).

13.	Shen, X. *et al.* Resting-state connectivity and its association with cognitive performance, educational attainment, and household income in UK Biobank. *Biol. Psychiatry Cogn. Neurosci. Neuroimaging* 1–9 (2018). doi:10.1016/J.BPSC.2018.06.007

14.	Bijsterbosch, J. *et al.* Investigations into within- and between-subject resting-state amplitude variations. *Neuroimage* **159,** 57–69 (2017).

Figure S1. Illustration of the discovery and replication samples. For the release in May 2018, 10,674 people attended the site in Cheadle, and 343 people participated in New Castle. For the other release in Oct 2018, 6,850 people attended the site in Cheadle, and 4,021 people participated in New Castle. For data collected in Cheadle, there were 17,524 people in total (49.3% were men, mean age = 63.38 years, SD of age = 7.49 years). There were 4,364 people assessed at Newcastle (47.3% were men, mean age = 64.49 years, SD of age = 7.36 years). Combining the samples from Cheadle and Newcastle, there were 21,888 participants in total (48.9% were men, mean age = 63.66 years, SD of age = 7.48 years).

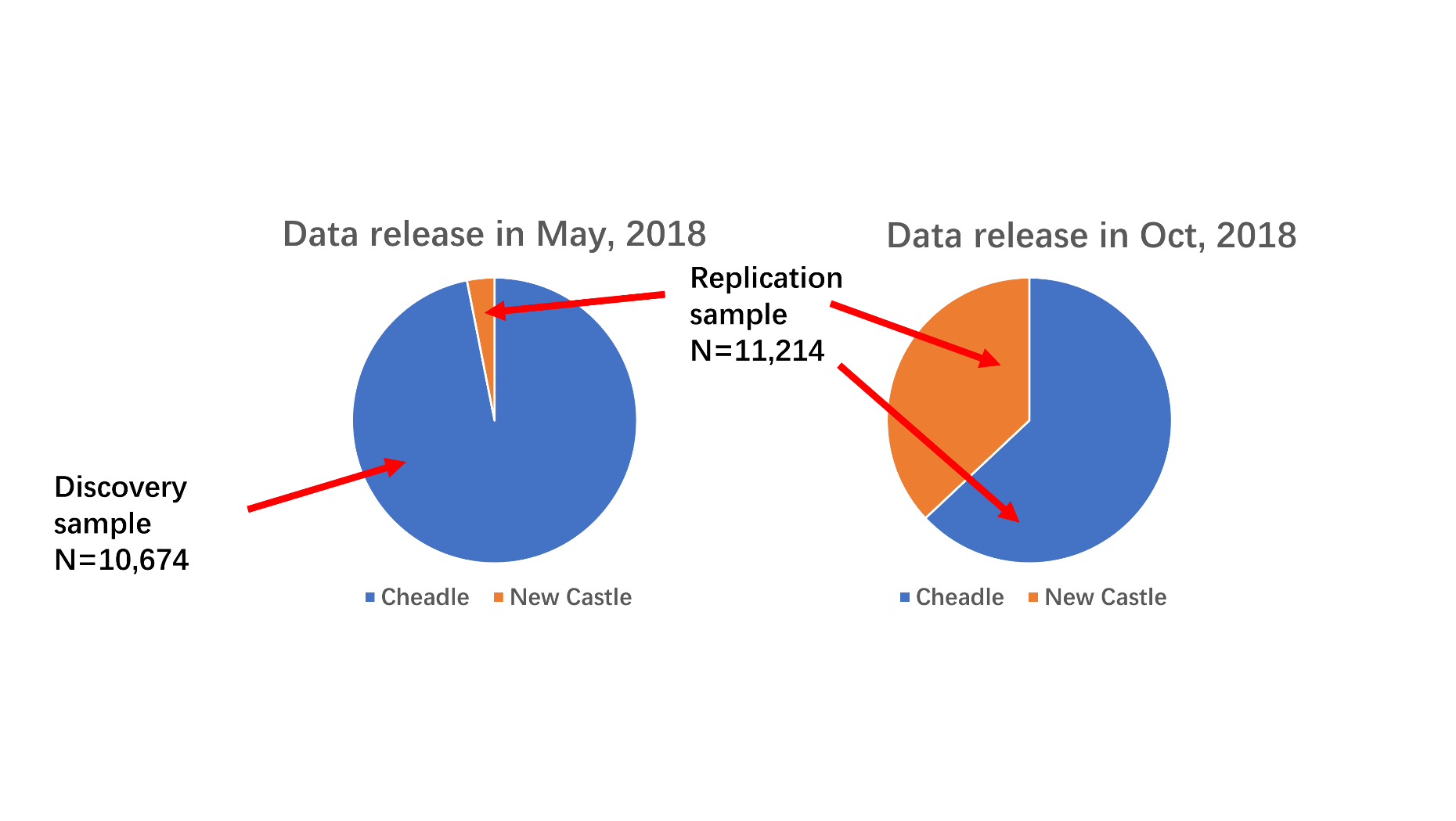

Figure S2. Visualisation of subcortical structures. Brain maps were shown in anterior, inferior and right views (from left to right). Masks for the structures were the group-mean data acquired at the UK Biobank data showcase site (source 9028).

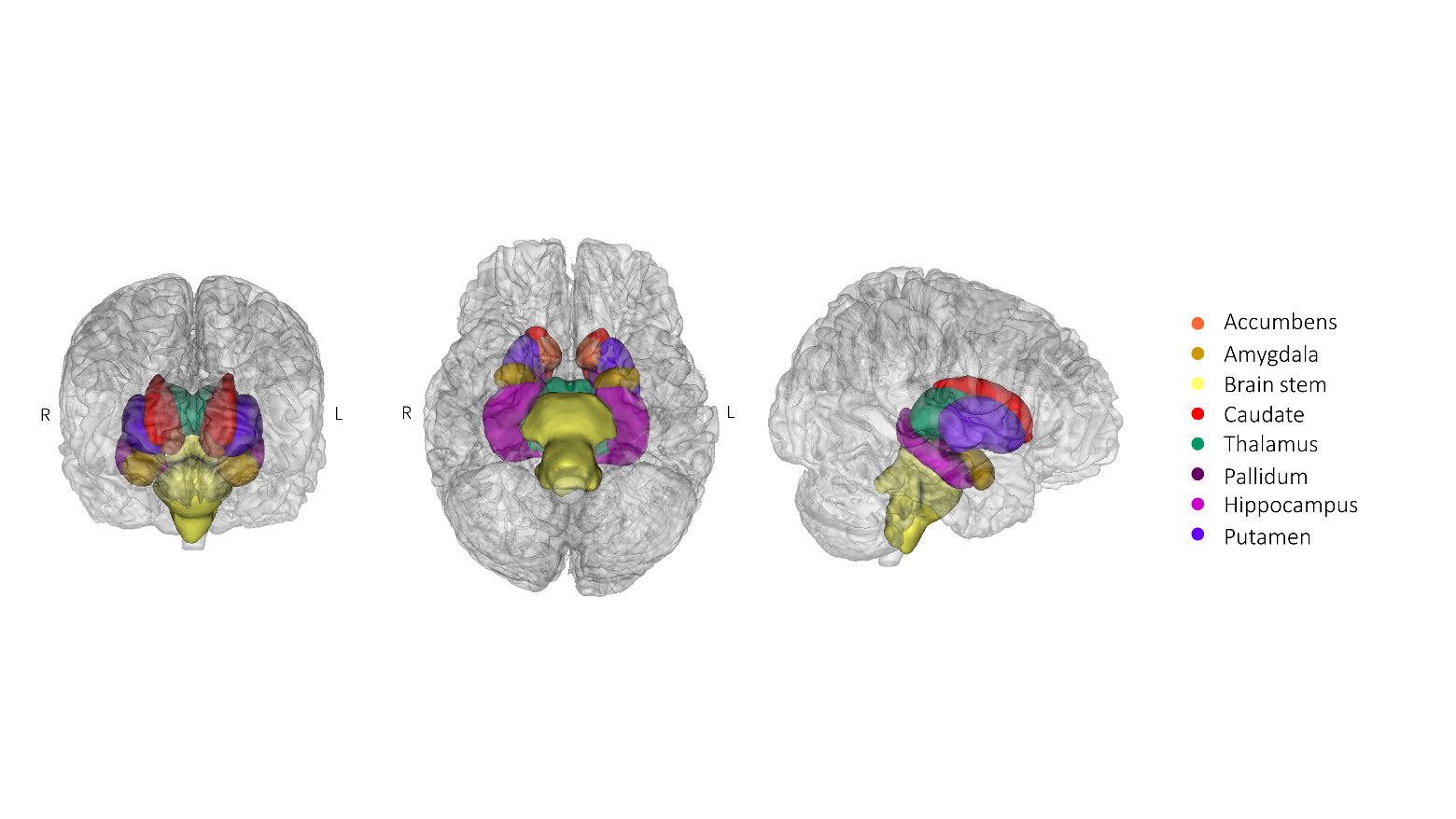

Figure S3. Visualisation of all individual white matter tracts. White matter tracts are presented in three subsets. From left to right, brain maps are presented in the superior, anterior and right views. Masks for the structures were the group-mean data acquired at the UK Biobank data showcase site (source 9028). To achieve better results of visualisation, a threshold at >30% of the highest intensity was applied to filter out weaker regions.

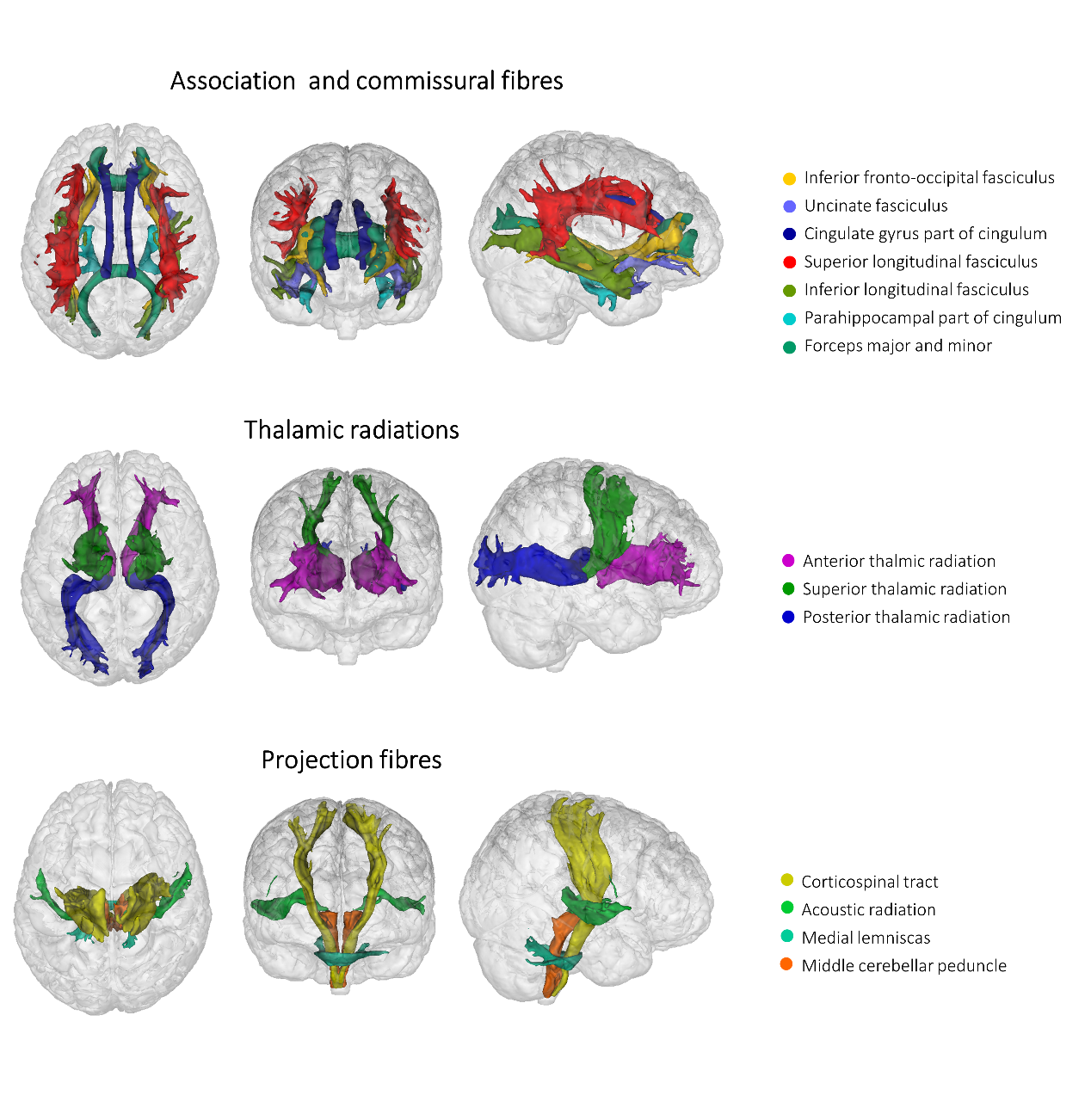

Figure S4. Illustration of resting-state connectivity between 21 ICA components. Thickness and the shade of the lines connecting the nodes represent the Fisher-transformed z score of partial correlation. Other than functional connectivity, the low-frequency amplitude of the components/nodes below were estimated for analysis. 3-D brain maps for the components/nodes can be found in: <http://www.fmrib.ox.ac.uk/datasets/ukbiobank/group_means/rfMRI_ICA_d25_good_nodes.html>.

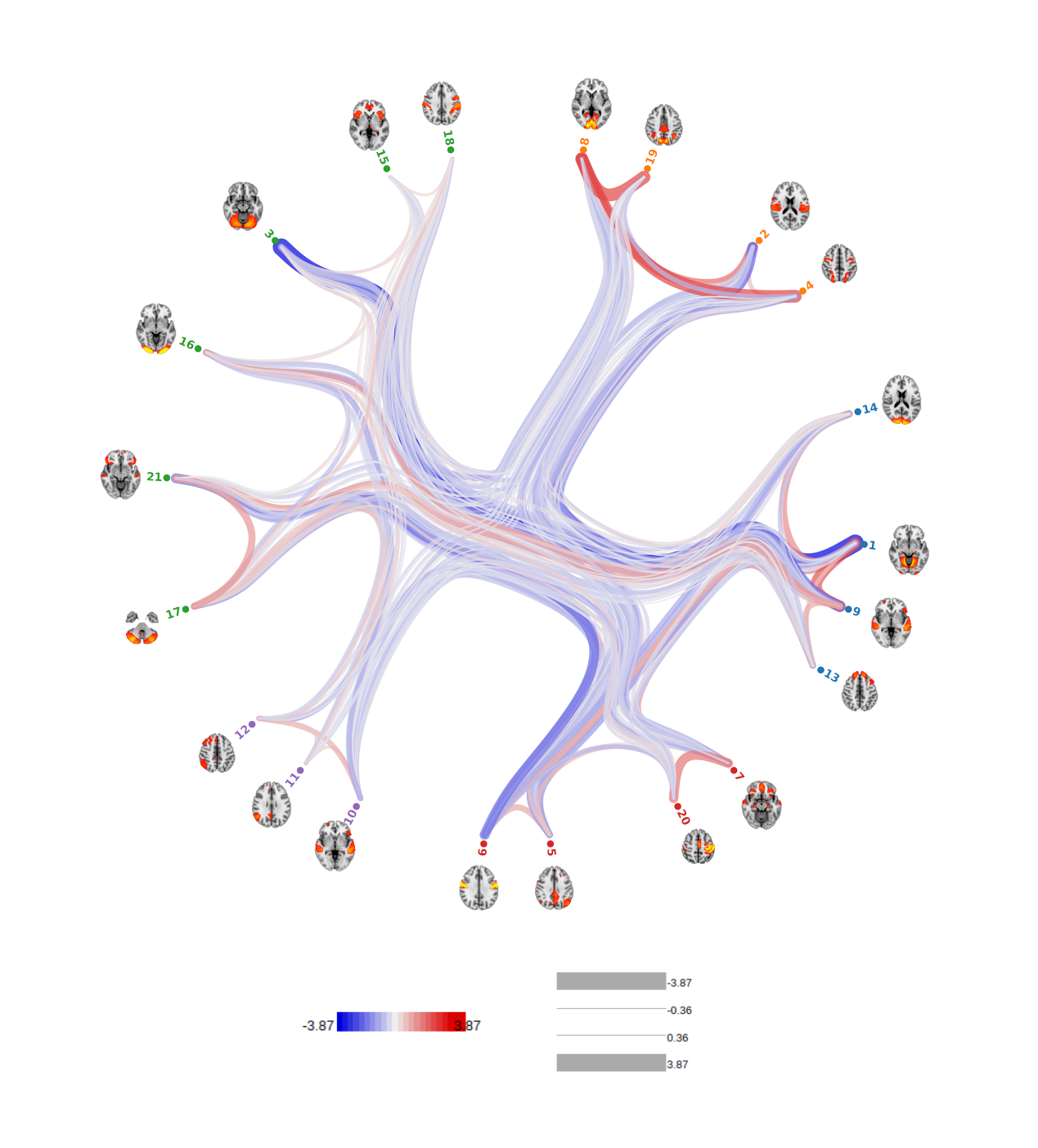

Figure S5. Flow-chart for the procedures of Mendelian Randomisation analyses. In the chart, from left to right, the main steps are listed by sequence (the colour-filled labels). Within each major step, the procedures are listed underneath the main labels (as the boxes with colours consistent with the main labels).

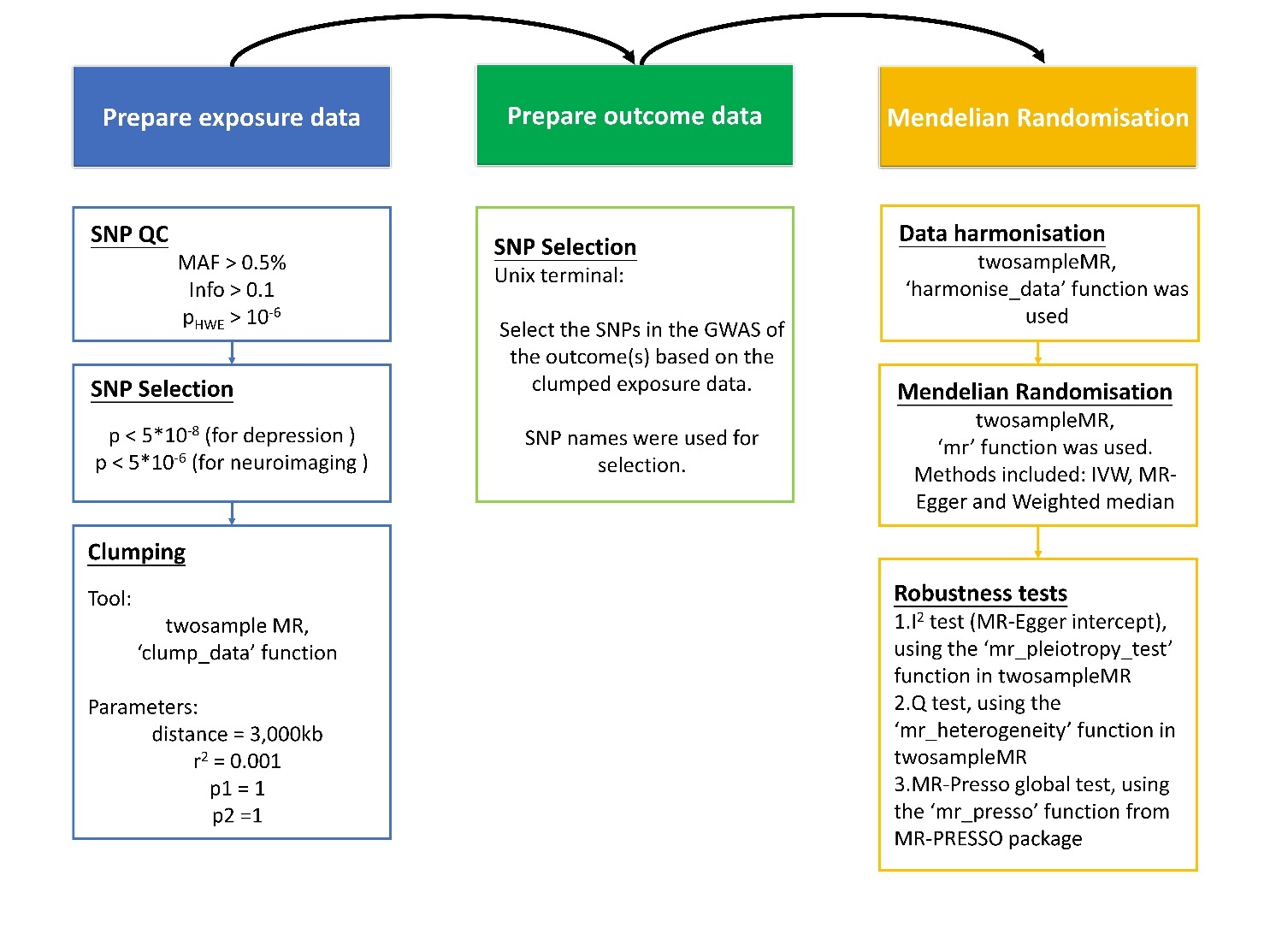

Figure S6. Two types of mediation models. (A) Mediation model testing the mediating effect of manifestations of depression for the association between genetic risk and neuroimaging variables. (B) Mediation model testing the mediating effect of neuroimaging variables for the genetic association of depression. Neuroimaging variables were selected based on the results from Mendelian Randomisation analyses. Those that showed as causal consequences of depression were tested in model (A) and the ones showed significant causal effect to depression were tested in model (B). Variables that were considered as manifestations of depression include CIDI depression, severity of depression assessed by CIDI short form and current depressive symptoms assessed by PHQ-4 at the imaging assessment. The results for depression-PGRS is reported in the main text.

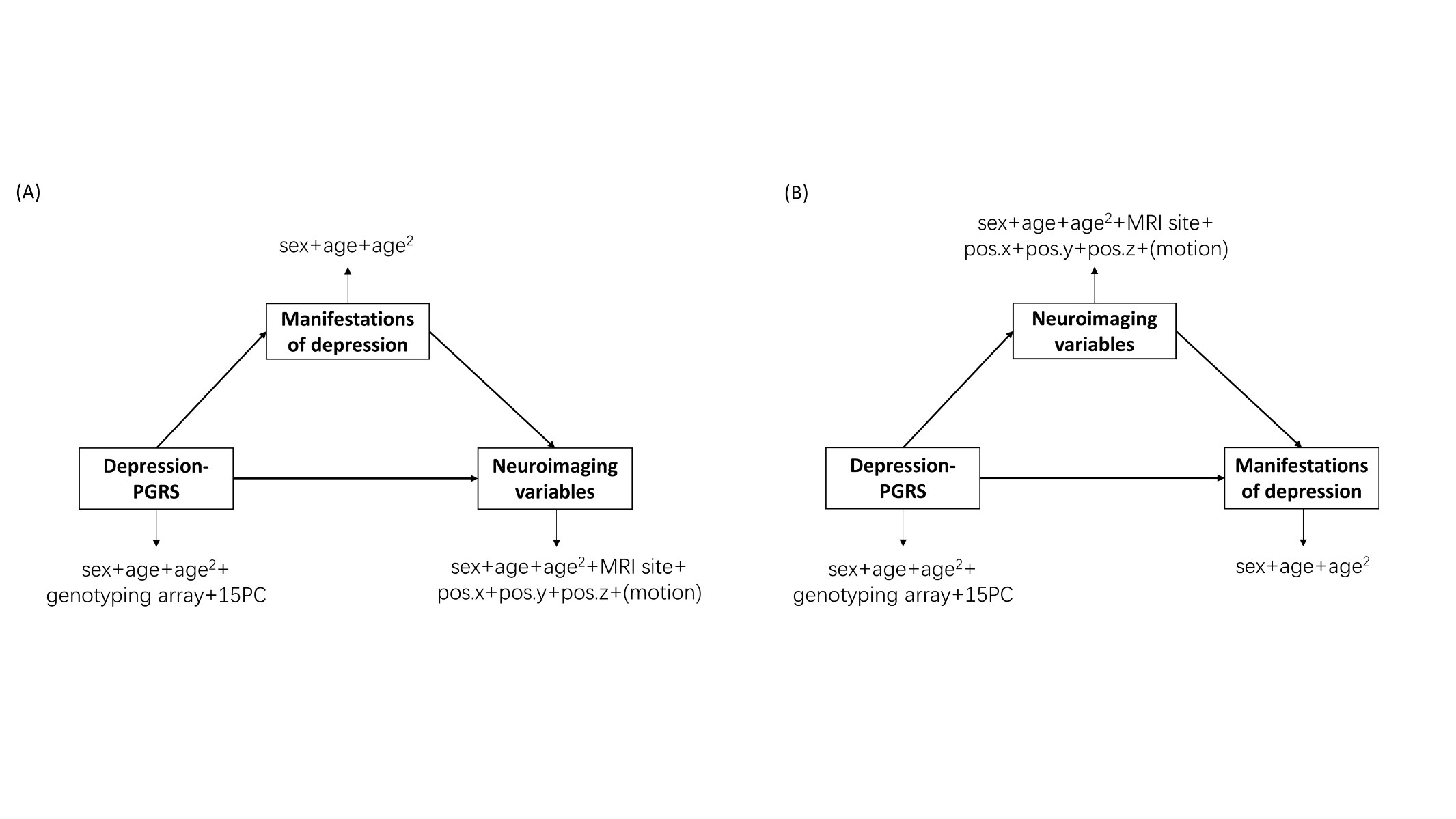

Figure S7. Significance plots for PheWAS results on all depression-PGRS p thresholds (pT). Panels (A-D) are shown on this page and the other panels (E-H) are shown on the next page. Top associations found in depression-PGRS at pT<1 and pT<0.01 (see the main text) are annotated in each panel.

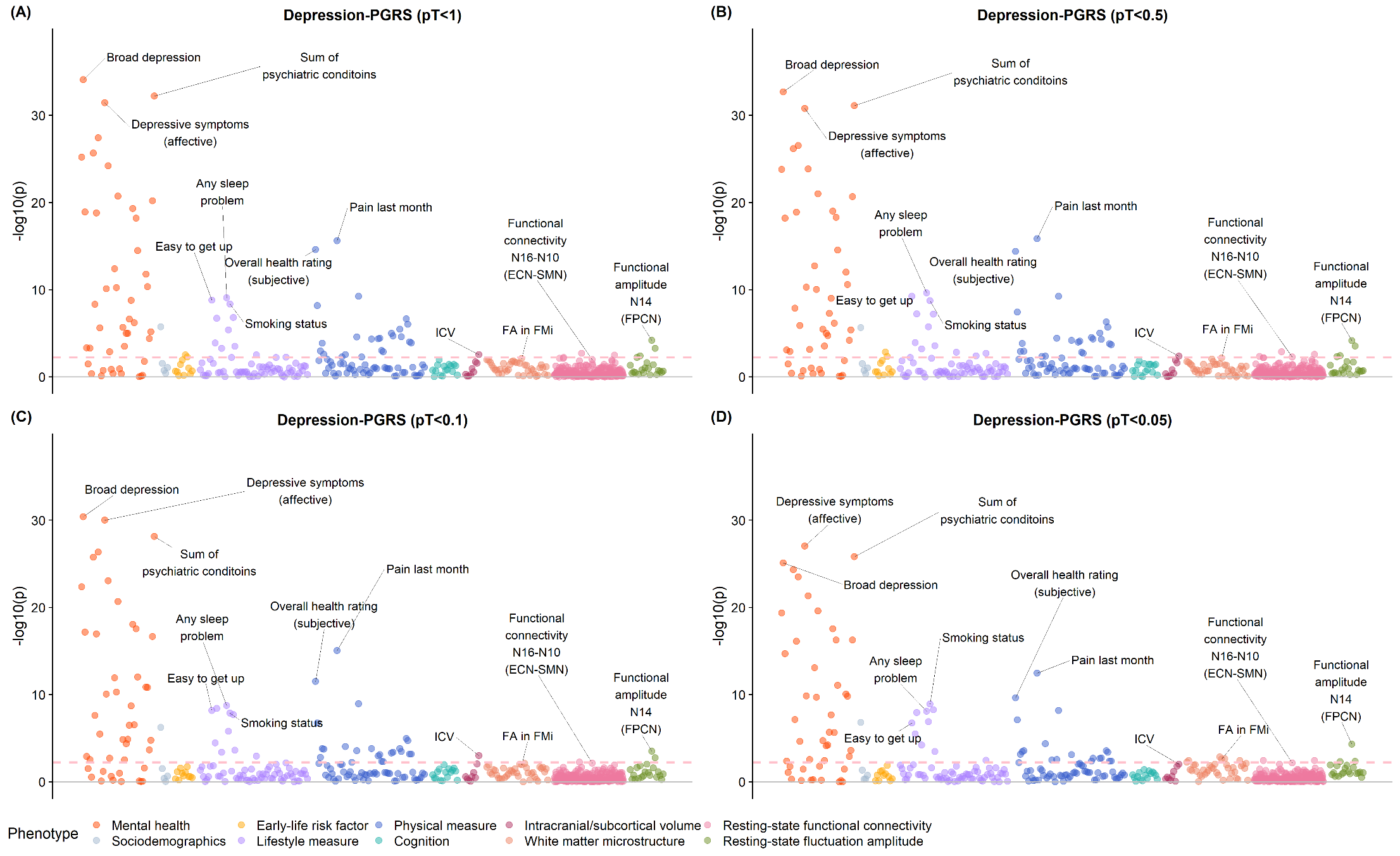

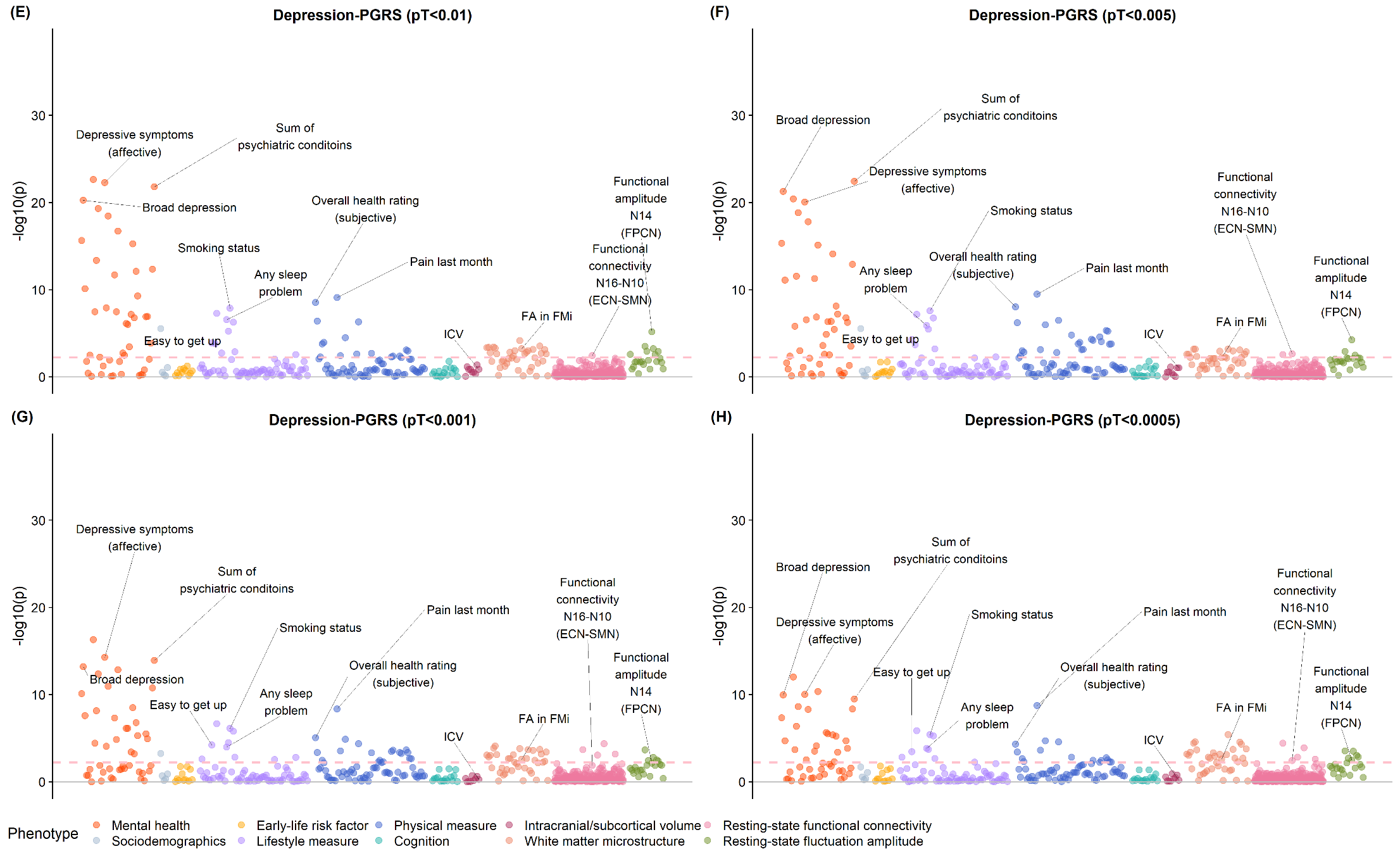

Figure S8. Heatmap for results from the replication sample. Traits were selected based on whether they were associated with depression-PGRS at four p thresholds minimal in the discovery sample.

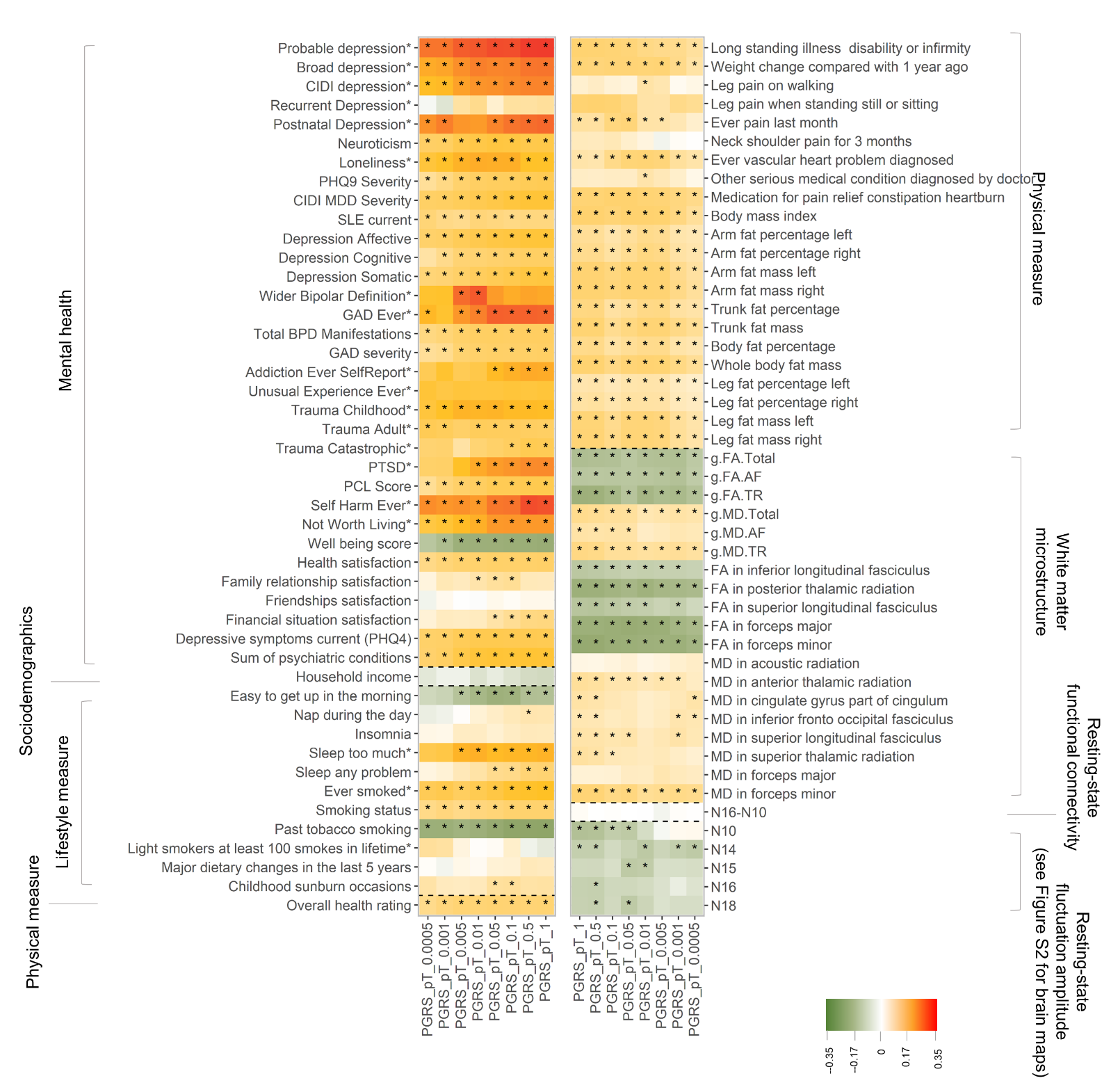

Figure S9. Significance maps for replication analysis on all depression-PGRS on eight pT. In the replication analysis, traits that were significantly associated with depression-PGRS at minimal two pT were selected. The selected traits were tested in the replication sample at all eight pT. These tests were then FDR-corrected altogether (see methods). Top associations found in the discovery sample were annotated in each panel. Although the p-value maps for depression-PGRS pT<1 and pT<0.01 are presented in Figure 3, here for the purpose of completeness, figures for these two pT are presented again below. The red dash lines in the panels represent the nominally significance threshold (p=0.05) and the yellow dash lines represent the FDR-corrected significance threshold (q_FDR_=0.05).

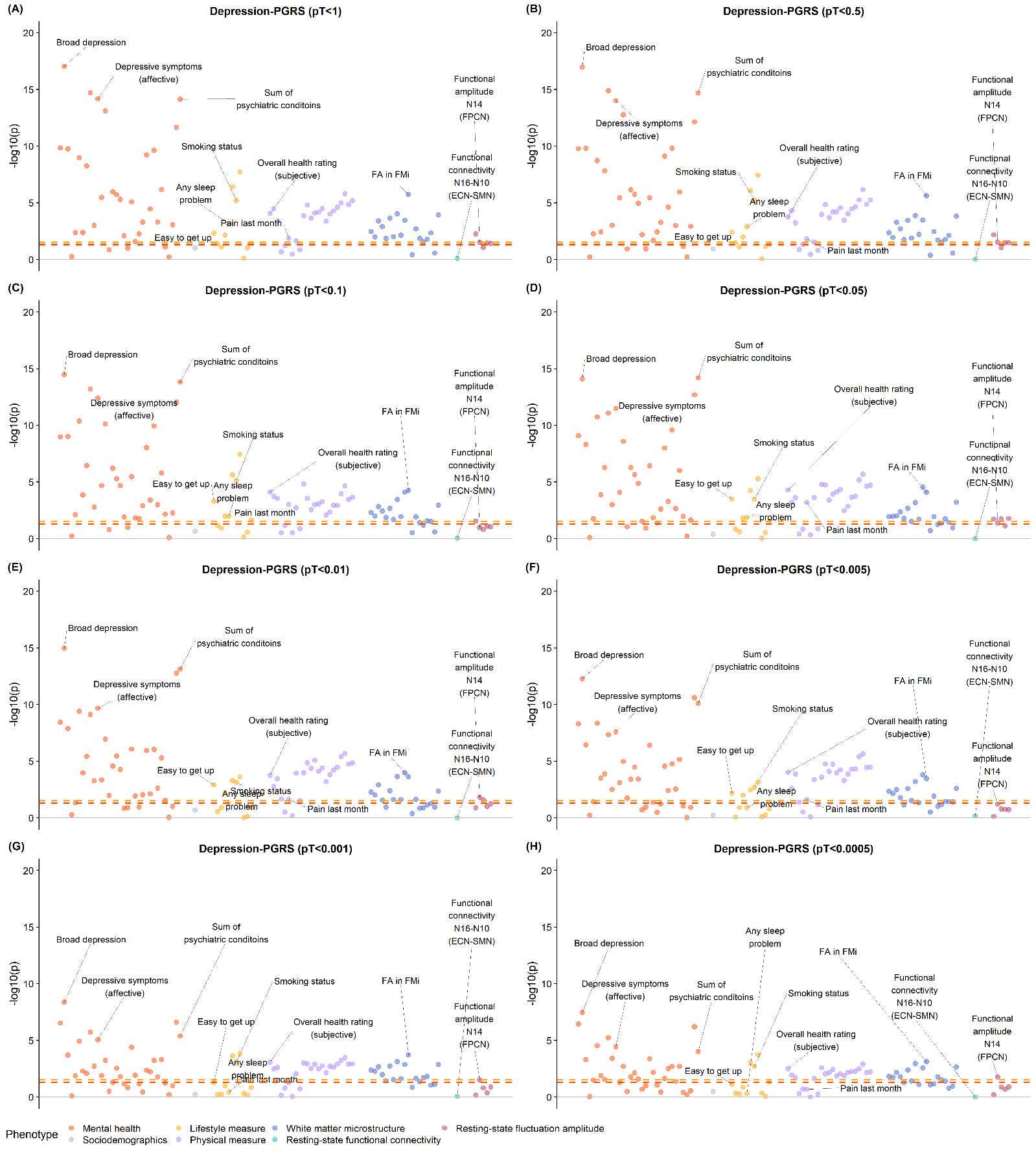

Figure S10. Comparison of the effect sizes between the datasets acquired in the Cheadle and Newcastle MRI sites. Traits selected were the ones that were associated with depression-PGRS at four p thresholds minimal in Cheadle. List of traits can be found in Table S5.

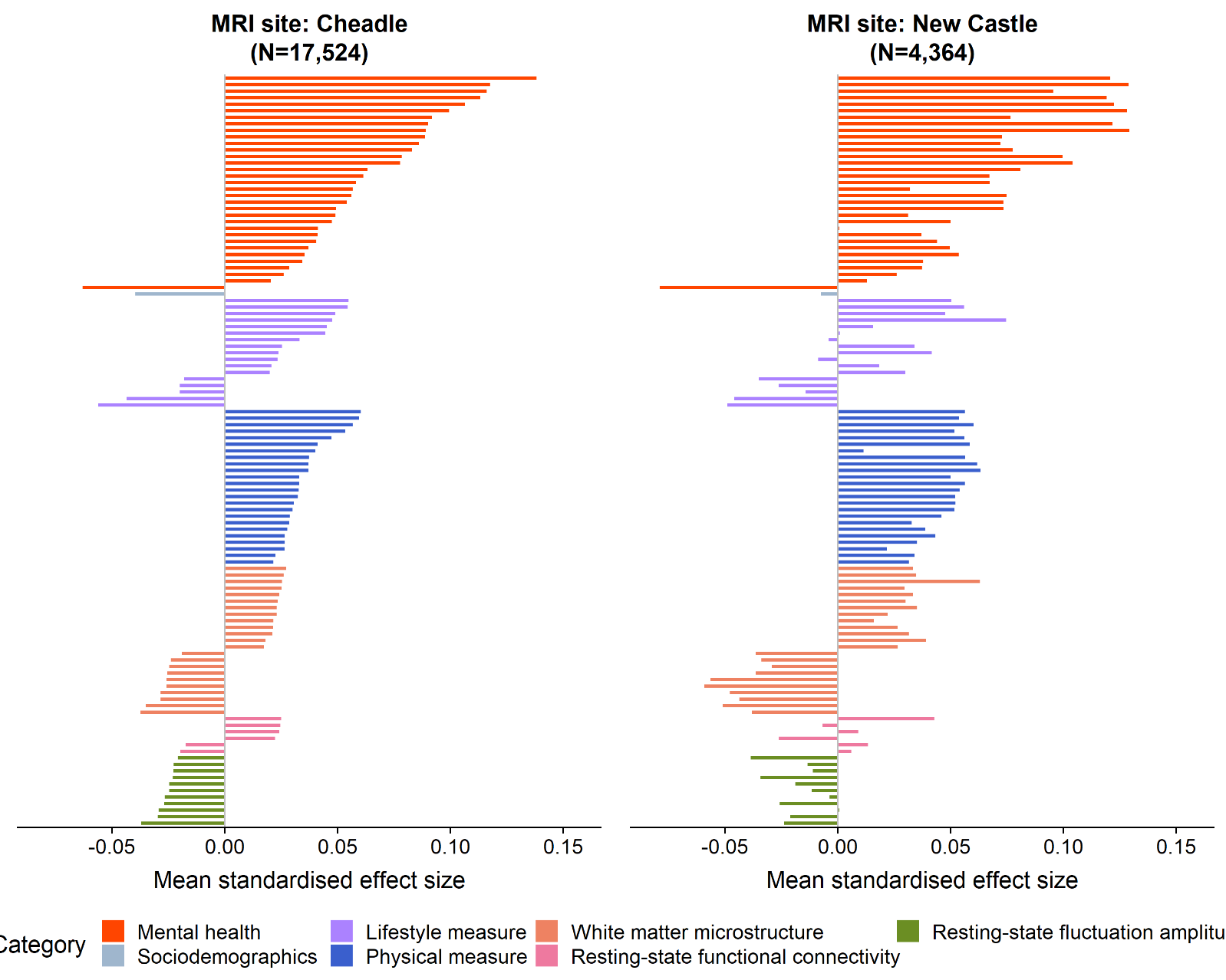

Figure S11. Significance maps for the interaction term of MRI site × depression-PGRS for the traits that were found associated with depression-PGRS at minimal four pT.

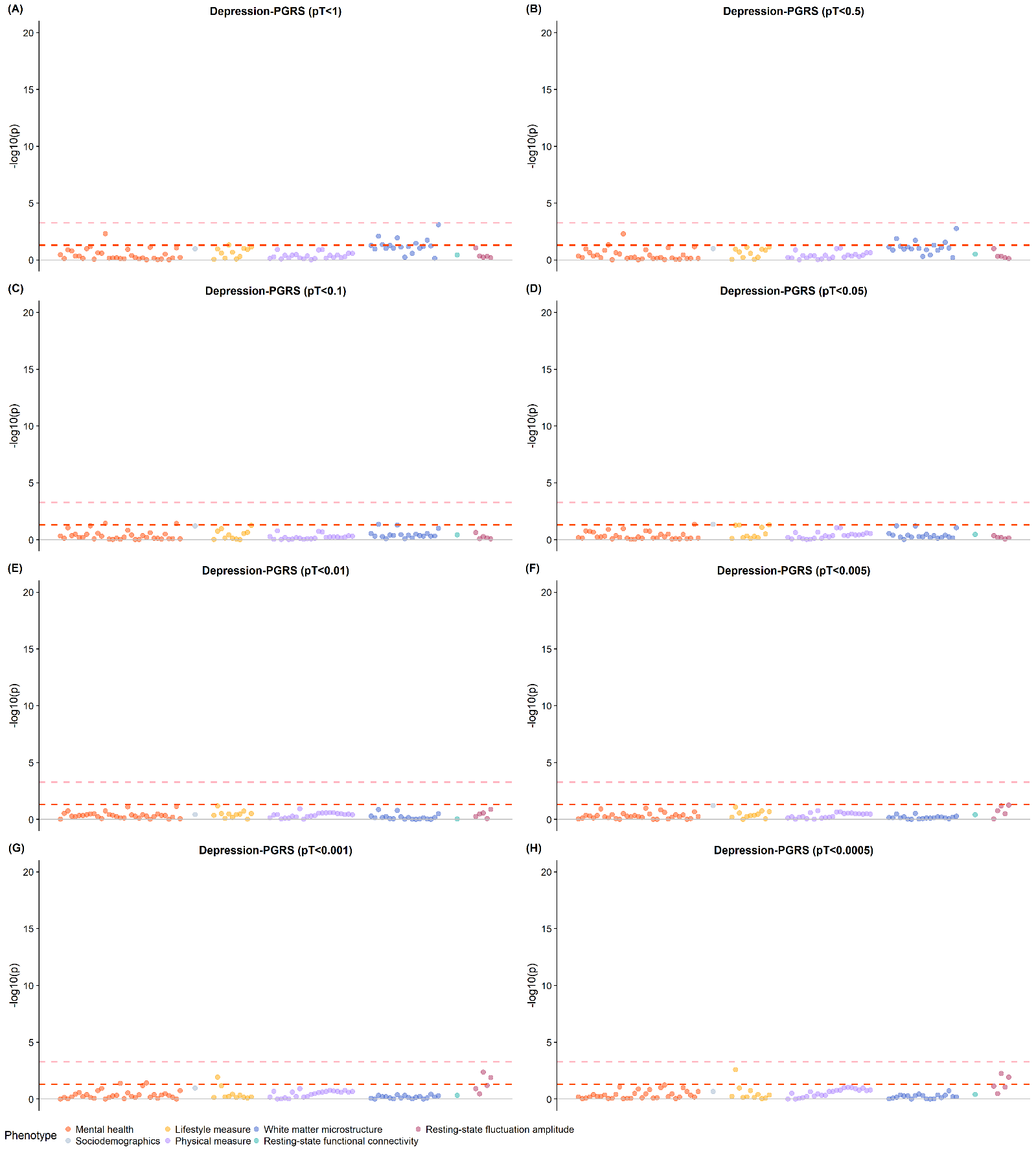

Figure S12. Significance plots for PheWAS results on all depression-PGRS p thresholds (pT) in the total sample including both the discovery and the replication samples (N=21,888). Panels (A-D) are shown on this page and the other panels (E-H) are shown on the next page. Top associations found in depression-PGRS at pT<1 and pT<0.01 (see the main text) are annotated in each panel.

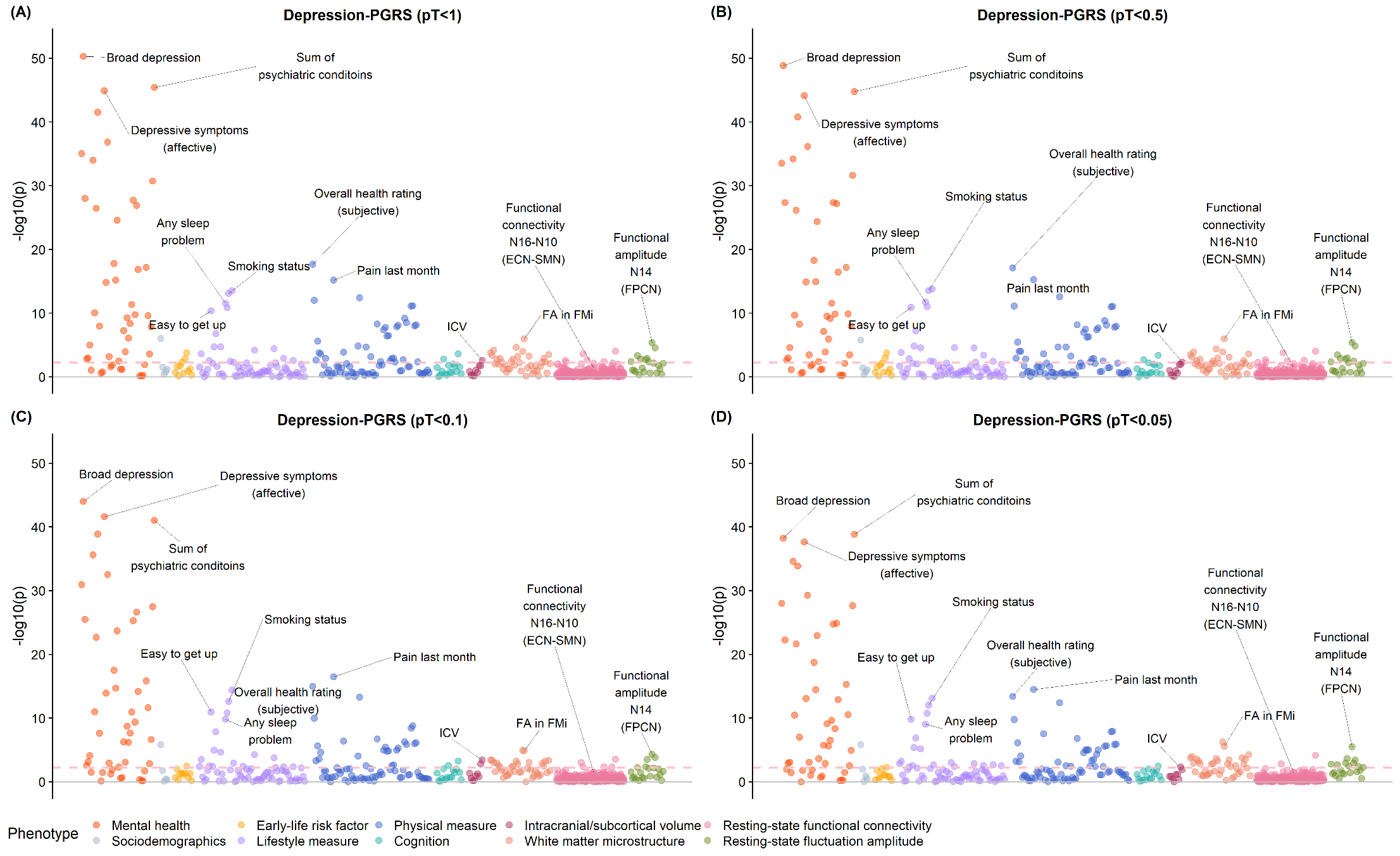

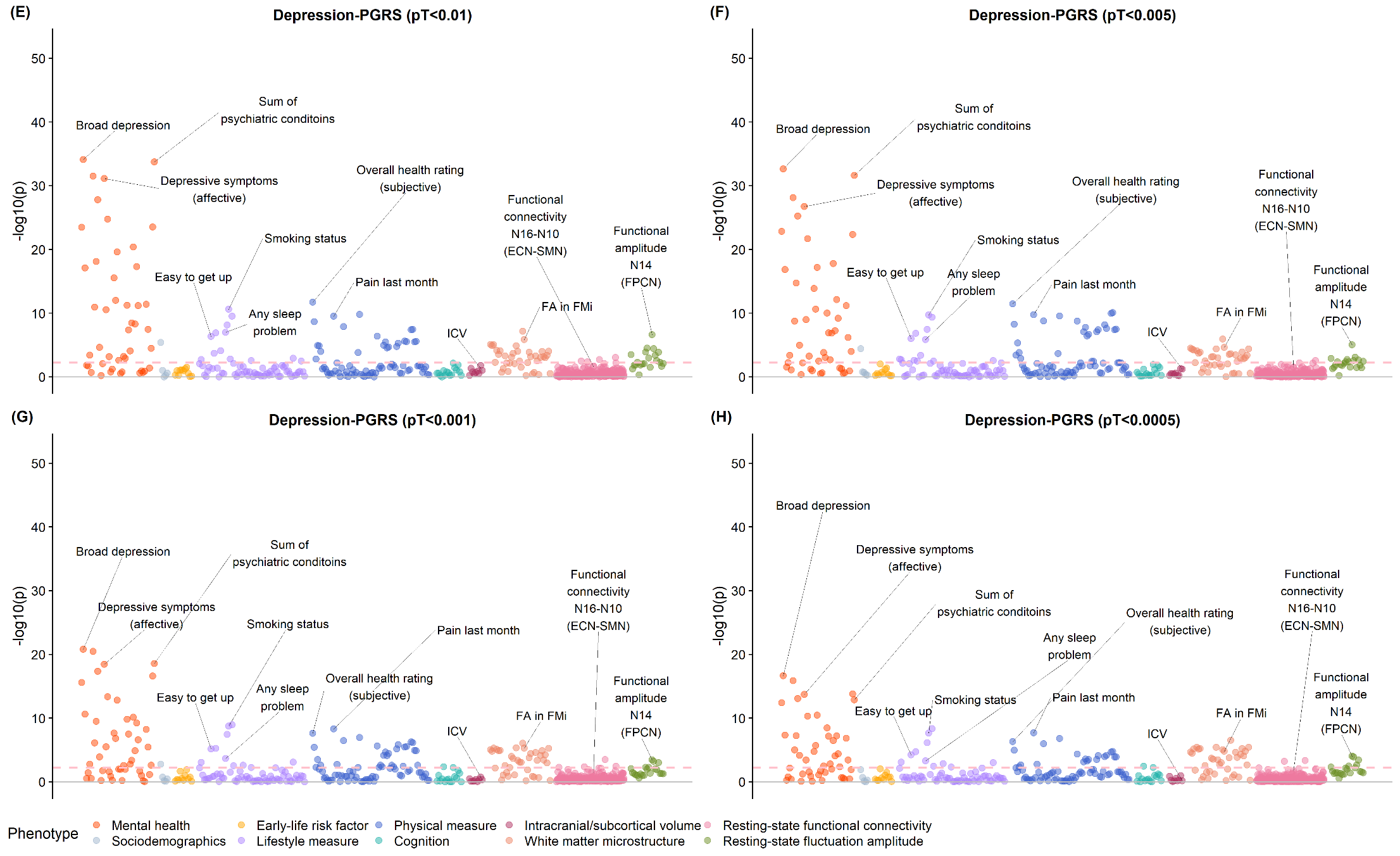

Figure S13. Heatmap for the traits that associated with depression-PGRS at four minimum four pT in the total sample including both the discovery and the replication samples (N=21,888).

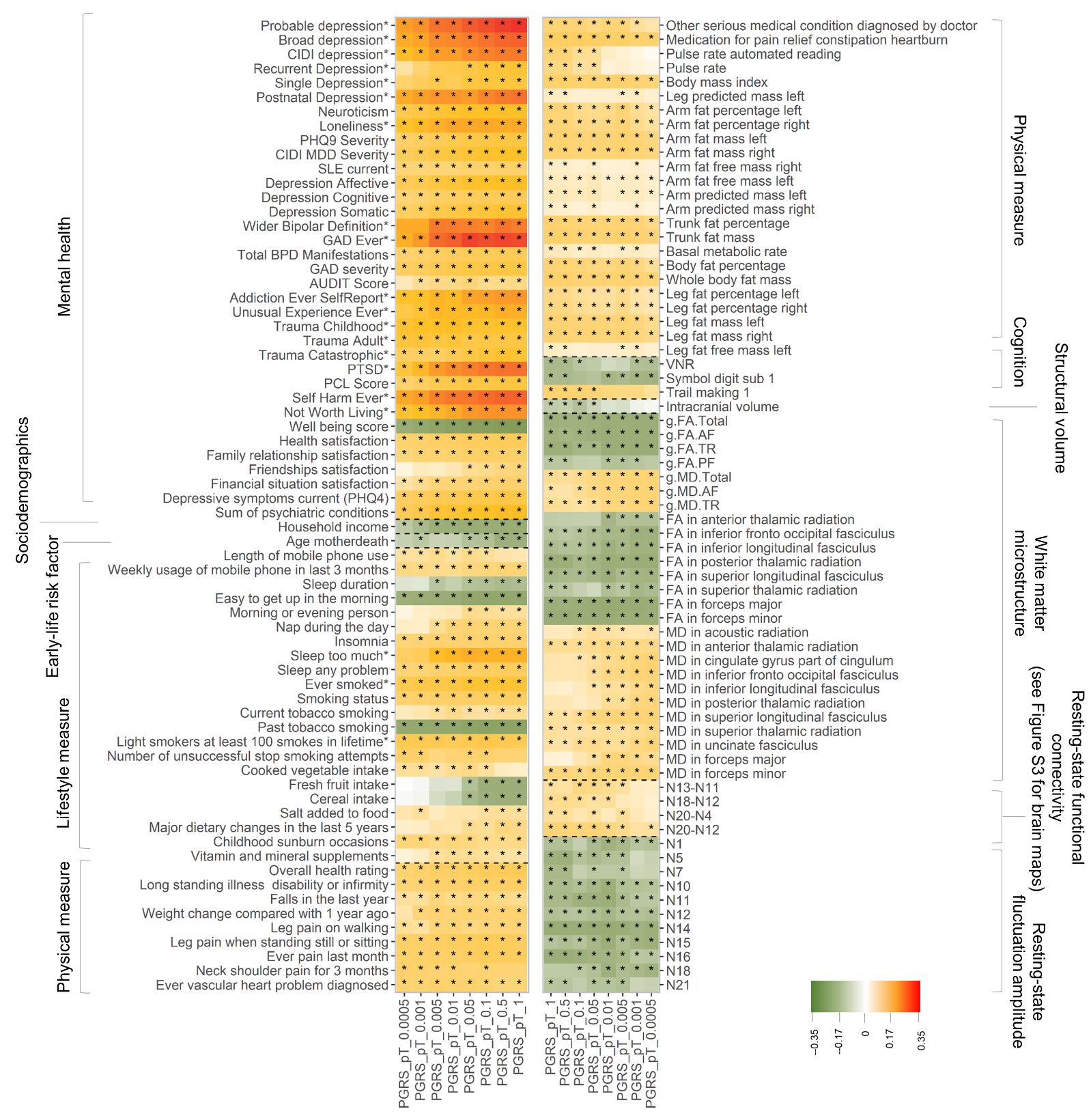

Figure S14. Mendelian Randomisation analysis testing the causal effect of depression on gMD-Total.

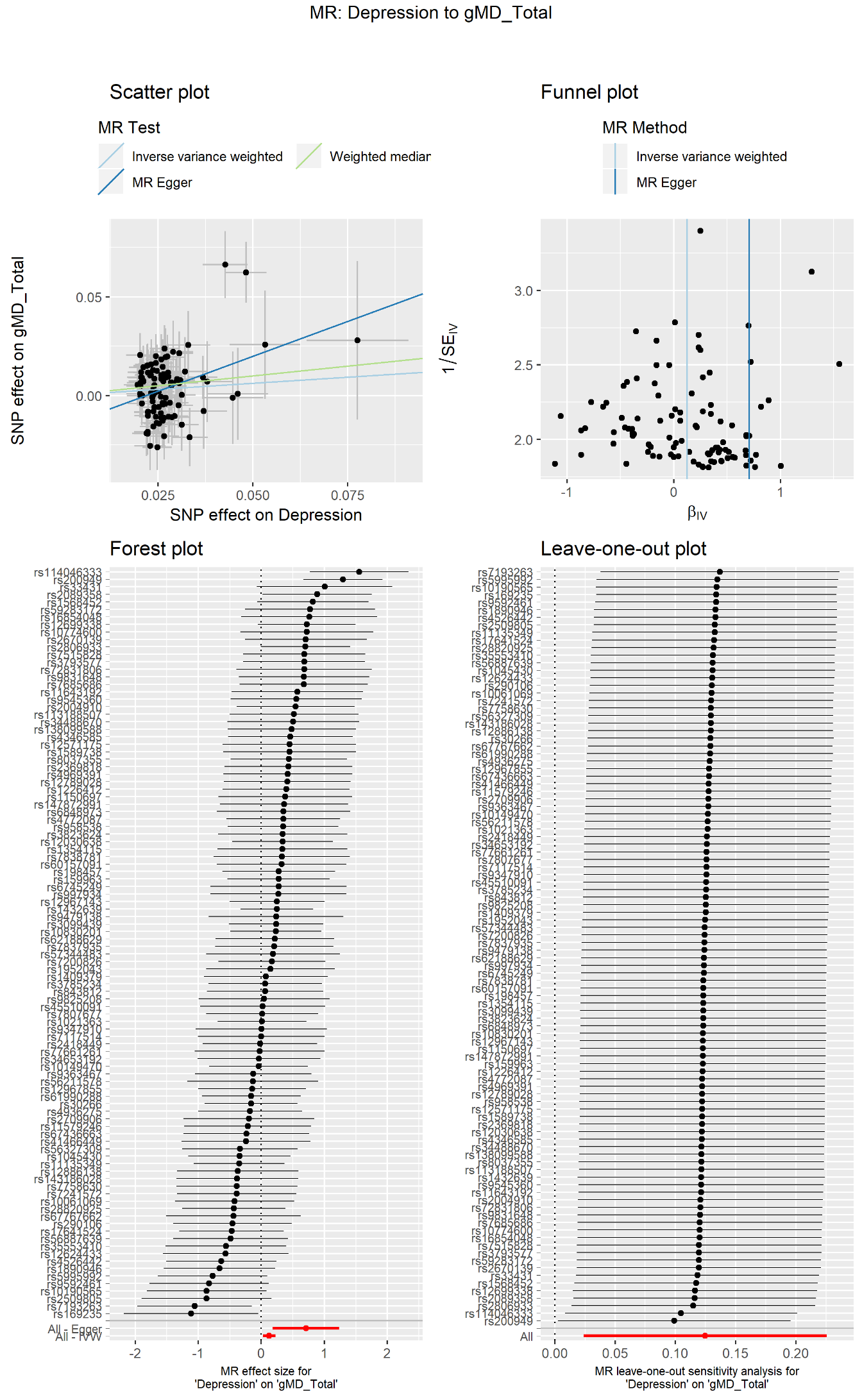

Figure S15. Mendelian Randomisation analysis testing the causal effect of depression on gMD-TR.

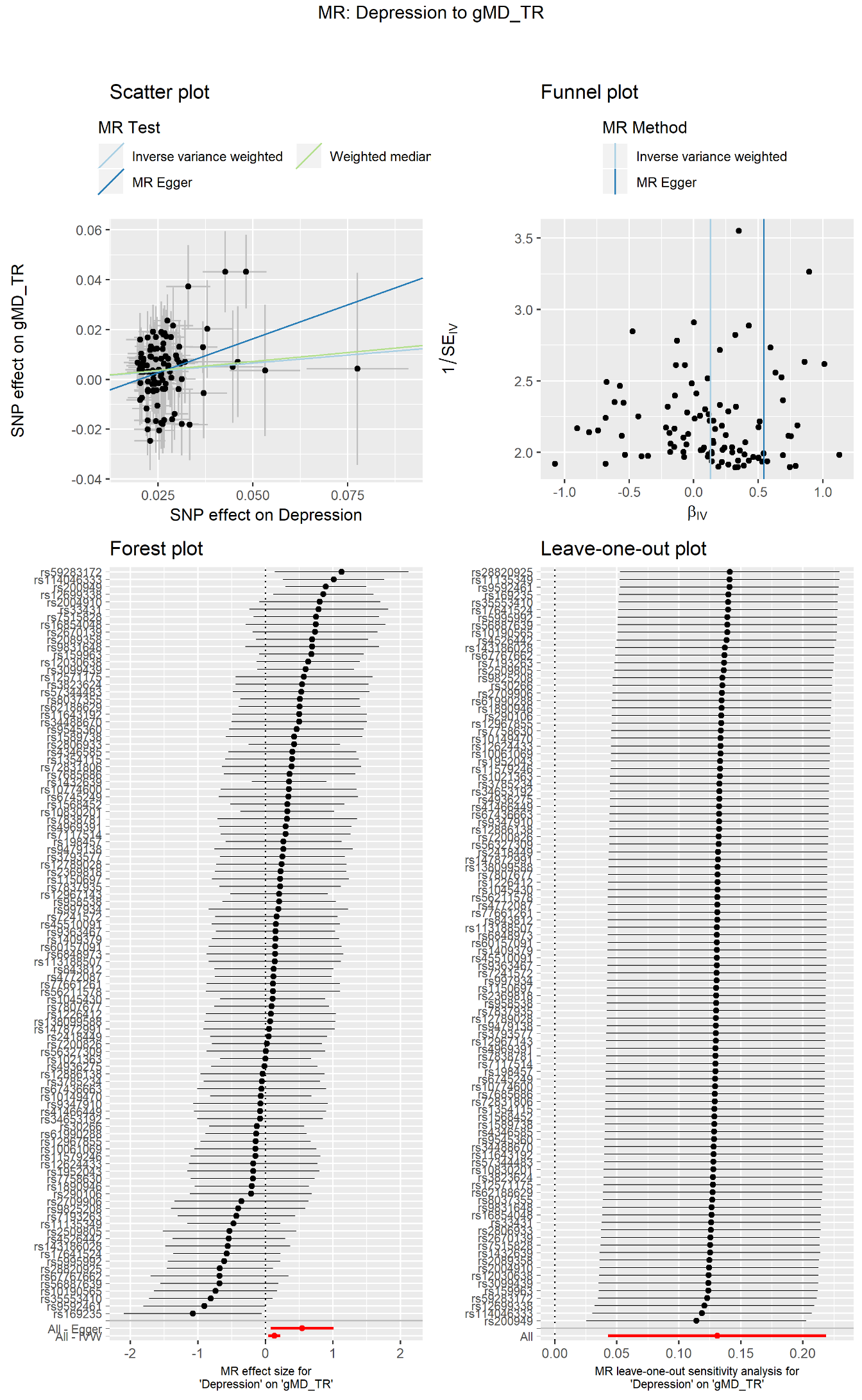

Figure S16. Mendelian Randomisation analysis testing the causal effect of depression on MD in forceps minor.

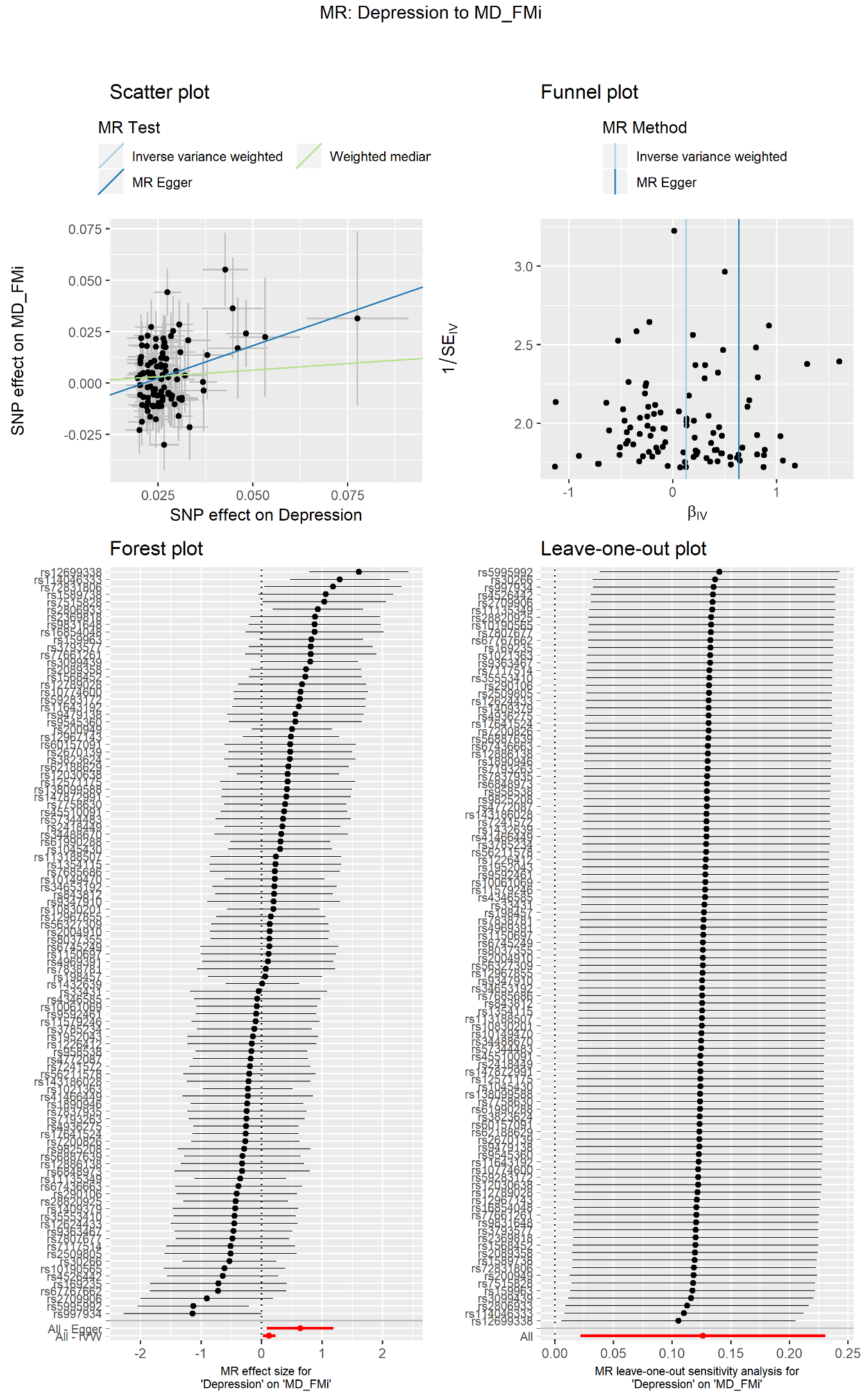

Figure S17. Mendelian Randomisation analysis testing the causal effect of depression on MD in inferior fronto-occipital fasciculus.

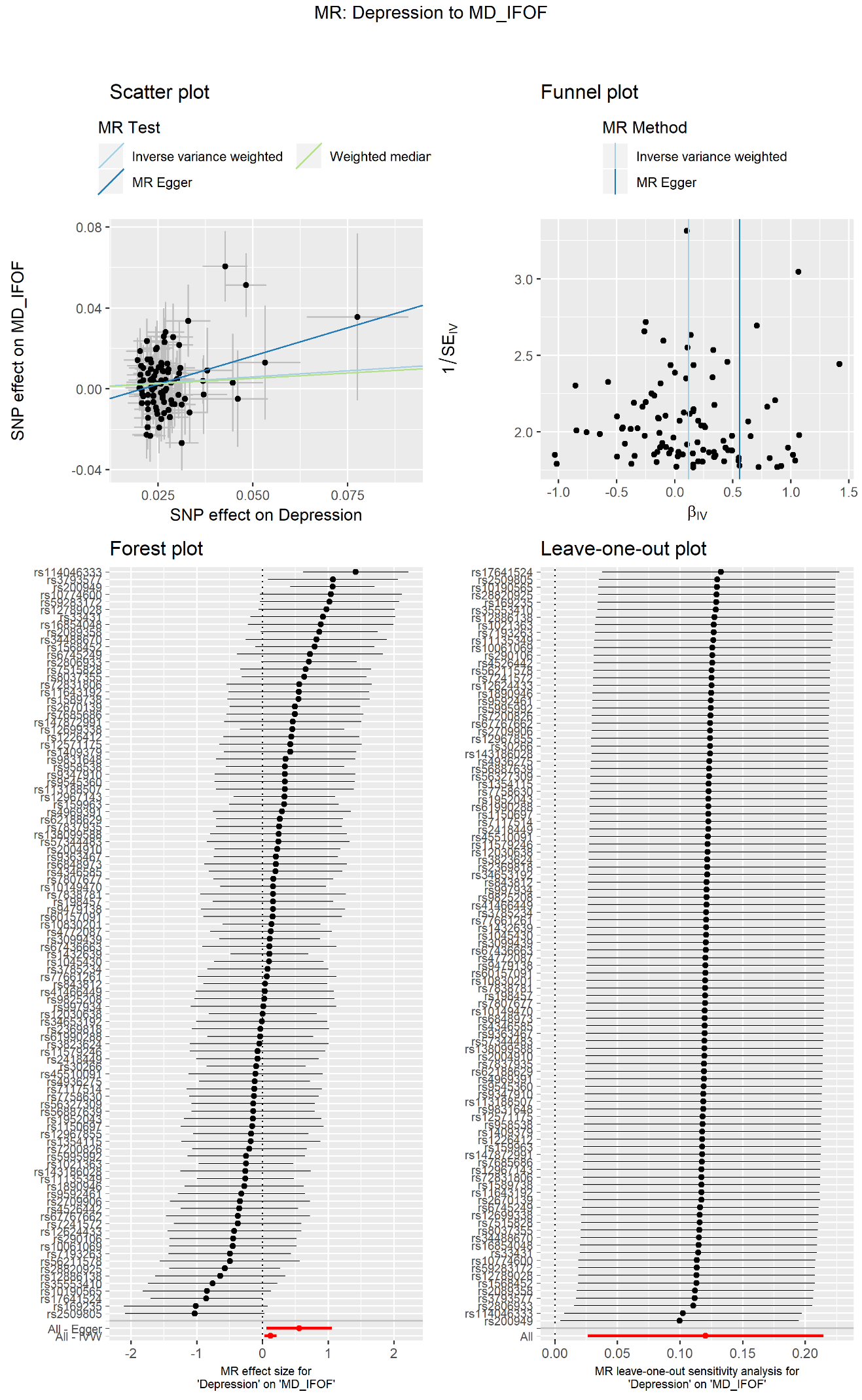

Figure S18. Mendelian Randomisation analysis testing the causal effect of depression on MD in superior longitudinal fasciculus.

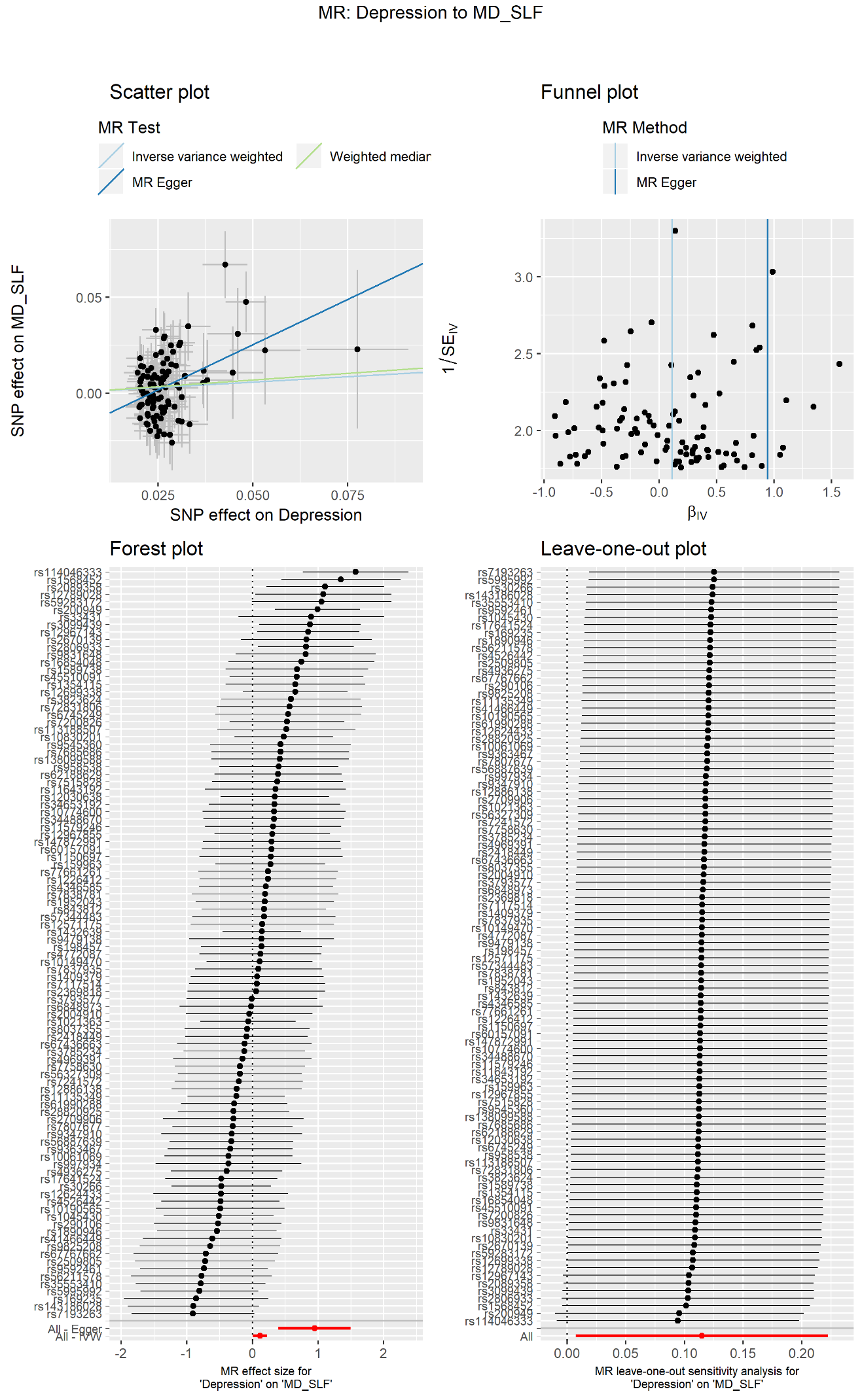

Figure S19. Mendelian Randomisation analysis testing the causal effect of depression on MD in fluctuation amplitude of the Salience Network (Node 14).

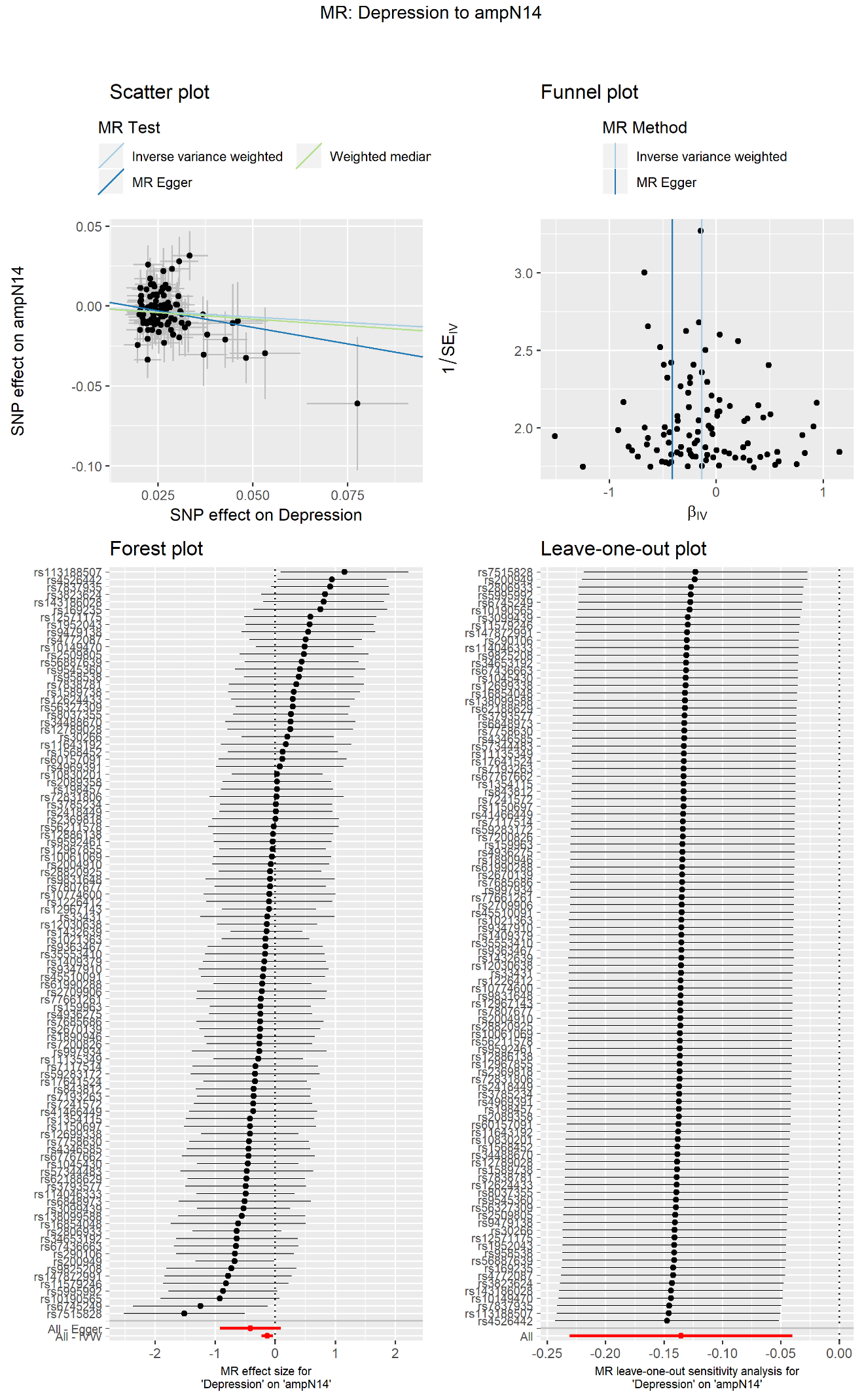

Figure S20. Mendelian Randomisation analysis testing the causal effect of MD in anterior thalamic radiation on depression.

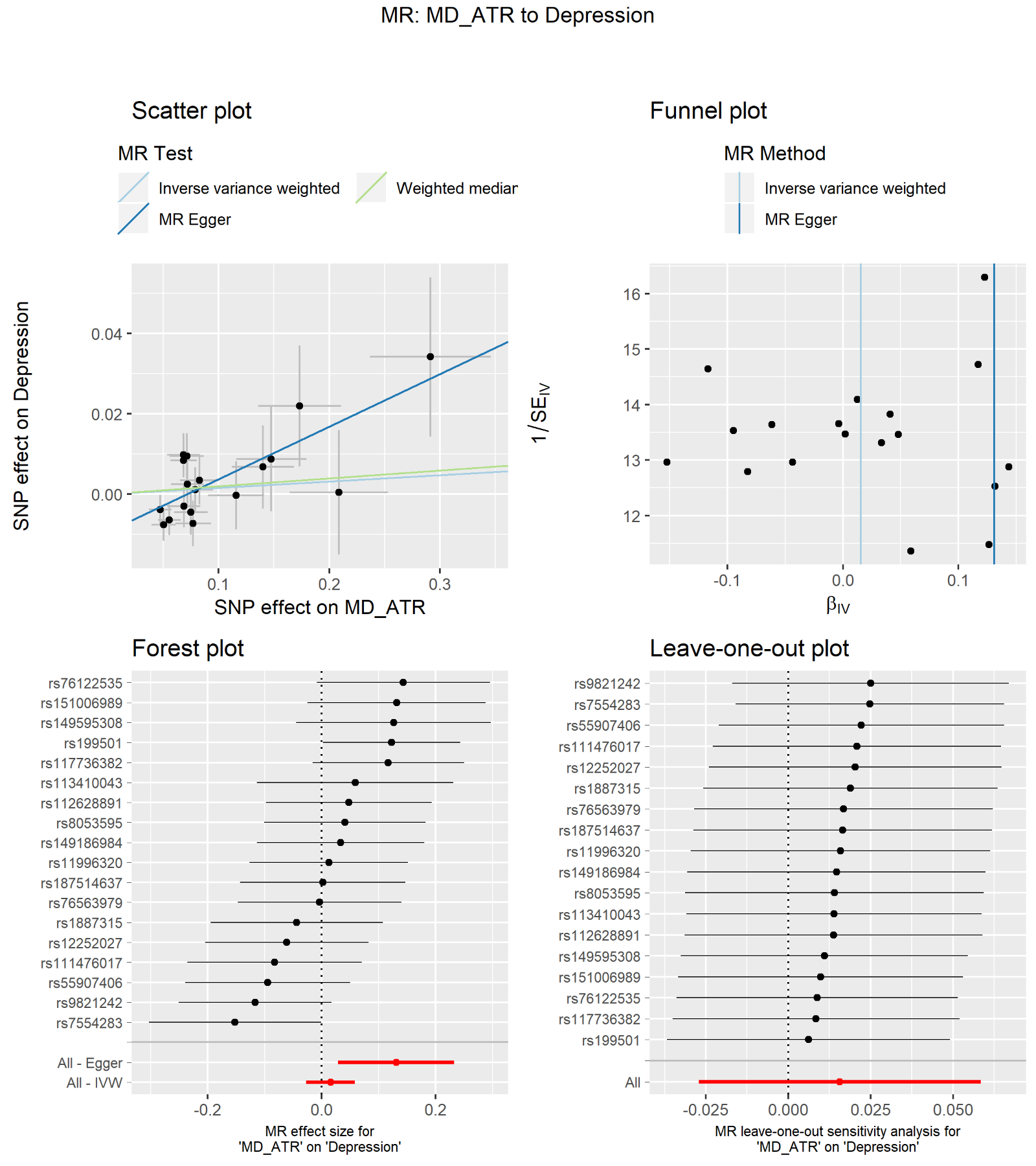

Figure S21. Interaction between depression-PGRS and variables including sociodemographic variables and early-life risk factors for depression. Dependent variables are the traits that showed significant associations with depression-PGRS in at least four thresholds in both the discovery and replication samples. In the panels, each dot represents the p value for a dependent variable. The red lines represent the threshold for nominal significance, whilst the yellow lines represent the threshold for significance after FDR correction.

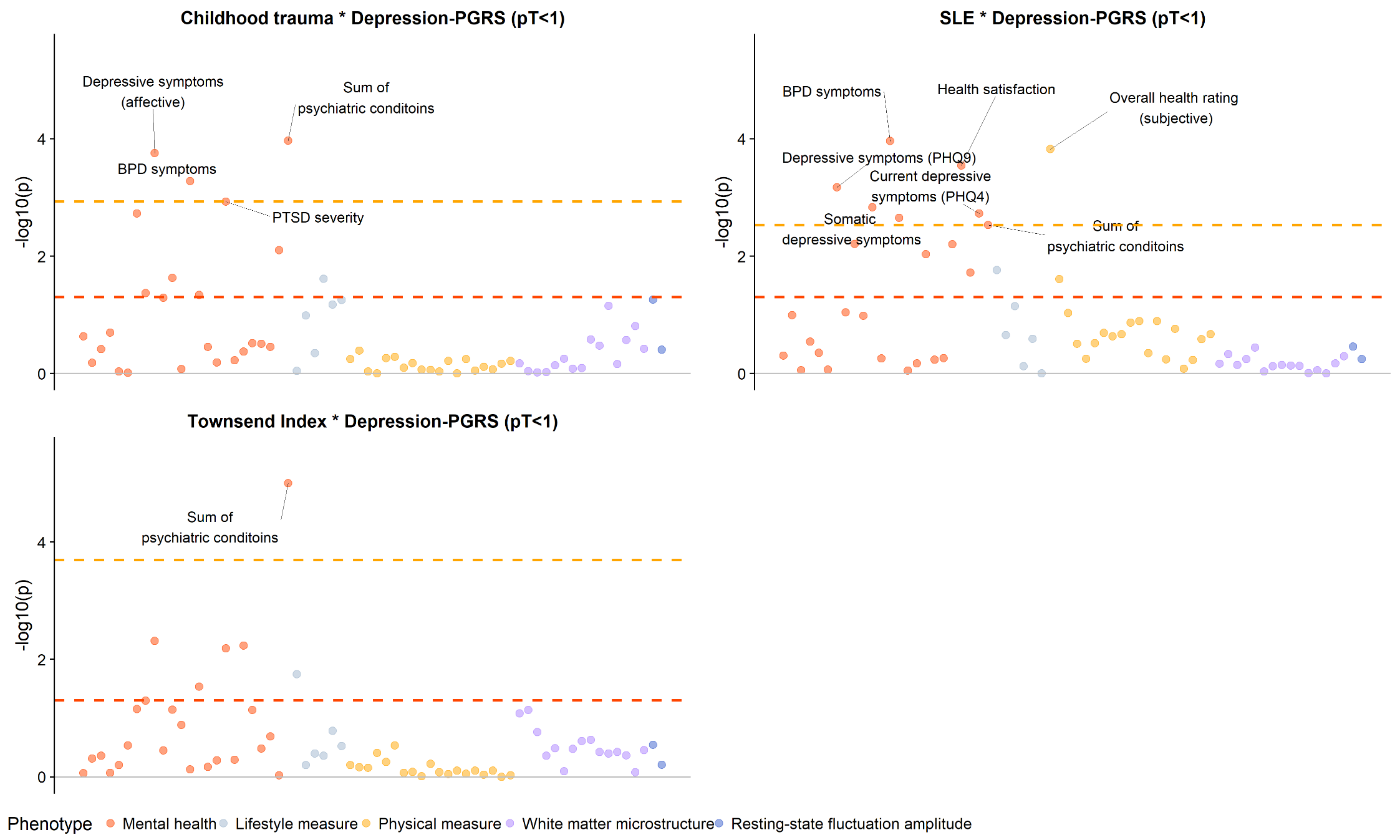

Figure S22. Scree plots of PCA on fractional anisotropy (FA) and mean diffusivity (MD). The x axes represent the principal components. The y axes represent variance explained by each principal component. PCA was conducted on (i) all the tracts (gTotal), (ii) association fibres (gAF), (iii) thalamic radiations (gTR) and (iv) projection fibres (gPF).

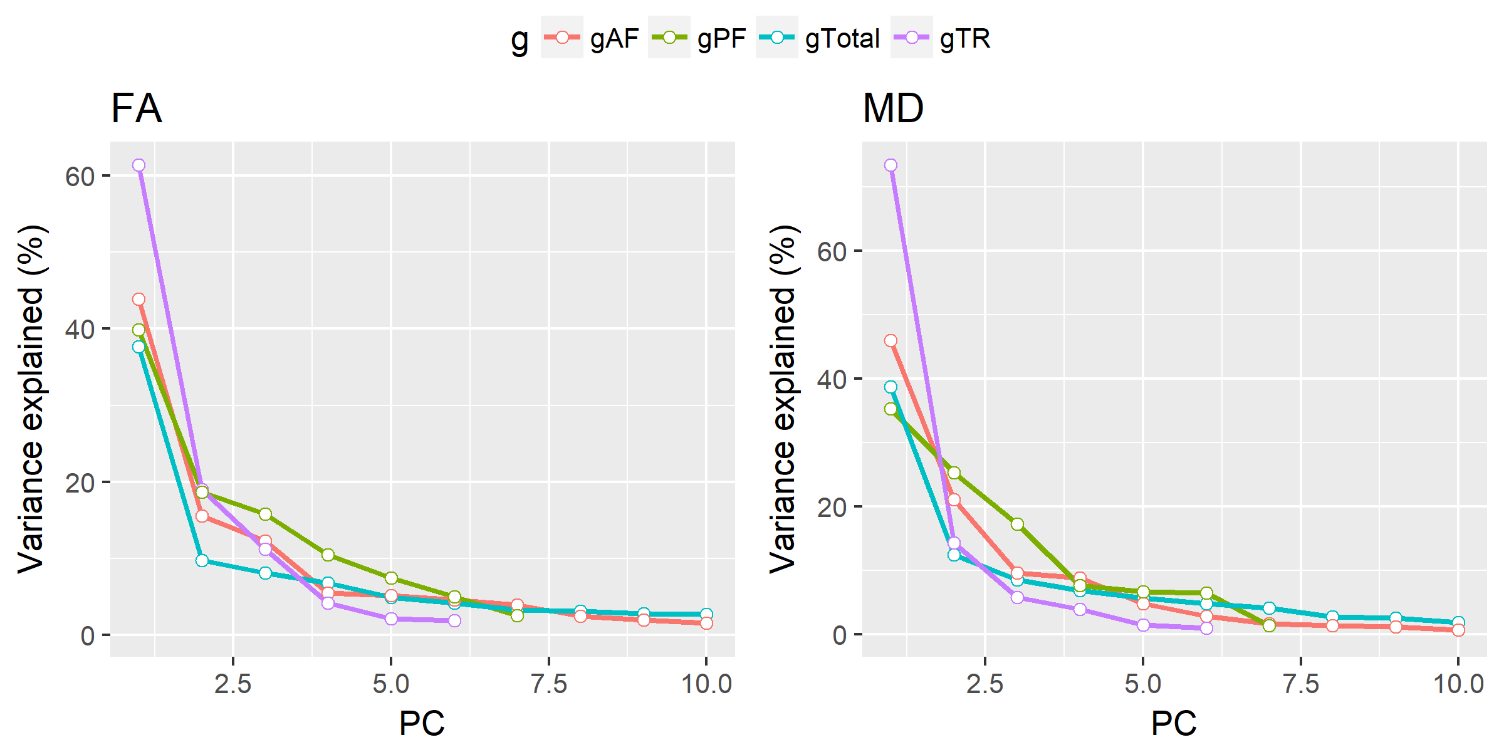

Table S1. Descriptions of phenotypes included in the PheWAS (file name: TableS1.all_phenos.xlsx). In this file, all phenotypes included in the analyses were listed with their names and the according field names in the UK Biobank data showcase. The order of phenotypes was identical to that in table S3-8 where PheWAS results are listed.

Table S2. Summary of GWAS for neuroimaging (file name: TableS2.IM_GWAS_summary.xlsx). The sample sizes are the same with the ones for PheWAS combining both discovery and replication samples, therefore for brevity, they are not included in this table. In the table, estimated partitioned heritability using LDSC, numbers of genome-wide significant hits (>3,000kb, r^2^<0.001, p<5*10^-8^) are reported. Softwares and parameters used for GWAS can be found in the methods section in the main text.

Table S3. Results for Mendelian Randomisation analyses (filename: TableS3.MR_results.xlsx). Results for causal effect of depression to neuroimaging variables are presented in sheet ‘MR.Depression_to_IM’, and results for the reverse analysis are presented in ‘MR.IM_to_Depression’. Uncorrected and corrected p values for the MR analyses are reported, and the p values for tests for robustness were FDR-corrected.

Table S4. Model statistics for SEM models (file name: TableS4.PheWAS_mediation.xlsx). Predictors were depression-PGRS, mediators were neuroimaging traits/manifestations of depression, and dependent variables were manifestations of depression/neuroimaging traits.

Table S5. Results for PheWAS conducted on the discovery sample (file name: TableS5.PheWAS_discovery.xlsx). Where the dependent variable was binary, logistic regression and linear regression models were both conducted and therefore both Beta (regression coefficient for linear regression) and Log odds ratio were reported. Each tab contains the results for each depression-PGRS as the factor.

Table S6. Results for PheWAS conducted on the replication sample (file name: TableS6.PheWAS_replication.xlsx). Where the dependent variable was binary, logistic regression and linear regression models were both conducted and therefore both Beta (regression coefficient for linear regression) and Log odds ratio were reported. Each tab contains the results for each depression-PGRS as the factor. Only traits that were significantly associated with depression-PGRS at a minimum of four p thresholds in the discovery dataset were tested and reported.

Table S7. Results for PheWAS conducted on the dataset collected at Cheadle (file name: TableS7.PheWAS_Cheadle.xlsx).

Table S8. Results for PheWAS conducted on the dataset collected at Newcastle (file name: TableS8.PheWAS_Newcastle.xlsx). Only traits that were significantly associated with depression-PGRS at a minimum of four p thresholds in the dataset collected at Cheadle were tested and reported.

Table S9. Results for ‘depression-PGRS × MRI site’ interaction on the traits (file name: TableS9.PheWAS_site_interaction.xlsx). Only traits that were significantly associated with depression-PGRS at a minimum of four p thresholds in the dataset collected at Cheadle were tested and reported.

Table S10. Results for PheWAS conducted on the total sample (N=21,888) (file name: TableS10.PheWAS_meta.xlsx).

Table S11. List of regions for the resting-state node amplitude that negatively associated with higher Depression-PGRS. The report was generated using the ‘report’ function in the ‘xjview’ package in SPM 12.

| **Coordination of the voxel with the highest intensity in the cluster** | **AAL region** | **Cluster size** | **Highest intensity** |
| --- | --- | --- | --- |
| -10, 54, 2 | Cingulum_Ant_L (aal) | 6,679 | -0.704 |
| -44, -30, 46 | Postcentral_L (aal) | 2,278 | -0.776 |
| 44, -24, 40 | Postcentral_R (aal) | 1,184 | -0.647 |
| -38, -4, 16 | Insula_L (aal) | 811 | -0.646 |
| 28, -54, -22 | Cerebelum_6_R (aal) | 408 | -0.529 |
| 22, -58, -48 | Cerebelum_8_R (aal) | 318 | -0.679 |
| 30, 18, -16 | Insula_R (aal) | 300 | -0.402 |
| -2, -18, 10 | Thalamus_L (aal) | 216 | -0.403 |
| 50, -16, 18 | Rolandic_Oper_R (aal) | 211 | -0.380 |
| 32, 36, -10 | Frontal_Inf_Orb_R (aal) | 177 | -0.447 |
| -24, -52, -20 | Cerebelum_6_L (aal) | 177 | -0.446 |
| -34, 34, -12 | Frontal_Inf_Orb_L (aal) | 165 | -0.425 |
| -24, -54, -50 | Cerebelum_8_L (aal) | 120 | -0.474 |
| 40, -2, 16 | Insula_R (aal) | 116 | -0.510 |
| -18, 34, 40 | Frontal_Sup_L (aal) | 111 | -0.389 |
| 2, -64, -18 | Vermis_6 (aal) | 47 | -0.375 |

Table S12. Results for gene by environment (G×E) interaction, testing the interaction between depression-PGRS and traumatic events in adulthood (file name: TableS12.DepressionPGRS_X_Adult.Trauma.xlsx).

Table S13. Results for gene by environment (G×E) interaction, testing the interaction between depression-PGRS and traumatic events in childhood (file name: TableS13.DepressionPGRS_X_Childhood.Trauma.xlsx).

Table S14. Results for gene by environment (G×E) interaction, testing the interaction between depression-PGRS and current stressful life events (file name: TableS14.DepressionPGRS_X_Current.SLE.xlsx).

Table S15. Results for gene by environment (G×E) interaction, testing the interaction between depression-PGRS and household income (file name: TableS15.DepressionPGRS_X_Household.Income.xlsx).

Table S16. Results for gene by environment (G×E) interaction, testing the interaction between depression-PGRS and Townsend Index tertiles (file name: TableS16.DepressionPGRS_X_TDI.xlsx).

Table S17. Loadings of each tract on gTotal, gAF, gTR and gPF in fractional anisotropy (FA) and mean diffusivity (MD) (see Figure S13). The principal component analysis was conducted on the maximum sample size including both discovery and replication samples.

| **Tract** | **FA** | | | | **MD** | | | |
| --- | --- | --- | --- | --- | --- | --- | --- | --- |
|  | **g.Total** | **gAF** | **gTR** | **gPF** | **g.Total** | **gAF** | **gTR** | **gPF** |
| Parahippocampal part of cingulum | 0.361 | 0.379 | -- | -- | 0.472 | 0.61 | -- | -- |
| Parahippocampal part of cingulum | 0.394 | 0.421 | -- | -- | 0.455 | 0.588 | -- | -- |
| Forceps major | 0.547 | 0.552 | -- | -- | 0.488 | 0.518 | -- | -- |
| Cingulate gyrus part of cingulum | 0.558 | 0.635 | -- | -- | 0.609 | 0.576 | -- | -- |
| Cingulate gyrus part of cingulum | 0.582 | 0.658 | -- | -- | 0.614 | 0.59 | -- | -- |
| Uncinate fasciculus | 0.649 | 0.666 | -- | -- | 0.665 | 0.7 | -- | -- |
| Uncinate fasciculus | 0.682 | 0.697 | -- | -- | 0.734 | 0.75 | -- | -- |
| Superior longitudinal fasciculus | 0.81 | 0.796 | -- | -- | 0.859 | 0.795 | -- | -- |
| Superior longitudinal fasciculus | 0.824 | 0.8 | -- | -- | 0.847 | 0.777 | -- | -- |
| Forceps minor | 0.803 | 0.804 | -- | -- | 0.716 | 0.655 | -- | -- |
| Inferior longitudinal fasciculus | 0.822 | 0.814 | -- | -- | 0.862 | 0.837 | -- | -- |
| Inferior longitudinal fasciculus | 0.85 | 0.827 | -- | -- | 0.893 | 0.85 | -- | -- |
| Inferior fronto-occipital fasciculus | 0.827 | 0.83 | -- | -- | 0.871 | 0.836 | -- | -- |
| Inferior fronto-occipital fasciculus | 0.853 | 0.842 | -- | -- | 0.89 | 0.849 | -- | -- |
| Superior thalamic radiation | 0.654 | -- | 0.73 | -- | 0.773 | -- | 0.844 | -- |
| Superior thalamic radiation | 0.65 | -- | 0.736 | -- | 0.743 | -- | 0.82 | -- |
| Posterior thalamic radiation | 0.676 | -- | 0.785 | -- | 0.756 | -- | 0.863 | -- |
| Posterior thalamic radiation | 0.775 | -- | 0.808 | -- | 0.82 | -- | 0.851 | -- |
| Anterior thalamic radiation | 0.693 | -- | 0.813 | -- | 0.775 | -- | 0.88 | -- |
| Anterior thalamic radiation | 0.773 | -- | 0.817 | -- | 0.811 | -- | 0.847 | -- |
| Medial lemniscus | 0.262 | -- | -- | 0.501 | 0.253 | -- | -- | 0.329 |
| Middle cerebellar peduncle | 0.274 | -- | -- | 0.52 | 0.223 | -- | -- | 0.31 |
| Medial lemniscus | 0.365 | -- | -- | 0.539 | 0.351 | -- | -- | 0.933 |
| Acoustic radiation | 0.608 | -- | -- | 0.577 | 0.486 | -- | -- | 0.307 |
| Acoustic radiation | 0.617 | -- | -- | 0.631 | 0.504 | -- | -- | 0.275 |
| Corticospinal tract | 0.576 | -- | -- | 0.802 | 0.579 | -- | -- | 0.283 |
| Corticospinal tract | 0.581 | -- | -- | 0.821 | 0.58 | -- | -- | 0.296 |
